# Plasmatic immune extracellular vesicle profiles identify prodromal and early stages of Parkinson’s disease

**DOI:** 10.64898/2026.05.12.724498

**Authors:** Elena Vacchi, Jacopo Burrello, Alessio Burrello, Sara Bolis, Inigo Ruiz-Barrio, Ilaria Bertaina, Luca Baldelli, Maria Giulia Bacalini, Giacomo Chiaro, Rudolf Kaelin, Ankush Yadav, Sandra Pinton, Alberto Romagnolo, Simona Vittoria Maule, Sandra Hackethal, Silvia Riccardi, Silvia Miano, Giovanni Bianco, Claudio Staedler, Javier Pagonabarraga, Jaime Kulisevsky, Federica Provini, Georg Kägi, Mauro Manconi, Salvatore Galati, Alain Kaelin-Lang, Lucio Barile, Giorgia Melli

## Abstract

Extracellular vesicles (EVs) hold promise as minimally invasive biomarkers for neurodegenerative proteinopathies, but disease- and stage-specific profiles remain unclear.

For this study, we enrolled 378 participants across five centers and the MJFF-BioFIND cohort: 100 healthy controls [HC], 64 isolated REM sleep behavior disorder [iRBD], 41 DeNovo Parkinson’s Disease [PD], 89 Late PD, 32 other Synucleinopathies, and 52 Tauopathies. All participants underwent clinical evaluation and blood collection. The 77 subjects from the BioFIND cohort also provided CSF samples. EV concentration and size were assessed by nanoparticle tracking analysis; flow cytometry quantified tetraspanins (CD9/CD63/CD81) and 37 surface markers. Multivariable logistic regression, receiver operating characteristic analyses (ROC), and repeated random forest (rRF) classifiers evaluated diagnostic utility.

Late PD showed the highest EV concentrations compared to HC and other disease groups. Participants exhibited distinct EV surface immunophenotypes, with the iRBD group displaying the most extensive immune activation signature vs HC, followed by PD patients. Multivariate logistic regression analysis identified diagnostic marker panels: CD3/CD9/CD25/CD56 for iRBD, SSEA4 for Late PD, CD146/CD209 for Synucleinopathies, and CD8/CD45/CD62P for Tauopathies. ROC confirmed good discriminatory performance, with CD56 emerging as the strongest single predictor for iRBD vs HC, SSEA4 showing high sensitivity for Late PD, and marker combinations providing optimal balance for Synucleinopathy/Tauopathy classification vs HC. In the CSF BioFIND subset, Late PD EVs exhibited increased myeloid (CD1c), adhesion (CD29), activation (CD69), and epithelial (CD326) markers compared to HC. Among these, CD326 was independently associated with Late PD diagnosis. Machine learning classifiers using all 37 surface antigens achieved excellent training performance (91.7-94.3% accuracy for iRBD/Synucleinopathies vs HC) and maintained robust validation accuracy, particularly for iRBD (77.8%) and DeNovo PD (76.6%) vs HC.

EV immuno-phenotyping reveals distinct signatures across the neurodegenerative proteinopathies spectrum, with the highest diagnostic utility for prodromal iRBD detection. Longitudinal validation and cell-of-origin refinement represent key next steps toward clinical translation.

## Introduction

Parkinson’s disease (PD), the second most common neurodegenerative disorder, lacks definitive diagnostic tools, posing a growing challenge for healthcare systems worldwide^1,2^. Noninvasive biomarker-based tests are urgently needed for early diagnosis, differentiation from atypical parkinsonisms, clinical trial stratification, and disease-modifying therapy development.

PD is biologically characterized by progressive dopaminergic neurodegeneration, accumulation of pathological α-synuclein in the brain, peripheral tissues, and CSF, and genetic variants that increase disease susceptibility^3,4^. Many studies have leveraged these biological hallmarks to enhance diagnostic accuracy, and several promising strategies are currently under investigation^5,6^.

Neuroinflammation has recently emerged as another important biological key aspect of the disease. Indeed, PD exhibits a strong inflammatory component that involves both the central nervous system (CNS) and the peripheral immune system from the early stages of the disease^7,8^. Postmortem and *in vivo* data, confirmed by PET imaging^9^ and single-cell RNA sequencing^10^, reveal microglia activation from prodromal stages^11^ (e.g., isolated REM sleep behavior disorder, iRBD), through advanced PD^12^. Both brain and CSF of PD showed elevated cytokines and chemokines (TGF-β1, IL-6, and IL-1β)^7^, as well as infiltration of monocytes^13^ and T cells^14,15^; while peripheral blood exhibits CD4+ and CD8+ T cells depletion^16^ with activated-to-memory phenotypes shifting, regulatory T cells (Tregs) dysfunction^17,18^, pro-inflammatory Th1/Th17 expansion^17,18^, stage-dependent monocyte activation^19^ correlating with severity and dementia risk^19,20^, and altered neutrophil-to-lymphocyte ratios variably linked to disease duration^21^ and motor worsening^22^.

Extracellular Vesicles (EVs) are key mediators in immune functions, both in physiological and pathological conditions, and can be exploited for biomarker and therapeutic purposes^23,24^. With their cargo of DNA, RNA, and proteins, EVs can influence immune cell communication, modulate immune cell phenotypes and functions, and regulate inflammation, antigen presentation, and immune cell responses^23,24^. However, their role in PD immune response and inflammation has not yet been fully explored.

In our previous study^25^, we identified distinct subsets of plasma-derived EV, characterized by differential expression of surface markers related to inflammatory and immune cells. A diagnostic model based on these EV profiles stratified patients with PD and atypical parkinsonism (AP) according to their clinical diagnoses, suggesting that distinct inflammatory pathways may underlie different neurodegenerative conditions. As this pilot study primarily involved PD patients with long disease duration, we expanded our cohort to include prodromal iRBD patients and individuals at early disease stages, hypothesizing that EV surface marker profiling can serve as an early diagnostic tool in PD, reflecting specific inflammatory signatures active at the onset of the pathogenic process.

## Methods

### Study cohort

Participants of the study were consecutively recruited from the Movement Disorders and Sleep Medicine Clinic at Neurocenter of Southern Switzerland (NSI-EOC, Lugano, Switzerland), the Movement Disorders Clinic at St. Gallen Hospital (Switzerland), the Sleep Medicine Clinic at the University Hospital of Bologna (Italy), and the University Hospital of Torino (Italy), as part of the multicentric, prospective, case-control NSIPD001 study, and from the Movement Disorders Unit at Hospital de la Santa Creu i Sant Pau, Barcelona (Spain), as part of the IIBSP-PAR-2022-98. All patients provided informed consent in accordance with the principles outlined in the Declaration of Helsinki.

Inclusion criteria were based on published diagnostic criteria: PD^26^, Progressive supranuclear palsy^27^ (PSP), Corticobasal Degeneration (CBD), Multiple System Atrophy^28^ (MSA), Lewy Body Dementia (LBD)^29^. PD group included DeNovo patients, defined as subjects with criteria for a definite diagnosis of PD and less than 3 years since the diagnosis, and Late PD with > 3 years from the diagnosis. Inclusion criteria for iRBD were based on standard diagnostic criteria and a video-polysomnography documentation of REM sleep without atonia^30^. Healthy controls (HC), without any known neurological disease, were recruited among hospital staff and patients’ partners. Exclusion criteria for all participants were the presence of major cognitive impairment or major dysautonomic symptoms in the family history, and significant comorbidities such as diabetes, renal failure, thyroid pathology, HIV infection, syphilis, coagulopathy, fever, acute or chronic inflammatory diseases, and tumors.

### Clinical assessments

Disease subjects underwent a standard clinical evaluation. Disease severity was measured using the Hoehn and Yahr^31^ scale and the Movement Disorder Society-Unified Parkinson’s Disease Rating Scale^32^ (MDS-UPDRS; scale I—clinical evaluation of mental state, behavior, and mood; scale II—self-assessment by the patient of specific daily activities; scale III—clinical evaluation of motor skills). Cognitive impairment was assessed by the Mini-Mental State Examination^33^ (MMSE) and the Montreal Cognitive Assessment (MoCA) scales^34^. Autonomic dysfunction was analysed by the Composite Autonomic Symptom Score 31^35^ (COMPASS-31, OH: orthostatic hypotension, VM: vasomotor, SM: sudomotor, GI: gastrointestinal, BL: bladder, PM: pupillomotor). Mood disorder was assessed by the Beck Depression Inventory-II (BDI-II) scale^36^. Rapid eye movement sleep Behavior Disorder (RBD) was measured by the RBD screening questionnaire^37^. Finally, Levodopa equivalent daily dose^38^ (LEDD) was calculated.

### Blood Collection

All patients underwent blood collection. Ten milliliters of blood were collected into anticoagulant ethylenediamine tetraacetic acid (EDTA) tubes in the morning after a 4-hour fast. Blood was centrifuged to deplete cellular components (15 minutes at 1,600g at 10°C). The obtained plasma was aliquoted and stored at ™80°C until use. On the day of the experiment, after thawing, plasma was centrifuged three times to deplete platelets and cellular debris (15 minutes at 3,000g at 4°C), apoptotic bodies (15 minutes at 10,000g at 4°C), and larger EVs (30 minutes at 20,000g at 4°C), and then tested with Nanoparticle Tracking Analysis (NTA) and Flow Cytometry.

### BioFIND cohort

The Michael J. Fox Foundation for Parkinson’s Research (MJFF) provided demographic data (age, sex, BMI), clinical information (H&Y, MDS-UPDRS, BDI-II, MoCA, RBDSQ), plasma, and CSF samples of 77 subjects (26 HC and 51 Late PD) from the BioFIND cohort, a multicentric cohort for biomarker discovery, including participants with moderate to advanced PD and age- and sex-matched healthy controls^39^.

On the day of the experiment, after thawing, plasma and CSF samples underwent the same preparation protocols as the other samples and were analyzed.

### Nanoparticle Tracking Analysis (NTA)

Nanoparticle concentration (N° Nanoparticles/mL) and diameter (nm) were measured using the ZetaView instrument (Particle Metrix, Germany), which employed an embedded laser (40 mW at 520 nm) and a CMOS camera. For the analysis, plasma was diluted 1:1000, while CSF 1:500 in PBS1X. The manufacturer’s default software settings for EVs were selected. For each measurement, videos of 11 cell positions were acquired at 25°C. After capture, videos were analyzed by the inbuilt ZetaView Software 8.02.31. The number of completed tracks in NTA measurements was always greater than the proposed minimum of 1000, which helps minimize data skewing due to a single large particle.

### Flow Cytometry

The MACSPlex Human Exosome Kit (Miltenyi, Bergisch Gladbach, Germany) was used to investigate the expression of multiple EV surface antigens. As previously reported^25,40^, 60μL of plasma were incubated with 60μL of Miltenyi buffer solution and 10μL of capture beads overnight in a shaking thermomixer at 10°C and 800rpm. For CSF analysis, 10μL of capture beads were added to 200μL of CSF before overnight incubation. The day after, the beads were pelleted with a 3000g centrifuge for 15’ at 10°C and then incubated for 1h at 10°C, 800rpm with 10μL of allophycocyanin (APC)-detection antibodies against CD9, CD63, and CD81.

Samples were washed and analyzed with the MACSQuant Analyzer-10 flow cytometer (Miltenyi). Triggers for side and forward scatters were selected to confine the measurement to the multiplex beads. PE and FITC lasers were used to visualize the subsets of capture beads. For each population, the APC signal was measured. A blank control composed only of MACSPlex Buffer and incubated with capture beads and detection antibodies was used to measure the background signal.

Analysis was performed according to the manufacturer’s protocol. The median fluorescence intensity (MFI) of each EV marker was calculated by subtracting the blank control to remove the background signal; negative values were set to zero. The MFI of each EV marker was normalized to the mean MFI for specific EV markers (CD9, CD63, and CD81). The normalized mean fluorescent intensity (nMFI) was used for all the subsequent analyses.

## Statistical analyses

Statistical analyses were performed with IBM SPSS Statistics 26.0 and GraphPad PRISM 9.0.

Normally distributed variables (were expressed as mean±SD and analyzed by the 1-way analysis of variance test with the post hoc Bonferroni test for multiple comparisons. Non-normally distributed variables were expressed as medians and interquartile ranges, and analyzed using the Kruskal-Wallis test. Categorical variables were expressed as absolute numbers and percentages (%) and analyzed by χ2 or Fisher’s exact tests. Multivariate logistic regression analysis was performed to assess the Odds Ratio (OR). Receiver operating characteristic (ROC) curve analysis was used to evaluate the area under the curve (AUC) and to compare the diagnostic performances of selected variables. The Youden index (J = Sensitivity + Specificity − 1) was calculated to determine the cutoff with the greatest accuracy. Correlations were assessed using Pearson correlation test after Z-score normalization. Correlations were considered strong for coefficient between |1.0| and |0.6|, moderate between |0.6| and |0.3|, and weak between |0.3| and |0.1|. Python 3.11 was used to draft the correlation matrix. P-values lower than 0.05 were considered significant. Missing data were handled via complete case analysis; patients lacking data for a given parameter were excluded from that specific analysis. No imputation methods were applied.

### Diagnostic modeling and validation

Supervised machine learning (ML) classifiers, and in particular random forest regressors (rRF), were applied to develop diagnostic models in the training cohort, which were then tested in the validation cohort. The overall cohort was randomly split into the training (n=253) and validation (n=125) cohorts according to a ratio of 70:30. rRF was applied to formulate predictions on the primary endpoint (patient diagnosis) on the base of a set of labeled multi-dimensional paired input-output data; input data were represented by EV profiling, with ML algorithms used to combine data of expression of the 37 evaluated surface antigens in a biomolecular signature.

The rRF classifiers create a pre-defined set of classification trees (“n” classification trees) with a fixed maximum number of splits for each tree. The predicted endpoint results from the outcome of each classification tree of the forest (healthy control versus disease condition); if at least (n/2)+1 of “n” trees of the RF predict a particular diagnosis, then this endpoint is assigned to the patient.

In order to correct for dataset imbalance, three different oversampling algorithms were applied to the rRF classifier: synthetic minority over-sampling technique (SMOTE), SMOTE and nearest neighbors (SMOTENN), and random oversampling (RO). Oversampling algorithms impute data of new simulated subjects starting from those of real subjects of our overall cohort, in order to balance the number of patients with the different disease conditions as compared to healthy subjects, at model training. These algorithms are applied to limit a falsely high accuracy due to over-prediction of the most represented class by the rRF (accuracy paradox).

A grid search was performed to explore the diagnostic performance of the 576 rRF trained and validated models, with and without correction by oversampling algorithms. The two hyperparameters to be tuned in the rRF algorithm were the maximum number of splits (leaves; from 10 to 320) and the number of classification trees in the forest (from 10 to 200). Analyses involving the application of machine learning techniques were performed using Python 3.5 (library, scikit-learn).

## Results

### Study cohort description

For this study, a total of 378 patients from five hospital research centers and the BioFIND cohort of MJFF were included (Supplementary Table 1): 100 HC, 64 iRBD, 41 DeNovo PD, 89 Late PD, 32 Synucleinopathies, and 52 primary Tauopathies. Among Synucleinopathies, 25 were MSA and 7 LBD, while among Tauopathies, 43 were PSP and 9 CBD. Demographic data, medical history, and clinical assessments for each group are summarized in Supplementary Tables 2 and 3.

Tauopathies were significantly older than HC, iRBD, DeNovo, and Late PD, while no significant age differences were observed among other groups. As expected based on their epidemiology, iRBD were predominantly male; thus, sex differences were detected between iRBD and all other groups. Additionally, in HC, a higher percentage of females was present compared to Late PD. No differences in BMI (a possible confounding factor for EVs) were observed between categories.

Among patients, Late PD had the longest disease duration, while Tauopathies showed a later age of onset than all the other disease groups, except for Synucleinopathies (Figure 1A-B). A higher degree of motor impairment was observed in Late PD, Synucleinopathy, and Tauopathy, as assessed by the H&Y and MDS-UPDRS-III scales (Figure 1C-D). In contrast, the MMSE and MoCA scales highlighted more compromised cognitive functions in subjects with Synucleinopathy and Tauopathy (Figure 1E-F).

**Figure 1.**
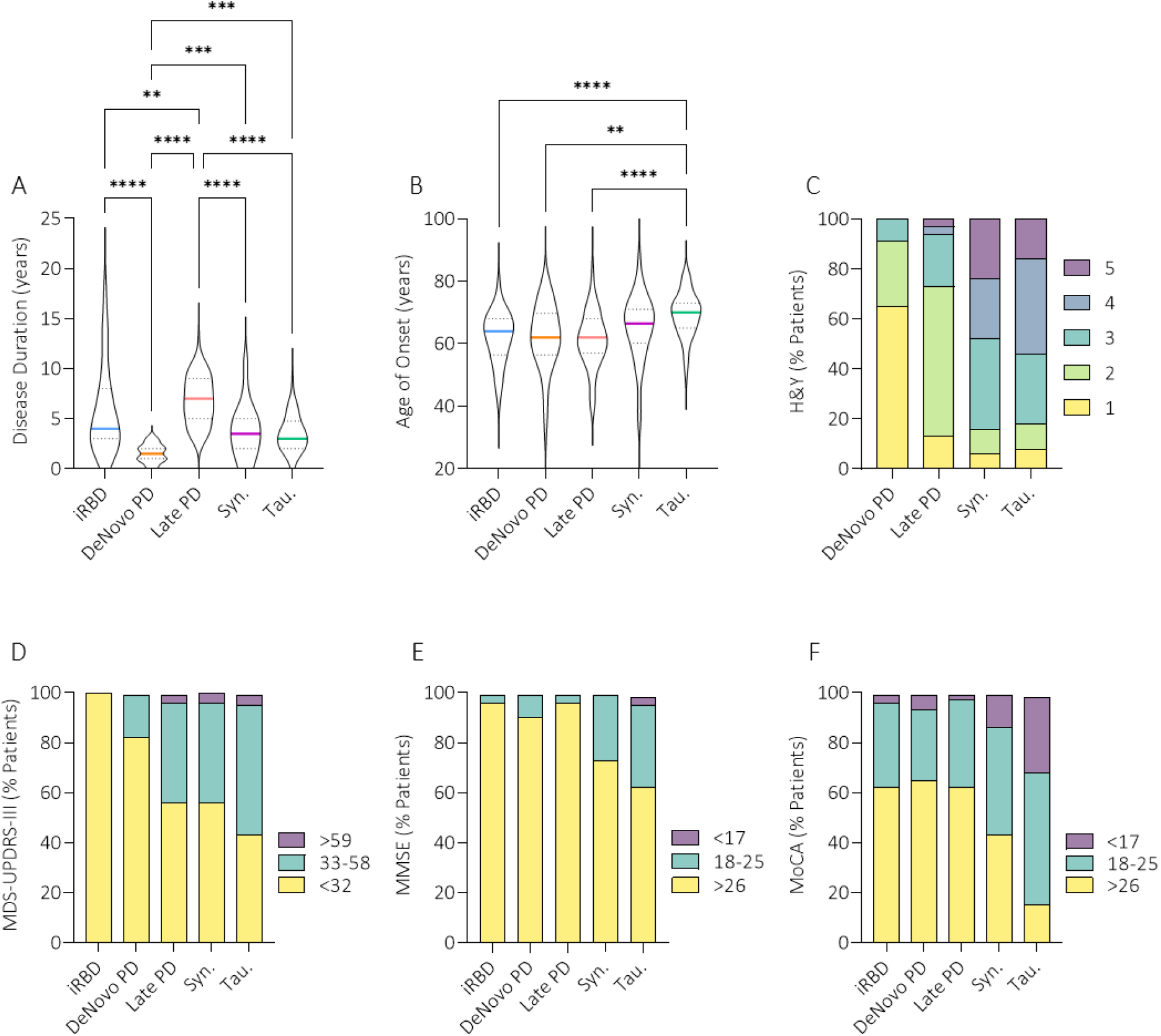
Disease patients’ clinical assessment. Violin plots showing the median (colour line) and interquartile range (dashed line) of disease duration (A) and age of onset (B) across the different disease groups. P-values <0.05 were considered significant: *p<0.05, **p<0.01, ***p<0.001, ****p<0.000. Bar plots showing the distribution of H&Y stages (C), MDS-UPDRS part III scores (D), MMSE (E), and MoCA scores (F) expressed as a percentage of patients per group.

### Late PD show highest EV concentration

NTA analysis did not reveal between-group differences in particle concentration and size (Supplementary Tables 4 and 5). Because NTA cannot discriminate EVs from other small particles, the mean MFI of CD9, CD63, and CD81 obtained by flow cytometry was used as a measure of EV concentration, revealing a significantly higher EV abundance in Late PD compared with HC and other disease groups, and reduced EV expression in DeNovo PD and Tauopathies compared with HC (Figure 2A).

**Figure 2.**
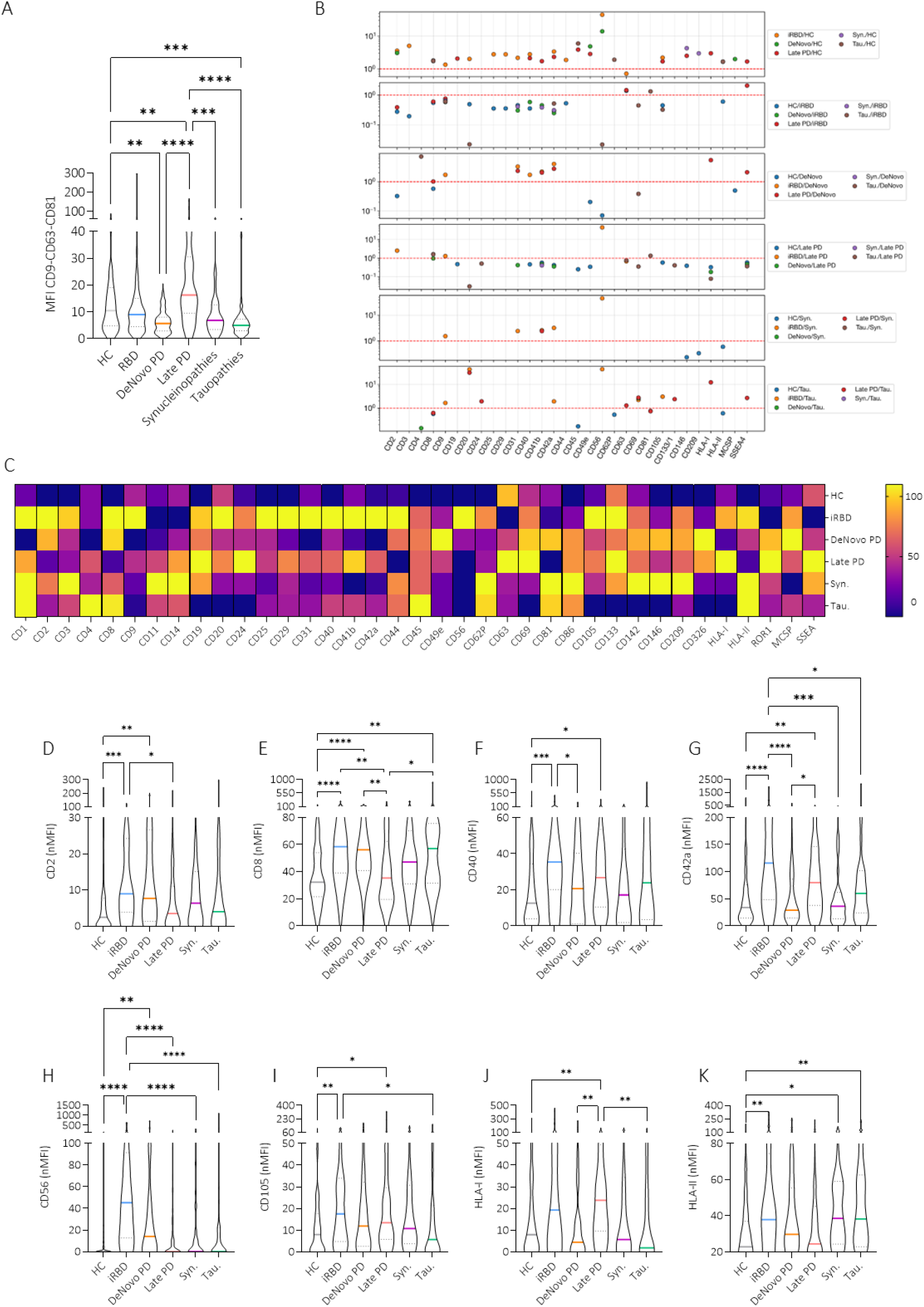
Plasma EV surface markers expression. (A) Violin plots showing the median (colour line) and interquartile range (dashed line) of MFI of CD9-CD63-CD81 across the different groups. P-values <0.05 were considered significant: *p<0.05, **p<0.01, ***p<0.001, ****p<0.000. (B) Multi-panel dot plots depicting the fold-change expression (y-axis, log10 scale) of individual EV-surface markers (x-axis), differently expressed between groups. Red dashed lines indicate the reference threshold (no difference between groups); values above or below this line denote relative up- or down-regulation, respectively. (C) Heatmap representation of the 37 EV-surface markers analysed across the different groups, with colours indicating normalized expression levels from low (dark) to high (yellow). (D–K) Violin plots displaying the MFI of selected EV-surface markers (CD2, CD8, CD40, CD42a, CD56, CD105, HLA-I, and HLA-II) across the different groups, with coloured lines indicating the median and dashed lines the interquartile range; asterisks denote statistically significant differences as in panel A

### iRBD display broadest immune activation profile

Disease patients exhibited differential expression of EV surface markers compared with HC (Figure 2B-K, Supplementary Figure 1, Supplementary Tables 4 and 5). Specifically, among the 37 analyzed surface markers, iRBD showed increased expression of CD2, CD3, CD8, CD9, CD20, CD25, CD29, CD31, CD40, CD42a, CD44, CD56, CD105, and HLA-II, whereas CD63 was decreased. CD2, CD8, and CD56 were also overexpressed in DeNovo PD vs HC, along with CD49e and MCSP. Late PD vs HC showed overexpression of CD40, CD42a, and CD105, similar to iRBD patients, as well as CD19, CD41b, CD45, CD146, HLA-I, and SSEA4. Synucleinopathies showed increased expression of CD146, CD209, and HLA-II vs HC, whereas Tauopathies of CD8, CD45, CD62P, and HLA-II.

iRBD patients showed increased expression of various surface markers *vs* other disease groups (Figure 2B-K). In particular: CD9, CD31, CD40, CD41b, CD42a *vs* DeNovo PD; CD2, CD8, CD9, CD56 *vs* Late PD; CD9, CD31, CD41b, CD42a, CD56 *vs* Synucleinopathies; CD9, CD20, CD42a, CD56, CD69 and CD105 *vs* Tauopathies. CD63 and SSEA-4 were decreased in iRBD *vs* Late PD, and CD81 *vs* Tauopathies. DeNovo PD showed a decrease in CD8, CD31, CD41b, CD42a, HLA-I, and SSEA-4 *vs* Late PD, while *vs* Tauopathies, they showed a decrease in CD4 and an increase in CD69. Late PD had higher CD41b *vs* Synucleinopathies; while, *vs* Tauopathies, CD20, CD24, CD63, CD69, CD133/1, HLA-I, and SSEA4 were increase, CD8 and CD81 were decreased. Finally, no differences in EV surface markers expression were observed in the comparison of DeNovo PD *vs* Synucleinopathies and Synucleinopathies *vs* Tauopathies.

### EV profiling identifies disease-specific diagnostic signatures

In comparison with HC, multivariate logistic regression analysis (Figure 3A), corrected for age, sex, BMI, cohort, and number of experiments, associated four EV surface markers with iRBD diagnosis (CD3, CD9, CD25, CD56), one with Late PD (SSEA4), two with Synucleinopathies (CD146, CD209), and three with Tauopathies (CD8, CD45, CD62P). CD9 was associated with iRBD diagnosis in comparison with Tauopathies, while CD41b, CD63, and HLA-I were associated with Late PD diagnosis when compared to Synucleinopathies and Tauopathies.

**Figure 3.**
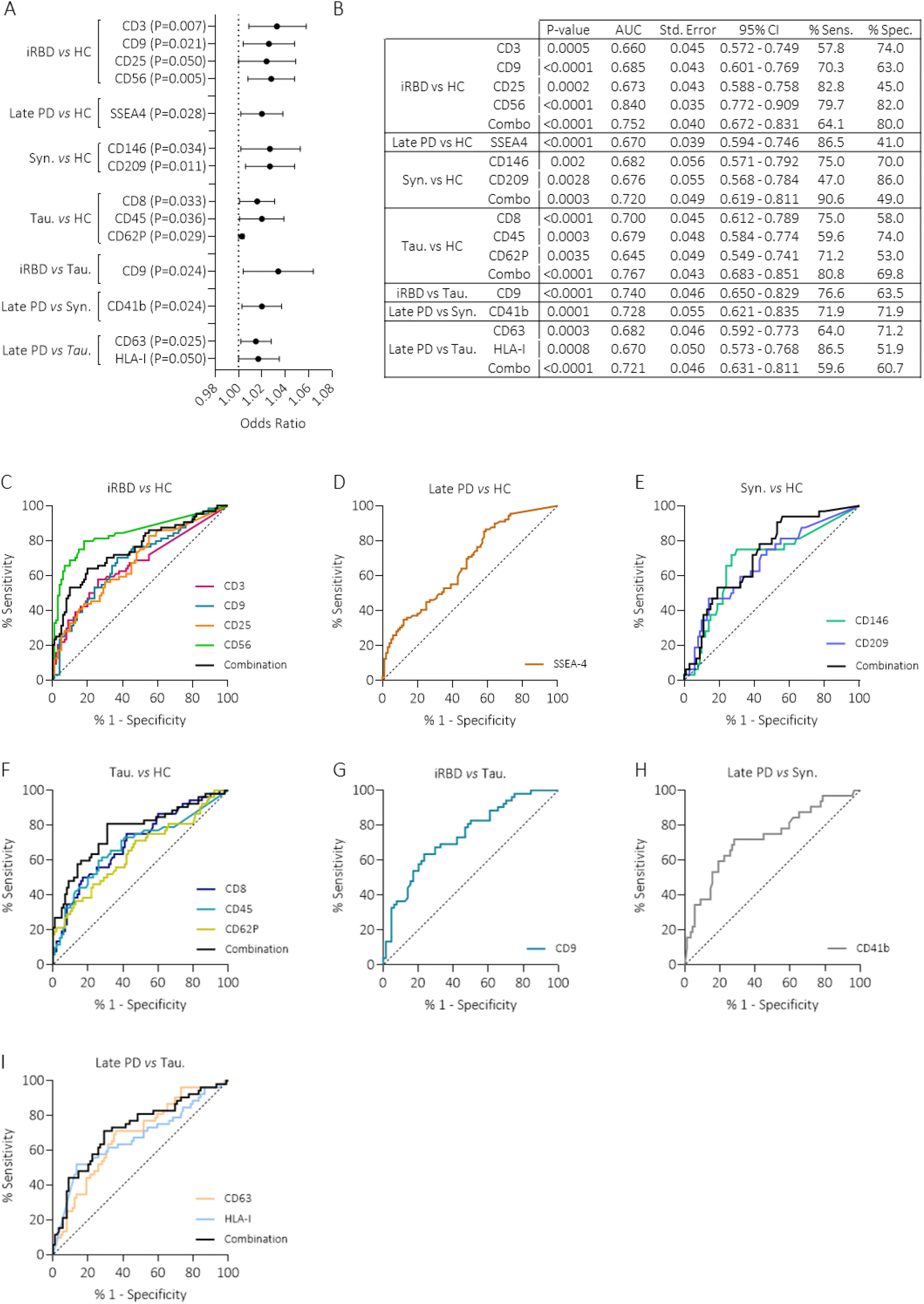
Diagnostic performance of EV-surface markers. (A) Forest plots showing the odds ratios and 95% confidence intervals of selected EV-surface markers that significantly discriminate each pair of groups; p-values for each marker are reported next to the corresponding comparison. (B) Summary table reporting, for each comparison and EV-surface marker (and their linear combination), the p-value, area under curve (AUC), standard error, 95% confidence interval (CI), and the corresponding sensitivity and specificity at the optimal cut-off. (C–I) ROC curves illustrating the diagnostic performance of single EV-surface markers and of their combined linear predictor for iRBD vs HC (C), Late PD vs HC (D), Synucleinopathies vs HC (E), Tauopathies vs HC (F), iRBD vs Tauopathies (G), Late PD vs Synucleinopathies (H), and Late PD vs Tauopathies (I). The referral line is reported (dashed line).

Markers significantly associated with the different diagnoses were tested to evaluate their diagnostic performance (Figure 3B-I). ROC curve analysis showed that all of them were able to discriminate between groups with good sensitivity and specificity. In particular, in the comparison between iRBD and HC, CD56 exhibited the best diagnostic performance, overcoming also the combination of all four markers together. In Late PD *vs* HC, SSEA4 has high sensitivity but low specificity, whereas in iRBD *vs* Tauopathies and Late PD *vs* Synucleinopathies, CD9 and CD41b showed moderate sensitivity and specificity, respectively. In Synucleinopathies *vs* HC, the combination of CD146 and CD209 achieved a really high sensitivity but low specificity. On the contrary, CD209 exhibits high specificity and low sensitivity, whereas CD146 displays moderate results. In Tauopathies *vs* HC, the combination of the three markers showed good sensitivity, while the best specificity was achieved with CD45. Finally, in Late PD *vs* Tauopathies, both CD63 and HLA-I performed slightly better than the combination of the two in terms of sensitivity, while the highest specificity was obtained with CD63, followed by the combination and then HLA-I.

### Late PD show disease-specific CSF-EV signature

The demographics and clinical scales of the 77 patients from the BioFIND cohort are reported in Supplementary Table 6. No differences were detected in age, sex, or BMI between Late PD and HC.

No differences in CSF-derived EV concentration and diameter were observed between the two groups. CD1c, CD29, CD69, and CD326 were significantly increased on the surface of EVs from Late PD compared to HC (Figure 4A-D, Supplementary Table 7). Among these, CD326 was associated with the clinical diagnosis of Late PD after adjusting for age, sex, and BMI by multivariate logistic regression analysis (P=0.017, OR=1.95, 95% CI [1.1-3.4]).

**Figure 4.**
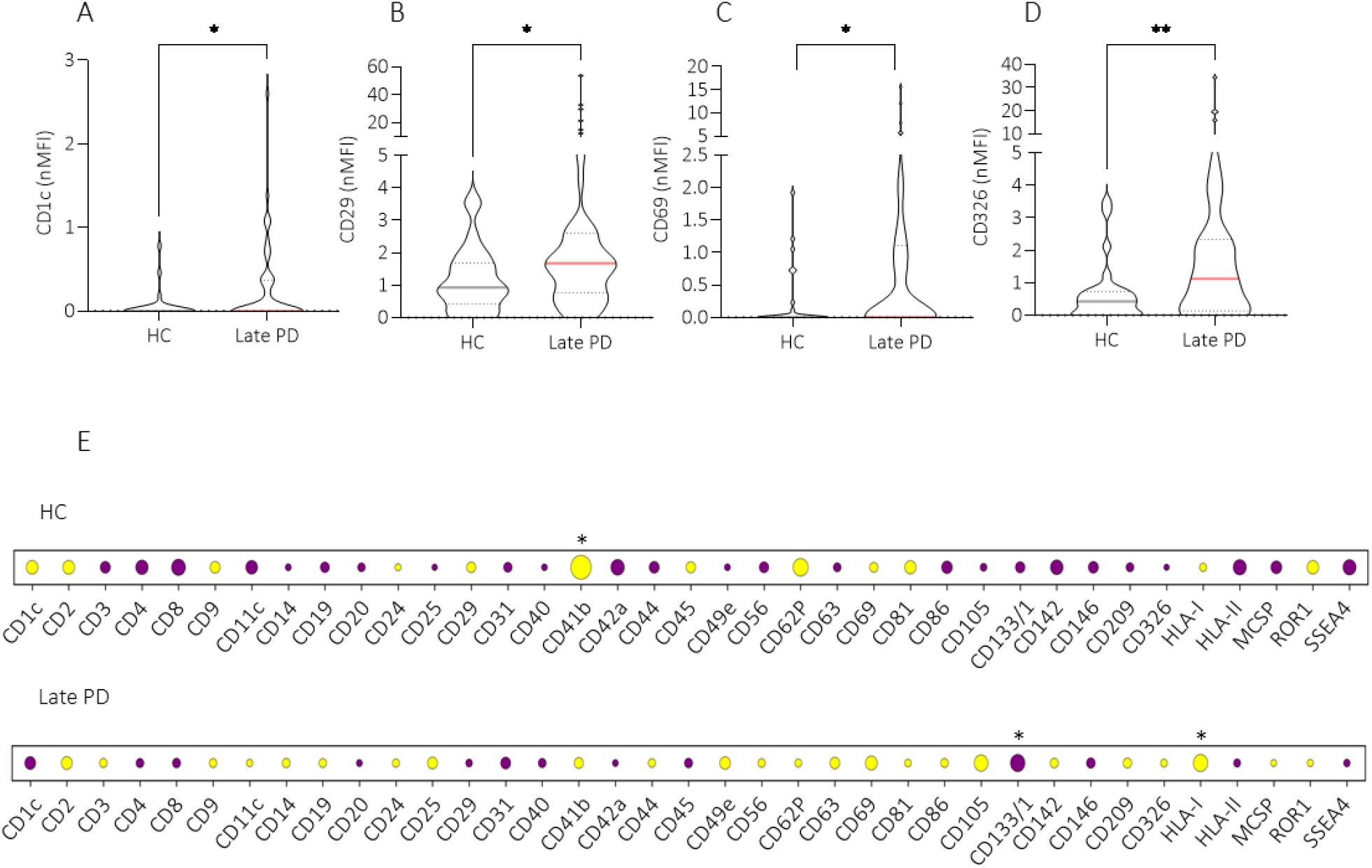
Differential expression of CSF-derived EVs. (A-D) Violin plots showing the median (colour line) and interquartile range (dashed line) of normalized mean fluorescence intensity (nMFI) for the indicated EV surface markers (CD1c, CD29, CD69, and CD326) in HC and LatePD from the BioFIND cohort. P-values <0.05 were considered statistically significant: *p<0.05, **p<0.01. (E) Dot plots showing correlations of EV antigens measured in plasma and CSF. The color indicates direct (yellow) or indirect (purple) correlation, while the dimension the strength of correlation (a higher radius corresponds to a stronger Pearson’s R value). * Highlights significant correlations.

Correlating the expression of the 37 surface markers obtained from plasma- and CSF-derived EVs of the same 77 patients, a moderate positive correlation of CD41b (P=0.013, R=482) was seen in HC, while a low negative correlation of CD133 (P=0.038, R=-0.291) and a low positive correlation of HLA-I (P=0.040, R=0.288) were observed in Late PD (Figure 4E, Supplementary File 2).

### Machine learning algorithms enable accurate prodromal classification

The population was randomly split into a training (n=253) and a validation cohort (n=125). No significant differences were observed between the two groups in age (training 67±10y, validation 67±10y, P=0.920), sex (training 60% male, validation 67% male, P=0.214), or BMI (training 25.5kg/m2 [22.9-28.9], validation 25.9kg/m2 [23.5-28.7], P=0.639). Within the individual groups (Supplementary Table 8), Late PD in the validation cohort had a higher LEDD intake than the training cohort, while the Tauopathy group in the validation cohort had a higher percentage of males than the training cohort.

Using all 37 surface antigens, 480 rRF diagnostic models were trained on the training cohort and then evaluated separately in the validation cohort (Supplementary File 2). The rRF classifiers with the highest performance at validation are shown in Figure 5. In the training cohort, the highest accuracy was achieved in predicting Synucleinopathies vs HC (94.3%; sensitivity 95.2%, specificity 94.0%), followed by iRBD vs HC (91.7%; sensitivity 90.6%, specificity 92.5%), DeNovo PD vs HC (89.4%; sensitivity 85.1%, specificity 91.0%), Late PD vs HC (85%; sensitivity 85%, specificity 85%), and Tauopathies vs HC (72.5%; sensitivity 88.5%, specificity 64.2%). In the validation cohort, the highest accuracy was obtained in predicting iRBD vs HC (77.8%; sensitivity 76.1%, specificity 78.7%), followed by DeNovo PD vs HC (76.6%; sensitivity 57.1%, specificity 84.8%), Tauopathy vs HC (76.0%; sensitivity 82.3%, specificity 72.7%), Synucleinopathy vs HC (70.5%; sensitivity 45.4%, specificity 78.7%), and Late PD vs HC (66.1%; sensitivity 72.4%, specificity 60.0%).

**Figure 5.**
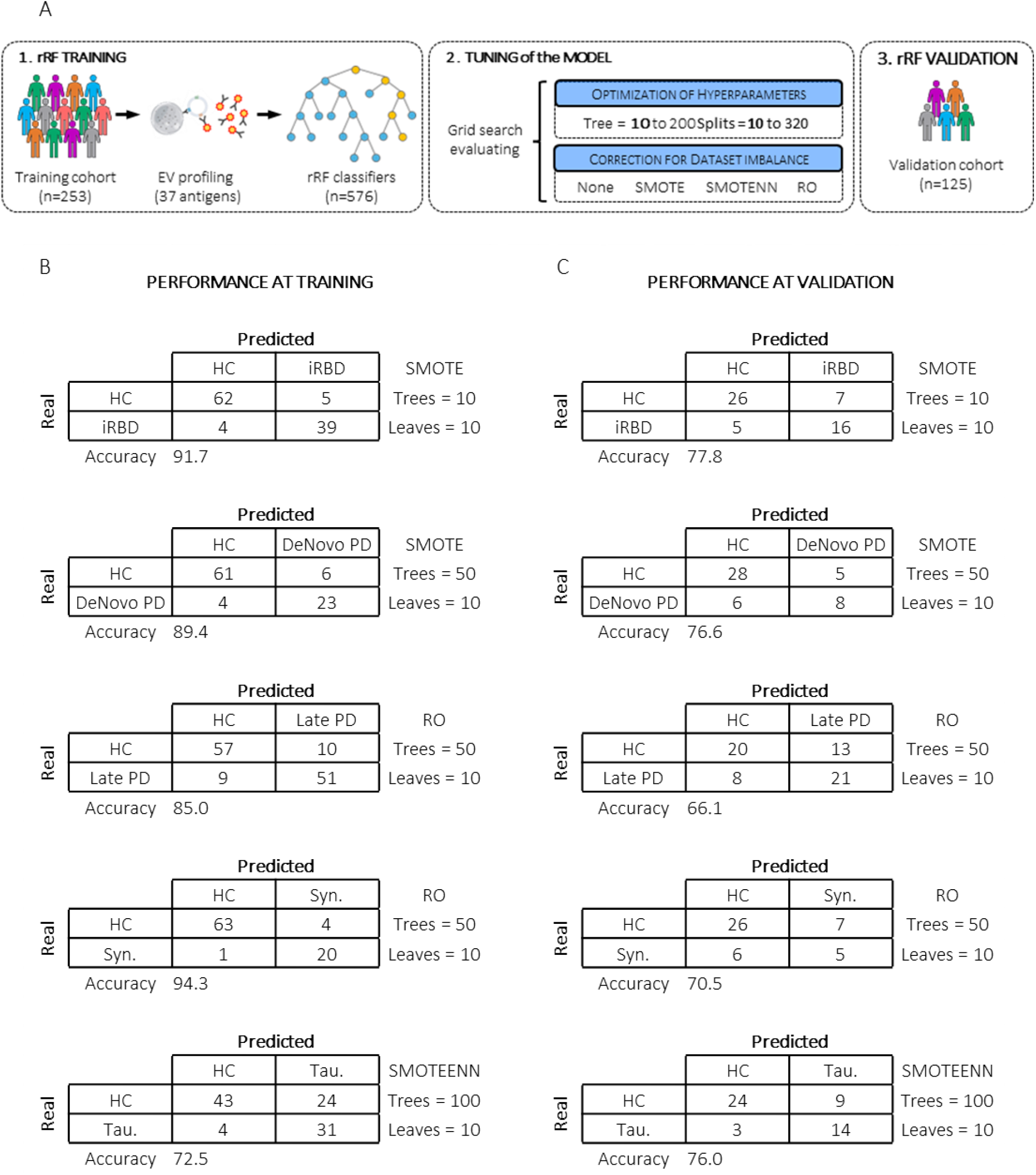
Machine learning workflow and classification performance across groups. (A) Schematic representation of the machine learning pipeline, including feature extraction, model construction, hyperparameter optimization, and correction for dataset imbalance (SMOTE, RO, SMOTEENN). (B) Confusion matrices showing classification performance at training and (C) validation for HC versus iRBD, DeNovo PD, Late PD, Synucleinopathies, and Tauopathies groups. For each comparison, the applied resampling strategy and model hyperparameters (number of trees and leaves) are reported. Overall accuracy (%) is indicated below each matrix.

## Discussion

In this multicenter study, we characterized plasma EVs across a large cohort comprising healthy subjects, patients with iRBD, DeNovo and Late PD, other Synucleinopathies, and Tauopathies. By integrating multiparametric flow cytometry and machine learning, we identified stage-specific EV surface marker profiles that reflect distinct immune and cellular signatures along the synucleinopathy/tauopathy continuum.

The analysis of 37 EV surface antigens revealed specific expression signatures consistent with known immunopathological features. iRBD displayed a general increase in almost all immune antigens, with marked overexpression of CD3 (T lymphocytes), CD56 (NK cells), and CD25 (activation and Treg marker) compared with HC. Reports of reduced CD3+ T cells^41^, increased NK cells^42^, and unchanged Treg^41^ cells in PBMCs from iRBD patients suggest that the overexpression of these EV surface antigens reflects early peripheral immune activation and reprogramming rather than changes in cell count. EV surface markers may provide information about the cellular origin of the vesicles, but they can also have a functional role in targeting specific cells. A recent study showed elevated T-cell reactivity against α-synuclein conformers in prodromal PD compared to HC^43^; thus, the release of CD3^+^/CD56^+^ EVs from activated T and NK cells may represent compensatory immune surveillance against emerging α-synuclein conformers, activating antigen-presenting cells or dendritic cells to amplify adaptive responses before neuronal loss becomes irreversible^44^. CD25^+^ EVs instead may bind circulating IL-2, attenuating T-cell responses, mimicking T-reg cell function, preventing cytokine storm from overzealous T/NK activation, and maintaining immune homeostasis during prodromal stress^45–47^.

CD56 showed high diagnostic performance in distinguishing iRBD from HC, suggesting that NK-derived EVs may represent an early biomarker of synucleinopathic conversion. Because CD56 is also a neuronal marker, CD56^+^ EVs could either have a neuronal origin (less likely given the very low proportion of neuron-derived EVs in plasma and the higher number of NK cells^48^) or a peripheral origin with neuronal targets. CD56^+^ EVs may show enhanced adhesion and interaction with tissues expressing compatible ligands (including neural compartments), facilitating peripheral-central nervous system crosstalk, particularly under prodromal conditions associated with increased blood-brain barrier permeability^49,50^.

Finally, CD9 overexpression vs both HC and Tauopathies may suggest enhanced EV biogenesis and release. However, CD81 and CD63 are much less expressed than in other groups, and no overall increase in EV number was detected in iRBD. Across groups, CD9 is high in iRBD, whereas other groups show similar levels, suggesting selective activation of specific cell types or compartments that preferentially release CD9+ EVs at the prodromal stage. CD63 increases in iRBD, De Novo, and Late PD but decreases in Synucleinopathies and Tauopathies, suggesting initial upregulation of endosomal-exosomal trafficking to handle pathological proteins or modulate the microenvironment^51^, later impaired in more aggressive disease. CD81 remains relatively stable in iRBD and PD but rises in Synucleinopathies and Tauopathies, potentially reflecting EV populations more specific to parkinsonian syndromes^52^. Overall, these data suggest dynamic regulation of EV release across diseases. The general increase in Late PD could reflect heightened systemic inflammation or ongoing neurodegenerative stress, whereas reduced levels in early PD and Tauopathies may indicate impaired vesicle biogenesis or selective vesicle clearance.

Late PD showed increased CD41b, HLA-I, and SSEA4 expression compared to other groups, indicating engagement of platelet, adaptive immune, and neuronal components. CD41b, a canonical marker of platelet-derived EVs, has been associated with PD diagnosis in our previous studies^25,40^, supporting a role for platelet activation and platelet-EV trafficking in disease pathophysiology. PD patients show platelet hyperactivation^53^ and morphological changes^54^, and coagulation dysfunction with elevated fibrinogen, impaired fibrinolysis, and platelet hyperactivity likely creates a hypercoagulable state that elevates thrombotic risk in PD^55^. Inflammation and oxidative stress drive this coagulopathy-neurodegeneration axis, with platelet α-synuclein accumulation disrupting aggregation and amplifying systemic microvascular damage^55^. In this context, platelet-derived EVs may contribute to systemic inflammation, endothelial dysfunction, and hypercoagulability in neurodegenerative disorders and may interact with misfolded α-synuclein in circulation. In contrast, patients with other Synucleinopathies, such as LBD^56^, show a non-significant reduction in platelet count compared with PD, consistent with differential platelet impairment and behavior across neurodegenerative diseases. Accordingly, we observed decreased CD41b+ EVs in Synucleinopathy patients. HLA-I upregulation on plasma EVs aligns with genetic, neuropathological, and functional evidence implicating HLA-I signaling in PD neuroinflammation and neurodegeneration. GWAS repeatedly identify HLA alleles in PD^57^, suggesting that altered antigen presentation modulates disease susceptibility and progression. Neuropathologically, HLA-I is expressed on nigral dopaminergic neurons in post-mortem PD brains^58^, where it facilitates CD8+ T-cell recognition of neuronal antigens, including α-synuclein-derived peptides and mitochondrial proteins exposed by PINK1/Parkin dysfunction^59^. HLA-I+ EVs in Late PD may represent a peripheral mirror of central dysregulation, potentially carrying neuronal/astrocytic antigens or amplifying systemic T-cell activation through EV-mediated antigen transfer. In our study, HLA-I was associated with PD diagnosis, especially compared to Tauopathies, which instead are characterized by HLA-II overexpression.

Finally, SSEA4, a glycolipid marker of neural progenitors^60^, discriminates Late PD from HC with high sensitivity, suggesting that neuronal/progenitor-derived EVs may contribute to the circulating pool and reflect chronic synaptic-neuroglial remodeling. No studies have directly associated this protein with PD so far; however, specific inhibitors targeting SSEA-4 have been developed to block its function and modulate signaling pathways essential for maintaining stem cell pluripotency, thereby promoting cell differentiation. These inhibitors are currently explored in cancer therapy, and could eventually be repurposed to stimulate the generation of specific neuronal cell types for the treatment of neurodegenerative disorders^61^.

Regarding atypical parkinsonisms, we confirmed increased CD146 and CD209 on the surface of plasmatic EVs from Synucleinopathy patients compared to HC^25,40^. CD146, an adhesion molecule overexpressed on blood-brain barrier endothelial cells in inflammatory conditions, promotes transmigration of inflammatory cells into the CNS^62^. Thus, elevated CD146+ EVs may reflect blood-brain barrier dysregulation. CD209, present on macrophages and dendritic cells, is fundamental for pathogen recognition and T cell activation, and likely reflects activation of these cells when increased on EV surface. Conversely, Tauopathies displayed predominant upregulation of cytotoxic lymphocyte and platelet markers (CD8, CD45, CD62P). CD8 and CD45 enrichment on Tauopathy EVs aligns with neuropathological evidence of CD8+ T-cell infiltration in tau-affected brain regions, where cytotoxic lymphocytes cluster around astrocytic tau inclusions and perivascular spaces, correlating with tau epitopes and glial reactivity^63,64^. CD62P, a platelet and endothelial activation marker, has been extensively implicated via platelet-derived EVs^65^ in coagulation, endothelial dysfunction, and inflammation, processes increasingly recognized in Tauopathies and other proteinopathies^66^.

In the BioFind cohort, CD1c, CD29, CD69, and CD326 were overexpressed on CSF-EVs in PD compared to HC. CD1c and CD29 had already been identified in our pilot study on an independent small cohort as enriched on CSF-derived EVs in PD^40^, supporting their relevance. In the overall cohort, all four markers also showed higher expression in Late PD vs HC in plasma, although without reaching statistical significance.

CD1c and CD69 are immune activation markers and may indicate an immuno-inflammatory CNS response involving CD1c^+^ antigen-presenting dendritic cells and recruitment/activation of CD69^+^ effector cells (B, T, and NK cells). Although dendritic cells are typically scarce in CSF, they are recruited from blood under pathological conditions, including PD^67,68^. Thus, increased CD1c+ EVs may therefore indicate increased number or activation of CSF dendritic cells. Furthermore, CD1c recognizes foreign lipid antigens, and CD1c+ dendritic cells typically present these antigens to CD4+ T helper cells. A recent study reported a rise in these cells within PD CSF and examined CD28 as an activation marker^69^. CD28 expression was elevated on CD4^+^ T cells in blood but not in the CSF; however, because CD28 and CD69 represent distinct activation states, it remains unclear whether CSF CD4^+^ T cells express CD69. The same uncertainty applies to B cells and NK cells; their increased presence in the CSF is documented^69,70^, but their activation status remains undefined. CD29, on the other hand, a transmembrane glycoprotein widely expressed across immune, endothelial, epithelial, and neural/glial cells during migration and trafficking, may increase on CSF-EVs owing to contributions from multiple cell types reflecting changes in activation, adhesion, and migration.

CD326^+^ CSF-EVs are particularly noteworthy due to their association with PD diagnosis. CD326, a transmembrane glycoprotein predominantly expressed by epithelial cells, supports epithelial cohesion and adhesion and mediates targeting of other CD326^+^ cells^71^. In CSF, CD326^+^ EVs may reflect epithelial blood-brain barrier alterations or origin from choroid plexus epithelial cells, consistent with structural and functional choroid plexus alterations and enlargement in PD^72,73^, which correlate with worsening motor severity and frontal/executive dysfunction^72,73^. Together, these four markers suggest an intense state of active neuroinflammation characterized by immune cell recruitment (CD1c), migration (CD29), and early stimulation (CD69), potentially linked to epithelial and blood–brain barrier dysfunction (CD326).

Finally, the cross-compartment correlations of EV surface marker expression between plasma and CSF revealed a differential redistribution of EVs between the two compartments. HC showed mainly inverse correlations while PD direct. This discrepancy may reflect an altered blood–brain/CSF barrier permeability, changes in EV trafficking, or compensatory and clearance mechanisms that arise in Late PD as the central and peripheral EV pools become progressively more associated.

RF models provided high discrimination accuracy between disease categories and controls, particularly for iRBD. Robust cross-validation performance underscores the diagnostic potential of combined EV signatures. Consistent accuracy for iRBD vs HC in training (91.7%) and validation (77.8%) cohorts supports the ability of EV profiling to identify prodromal synucleinopathy with clinically meaningful precision. The stability of the iRBD vs HC model is compatible with a distinct immune signature that is less fragmented by clinical variability than in more heterogeneous groups. Lower predictive performance for Late PD and Synucleinopathies in the validation set likely reflects greater biological heterogeneity, pre-analytical variability, and clinical/pharmacological confounding.

Overall, these findings support a model of progressive EV remodeling across the neurodegenerative spectrum, in which immune- and endothelial-derived EVs predominate in early stages, while neuronal and platelet components become more prominent in advanced disorders. The combination of CD3, CD9, CD25, and CD56 emerges as a promising peripheral biomarker panel for iRBD, whereas SSEA4 and CD146-CD209 potentially delineate distinct molecular signatures of Late PD and Synucleinopathies, respectively. The inclusion of CSF-derived markers such as CD326 further enhances the capacity of EVs to bridge peripheral and central pathology.

This study has several limitations that should be considered when interpreting the results. The marked male predominance in iRBD and the higher proportion of females in HC introduce potential sex-related confounding that, despite statistical adjustment, may partially influence EV immunophenotypes. Similarly, differences in age between Tauopathies and other groups may shape baseline EV signatures and complicate direct comparisons across the whole proteinopathy spectrum. Moreover, the panel used here was not optimized to define EV cell-of-origin; many markers are shared across leukocyte subsets, endothelial cells, and platelets, limiting precise attribution of specific vesicle populations and mechanistic interpretation of surface changes. Finally, the cross-sectional design precludes any inference on temporal trajectories or conversion risk, particularly in iRBD, where longitudinal EV profiling would be crucial to establish predictive value for phenoconversion to overt Synucleinopathy.

Future longitudinal studies, especially in iRBD cohorts, are recommended to assess EV signatures as predictors of progression to PD, MSA, or LBD and to identify optimal temporal windows. Integration of surface phenotyping with cargo quantification, alongside refined cell-of-origin profiling, could elucidate EVs’ role in neurodegeneration. Ultimately, multidimensional EV-based biosignatures, combined with established modalities, may facilitate early diagnosis, risk stratification, and therapeutic monitoring across Synucleinopathy/Tauopathy spectra.

## Author contributions

E.V.: acquisition, statistical analysis, interpretation of data, and manuscript writing. J.B., A.B.: statistical analysis, interpretation of data, and machine learning algorithm development. S.B., A.Y., S.P.: data acquisition. I.R.B., I.B., L.B., M.G.B., G.C., R.K., A.R., S.V.M., S.H., S.R., S.Mi., G.B., C.S., J.P., J.K., F.P., G.K., M.M., S.G., A.K.L.: patient enrollment. L.B.: study design and data interpretation. G.M.: study design, patient enrollment, data interpretation, and manuscript writing. All authors revised the manuscript for intellectual content and agreed on the published version of the manuscript.

## Data Availability

Raw data that support the findings of this manuscript are available upon reasonable request to the corresponding author.

## Acknowledgments

The authors are grateful to patients and their relatives who participated in this study.

Part of the data used in the preparation of this article were obtained from the Fox Investigation for New Discovery of Biomarkers (“BioFIND”) database (http://biofind.loni.usc.edu/). For up-to-date information on the study, visit https://www.michaeljfox.org/biospecimens. BioFIND is sponsored by The Michael J. Fox Foundation for Parkinson’s Research (MJFF) with support from the National Institute for Neurological Disorders and Stroke (NINDS).

