## Supplementary material for "Plasmatic immune extracellular vesicle profiles identify prodromal and early stages of Parkinson’s disease": Supplem.File1

Supplementary Table 1. Multicentric enrollment

| | Lugano | Bologna | Barcellona | Torino | St. Gallen | BioFIND | Total |
|---|---|---|---|---|---|---|---|
| HC | 74 | - | - | - | - | 26 | 100 |
| iRBD | 18 | 43 | - | - | 3 | - | 64 |
| DeNovo | 30 | - | - | 7 | 4 | - | 41 |
| Late PD | 34 | - | 1 | - | 3 | 51 | 89 |
| Syn. | 21 | - | 9 | 2 | - | - | 32 |
| Tau. | 23 | - | 26 | 2 | 1 | - | 52 |
| Total | 200 | 43 | 36 | 11 | 11 | 77 | **378** |

The Movement Disorder Unit in Lugano (Switzerland) enrolled a total of 200 patients of all categories. Eleven samples of DeNovo and atypical Parkinsonisms arrived from the University Hospital of Torino (Italy). The Kantonsspital St. Gallen (Switzerland) provided samples from PD and iRBD (n=11). The Sleep Unit in Bologna (Italy) enrolled 43 subjects with primary iRBD. The Sant Pau Hospital in Barcelona (Spain) provided mainly atypical Parkinsonism, both Synucleinopathies (n=9) and Tauopathies (n=26). Finally, 77 samples of HC and Late PD were part of the BioFIND cohort of the MJFF. Subjects from Lugano, Torino, Bologna, and St. Gallen were prospectively recruited according to the NSI-PD001 study protocol, while samples from Barcelona were collected as part of a local clinical study, using the same protocols for plasma collection and storage.

Supplementary Table 2. Demographic data, medical history, and clinical scores

| Variable | HC (n=100) | iRBD (n=64) | DeNovo PD (n=41) | Late PD (n=89) | Syn (n=32) | Tau (n=52) |
|---|---|---|---|---|---|---|
| Age (y) n=378 | 65 ± 11 | 68 ± 7 | 64 ± 11 | 68 ± 9 | 69 ± 9 | 72 ± 12 |
| Sex (%M) n=378 | 51.0 | 84.0 | 56.0 | 66.0 | 56.0 | 60.0 |
| BMI n=309 | 25.0 [22.7-27.8] | 26.7 [24.3-28.7] | 25.1 [22.8-29.2] | 25.7 [23.3-29.2] | 24.7 [22.0-26.5] | 25.8 [23.0-29.9] |
| Disease Duration (y) n=278 | - | 6 ± 5 | 2 ± 1 | 7 ± 3 | 4 ± 3 | 4 ± 2 |
| Age of onset (y) n=278 | - | 62 ± 8 | 62 ± 11 | 61 ± 9 | 65 ± 11 | 69 ± 7 |
| H&Y n=214 | - | - | 1.0 [1.0-2.0] | 2.0 [2.0-3.0] | 3.0 [3.0-4.5] | 4.0 [3.0-4.0] |
| MDS-UPDRS I n=134 | - | 8.0 [6.3-15.8] | 6.5 [3.3-13.8] | 9.0 [6.0-15.0] | 10.5 [8.0-12.0] | 8.0 [6.0-15.0] |
| MDS-UPDRS II n=134 | - | 1.0 [0.0-2.0] | 5.0 [3.0-9.8] | 9.0 [4.0-12.0] | 13.05 [9.5-15.0] | 10.0 [8.0-24.0] |
| MDS-UPDRS III n=278 | - | 4.0 [2.0-6.0] | 13.5 [8.5-23.0] | 29.0 [18.0-45.5] | 31.0 [22.5-41.0] | 34.0 [24.3-41.0] |
| MDS-UPDRS Total n=134 | - | 13.5 [7.3-32.5] | 28.0 [18.3-47.5] | 50.0 [30.-73.0] | 50.5 [39.8-63.0] | 47.0 [35.0-90.0] |
| COMPASS 31 – OH n=167 | - | 0.0 [0.0-16.0] | 0.0 [0.0-16.0] | 0.0 [0.0-16.0] | 0.0 [0.0-12.0] | 0.0 [0.0-14.0] |
| COMPASS 31 - VM n=167 | - | 0.0 [0.0-0.0] | 0.0 [0.0-0.0] | 0.0 [0.0-0.0] | 0.0 [0.0-0.0] | 0.0 [0.0-0.0] |
| COMPASS 31 - SM n=167 | - | 0.0 [0.0-4.8] | 2.1 [0.0-4.2] | 0.5 [0.0-4.2] | 4.2 [0.0-6.3] | 2.1 [0.0-5.7] |
| COMPASS 31 - GI n=167 | - | 4.5 [0.0-6.3] | 3.6 [0.9-7.7] | 4.2 [0.0-6.3] | 5.4 [0.0-6.3] | 3.6 [0.0-6.3] |
| COMPASS 31 - BL n=167 | - | 1.1 [0.0-2.2] | 0.0 [0.0-1.1] | 0.0 [0.0-1.0] | 1.1 [0.0-3.3] | 0.0 [0.0-1.1] |
| COMPASS 31 - PM n=167 | - | 0.8 [0.0-1.4] | 0.0 [0.0-1.7] | 0.0 [0.0-0.0] | 0 [0.0-0.0] | 0 [0.0-0.0] |
| COMPASS 31 - Total n=167 | - | 12.7 [7.0-28.4] | 11.4 [4.7-29.7] | 11.6 [4.2-21.7] | 14.4 [9.2-21.3] | 12.0 [6.4-24.7] |
| BDI-II n=163 | - | 7.0 [2.0-11.0] | 7.5 [2.3-14.5] | 6.0 [3.0-9.5] | 10.0 [6.0-14.0] | 11.0 [6.5-16.0] |
| MMSE n=230 | - | 29.0 [28.0-30.0] | 30.0 [28.3-30.0] | 29.0 [28.0-30.0] | 28.0 [25.0-29.0] | 27.0 [24.0-28.0] |
| MoCA n=230 | - | 26.0 [23.0-28.0] | 26.5 [23.0-28.8] | 27.0 [25.0-29.0] | 23.0 [18.0-26.0] | 20.0 [16.8-24.3] |
| RBDSQ n=180 | - | 10.0 [8.0-11.0] | 3.0 [2.0-7.0] | 3.0 [1.06.0] | 4.0 [1.5-6.0] | 2.0 [1.0-4.0] |
| LEDD n=278 | - | 0.0 [0.0-0.0] | 100.0 [0.0-157.0] | 375.0 [223.8-585.8] | 300.0 [0.0-500.0] | 500.0 [77.5-750.0] |

For each group, demographic, medical, and clinical characteristics are reported as mean±standard deviation (SD), percentage (%), or median and [interquartile range]. BMI= Body Mass Index; H&Y = Hoehn and Yahr scale; MDS-UPDRS = Movement Disorder Society–Unified Parkinson's Disease Rating Scale; COMPASS 31 (OH= orthostatic hypotension, VM= vasomotor, SM= sudomotor, GI= gastrointestinal, BL = Bladder, PM= pupillomotor); BDI-II = Beck Depression Inventory II; MMSE = Mini-Mental State Examination; MoCA = Montreal Cognitive Assessment; RBDSQ = REM sleep Behavior Disorder Screening Questionnaire, LEDD = Levodopa Equivalent Daily Dose. "n" refers to the number of subjects for whom data is available.

Supplementary Table 3. Demographic data, medical history, and clinical scores- statistics

| Variable | HC vs. iRBD | HC vs. DeNovo | HC vs. Late PD | HC vs. Syn. | HC vs. Tau. | iRBD vs. DeNovo | iRBD vs. Late PD | iRBD vs. Syn. | iRBD vs. Tau. | DeNovo vs. Late PD | DeNovo vs. Syn. | DeNovo vs. Tau. | Late PD vs. Syn. | Late PD vs. Tau. | Syn. vs. Tau |
|---|---|---|---|---|---|---|---|---|---|---|---|---|---|---|---|
| Age (y) | 0.833 | >0.999 | 0.344 | 0.490 | **<0.0001** | 0.595 | >0.999 | >0.999 | **0.030** | 0.311 | 0.335 | **<0.0001** | >0.999 | **0.024** | 0.683 |
| Sex (%M) | **0.000** | 0.582 | **0.033** | 0.605 | 0.312 | **0.001** | **0.012** | **0.003** | **0.003** | 0.263 | 0.990 | 0.733 | 0.311 | 0.426 | 0.761 |
| BMI | 0.565 | >0.999 | >0.999 | >0.999 | >0.999 | >0.999 | >0.999 | 0.551 | >0.999 | >0.999 | >0.999 | >0.999 | >0.999 | >0.999 | >0.999 |
| Disease Duration (y) | - | - | - | - | - | **<0.0001** | **0.031** | 0.251 | 0.054 | **<0.0001** | **0.002** | **0.001** | **<0.0001** | **<0.0001** | >0.999 |
| Age of onset (y) | - | - | - | - | - | >0.999 | >0.999 | 0.573 | **<0.0001** | >0.999 | >0.999 | **0.003** | 0.139 | **<0.0001** | 0.591 |
| LEDD | - | - | - | - | - | **0.011** | **<0.0001** | **<0.0001** | **<0.0001** | **<0.0001** | 0.175 | **<0.0001** | 0.229 | >0.999 | 0.565 |
| H&Y | - | - | - | - | - | - | - | - | - | **0.004** | **<0.0001** | **<0.0001** | **0.001** | **<0.0001** | >0.999 |
| MDS-UPDRS I | - | - | - | - | - | >0.999 | >0.999 | >0.999 | >0.999 | >0.999 | >0.999 | >0.999 | >0.999 | >0.999 | >0.999 |
| MDS-UPDRS II | - | - | - | - | - | 0.423 | **0.026** | **0.002** | **0.007** | 0.362 | **0.016** | 0.111 | 0.425 | >0.999 | >0.999 |
| MDS-UPDRS III | - | - | - | - | - | **<0.0001** | **<0.0001** | **<0.0001** | **<0.0001** | **0.003** | **0.012** | **0.001** | >0.999 | >0.999 | >0.999 |
| MDS-UPDRS Total | - | - | - | - | - | >0.999 | 0.050 | 0.097 | 0.093 | **0.003** | 0.131 | 0.133 | >0.999 | >0.999 | >0.999 |
| COMPASS 31 - OH | - | - | - | - | - | >0.999 | >0.999 | >0.999 | >0.999 | >0.999 | >0.999 | >0.999 | >0.999 | >0.999 | >0.999 |
| COMPASS 31 - VM | - | - | - | - | - | >0.999 | >0.999 | >0.999 | >0.999 | >0.999 | >0.999 | >0.999 | >0.999 | >0.999 | >0.999 |
| COMPASS 31 - SM | - | - | - | - | - | >0.999 | >0.999 | >0.999 | >0.999 | >0.999 | >0.999 | >0.999 | 0.953 | >0.999 | >0.999 |
| COMPASS 31 - GI | - | - | - | - | - | >0.999 | >0.999 | >0.999 | >0.999 | >0.999 | >0.999 | >0.999 | >0.999 | >0.999 | >0.999 |
| COMPASS 31 - BL | - | - | - | - | - | 0.670 | 0.089 | >0.999 | >0.999 | >0.999 | 0.098 | >0.999 | **0.014** | >0.999 | 0.458 |
| COMPASS 31 - PM | - | - | - | - | - | >0.999 | **0.001** | **0.008** | **0.015** | 0.089 | 0.218 | 0.329 | >0.999 | >0.999 | >0.999 |
| COMPASS 31 - Total | - | - | - | - | - | >0.999 | >0.999 | >0.999 | >0.999 | >0.999 | >0.999 | >0.999 | >0.999 | >0.999 | >0.999 |
| BDI-II | - | - | - | - | - | >0.999 | >0.999 | 0.441 | **0.030** | >0.999 | >0.999 | 0.827 | 0.594 | 0.059 | >0.999 |
| MMSE | - | - | - | - | - | 0.987 | >0.999 | 0.125 | **0.000** | >0.999 | **0.004** | **<0.0001** | **0.003** | **<0.0001** | >0.999 |
| MoCA | - | - | - | - | - | >0.999 | 0.120 | 0.992 | **0.002** | >0.999 | 0.234 | **0.000** | **0.005** | **<0.0001** | 0.961 |
| RBDSQ | - | - | - | - | - | **<0.0001** | **<0.0001** | **0.000** | **<0.0001** | >0.999 | >0.999 | >0.999 | >0.999 | >0.999 | >0.999 |

For each multiple comparison, the P-value is reported. P-values lower than 0.05 were considered significant and shown in bold.

Supplementary Table 4. Nanoparticle characterization and Normalized Median Fluorescence Intensity of all 37 EV-surface markers

| Variable | HC (n=100) | iRBD (n=64) | DeNovo PD (n=41) | Late PD (n=89) | Synucleinopathies (n=32) | Tauopathies (n=52) |
|---|---|---|---|---|---|---|
| EVs All (N°/mL) *E10 | 4.6 [1.0-41.0] | 10.0 [2.7-30.0] | 23.0 [5.7-96.0] | 2.2 [0.8-1600.0] | 520.0 [1.4-1300.0] | 3.8 [1.3-240.0] |
| EVs 30-150nm (N°/mL) *E10 | 3.0 [0.7-29.0] | 7.4 [2.0-22.0] | 18.0 [3.5-75.0] | 1.5 [0.5-1300.0] | 440.0 [1.1-1100.0] | 2.8 [0.9-170.0] |
| EVs 151-500nm (N°/mL) *E10 | 1.6 [0.3-10.0] | 3.2 [1.0-7.6] | 4.0 [2.1-16.0] | 0.8 [0.3-3000.0] | 6.0 [0.5-3500.0] | 0.8 [03-6.5] |
| Diameter (nm) | 140.2 [125.1-150.8] | 135.3 [120.4-151.4] | 125.3 [100.7-144.7] | 140.8 [130.5-149.4] | 124.7 [102.7-152.0] | 132.8 [119.9-147.8] |
| Mean CD9-CD63-CD81 (MFI) | 10.5 [4.8-19.1] | 9.1 [4.5-15.1] | 5.7 [2.9-8.1] | 16.3 [9.6-30.7] | 6.9 [3.5-12.7] | 5.0 [3.0-7.3] |
| CD1c (nMFI) | 6.4 [0.2-12.5] | 10.5 [1.7-24.1] | 5.9 [0.0-20.2] | 9.0 [2.3-24.9] | 10.4 [4.6-37.7] | 10.5 [1.9-34.9] |
| CD2 (nMFI) | 2.5 [0.0-7.7] | 9.0 [3.9-24.2] | 7.7 [1.4-26.7] | 3.5 [0.0-11.1] | 6.4 [0.0-15.1] | 4.0 [0.0-18.5] |
| CD3 (nMFI) | 2.2 [0.0-10.1] | 11.2 [0.0-32.6] | 6.7 [0.0-45.8] | 5.8 [0.0-22.2] | 12.0 [0.5-33.8] | 7.2 [0.0-33.4] |
| CD4 (nMFI) | 4.4 [0.0-13.8] | 4.7 [0.0-22.5] | 1.7 [0.0-15.4] | 7.2 [2.1-24.3] | 5.5 [0.0-21.0] | 12.2 [1.2-33.3] |
| CD8 (nMFI) | 32.3 [21.5-54.0] | 58.5 [39.1-83.6] | 56.2 [41.0-93.5] | 35.3 [19.6-62.5] | 47.2 [31.1-70.2] | 57.1 [31.5-75.6] |
| CD9 (nMFI) | 80.6 [55.1-105.2] | 109.0 [82.8-160.6] | 63.8 [47.1-92.2] | 82.8 [55.1-121.6] | 71.1 [42.2-97.7] | 65.8 [39.1-101.0] |
| CD11c (nMFI) | 4.1 [0.0-11.2] | 2.4 [0.0-22.1] | 3.4 [0.0-8.9] | 7.8 [0.9-21.9] | 12.3 [1.1-21.3] | 7.1 [0.0-29.5] |
| CD14 (nMFI) | 5.1 [0.0-16.2] | 2.3 [0.0-16.0] | 5.9 [0.0-22.5] | 9.0 [1.7-26.5] | 11.8 [0.0-34.9] | 7.2 [0.0-31.7] |
| CD19 (nMFI) | 7.8 [1.4-19.2] | 15.2 [5.3-31.3] | 13.5 [4.6-23.6] | 16.2 [6.5-37.4] | 14.2 [4.4-45.7] | 7.7 [0.0-27.0] |
| CD20 (nMFI) | 8.8 [0.0-16.8] | 17.8 [4.8-41.1] | 7.6 [0.0-17.1] | 12.9 [4.8-28.4] | 5.9 [0.0-22.9] | 0.4 [0.0-17.1] |
| CD24 (nMFI) | 11.4 [1.9-30.3] | 17.3 [7.6-40.0] | 14.1 [0.4-34.3] | 20.5 [8.3-41.4] | 13.9 [3.9-31.3] | 10.6 [0.0-23.3] |
| CD25 (nMFI) | 6.8 [0.0-21.2] | 19.1 [6.5-55.6] | 11.6 [0.8-45.4] | 12.1 [5.4-25.0] | 8.4 [0.0-25.3] | 10.8 [0.0-39.5] |
| CD29 (nMFI) | 39.9 [12.8-130.5] | 111.8 [35.6-193.0] | 53.2 [26.6-92.9] | 90.2 [31.9-133.9] | 66.0 [25.2-106.5] | 64.4 [18.0-128.3] |
| CD31 (nMFI) | 17.2 [7.0-44.4] | 37.6 [13.3-73.2] | 11.5 [2.1-33.8] | 26.8 [13.5-62.0] | 15.5 [3.6-32.4] | 19.6 [9.6-45.6] |
| CD40 (nMFI) | 12.6 [3.9-34.5] | 35.4 [20.2-82.6] | 20.7 [1.3-40.4] | 26.8 [10.6-53.5] | 17.2 [1.9-42.7] | 24.0 [3.5-61.4] |
| CD41b (nMFI) | 27.8 [13.1-52.5] | 51.9 [11.3-115-1] | 23.6 [12.4-48.7] | 48.1 [27.0-82.3] | 20.0 [10.1-51.9] | 34.9 [16.2-59.1] |
| CD42a (nMFI) | 34.2 [14.6-75.8] | 116.0 [48.8-241.7] | 29.0 [15.0-86.1] | 79.8 [38.2-146.7] | 36.1 [13.1-61.9] | 60.3 [24.1-102.1] |
| CD44 (nMFI) | 11.2 [3.8-21.7] | 21.1 [11.5-39.8] | 17.1 [3.5-34.0] | 10.1 [4.1-28.2] | 18.4 [4.6-37.4] | 17.4 [4.5-52.2] |
| CD45 (nMFI) | 2.9 [0.0-15.5] | 11.5 [0.0-49.6] | 12.3 [0.0-23.6] | 11.3 [3.7-30.3] | 11.7 [4.3-58.4] | 17.4 [3.2-40.5] |
| CD49e (nMFI) | 3.8 [0.0-15.7] | 8.9 [0.0-40.8] | 18.6 [5.8-35.7] | 10.9 [3.4-22.8] | 6.4 [1.3-30.9] | 6.5 [0.0-36.5] |
| CD56 (nMFI) | 0.0 [0.0-5.2] | 45.3 [12.7-91.4] | 14.1 [0.0-58.0] | 0.0 [0.0-12.1] | 0.0 [0.0-15.9] | 0.0 [0.0-22.2] |
| CD62P (nMFI) | 65.9 [32.7-136.4] | 112.9 [54.1-226.6] | 83.2 [37.2-154.2] | 86.8 [42.7-139.9] | 130.5 [47.2-314.0] | 126.0 [52.1-360.8] |

| | | | | | | |
|---|---|---|---|---|---|---|
| CD63 (nMFI) | 88.2 [67.0-131.4] | 63.2 [47.0-94.0] | 77.0 [64.7-104.2] | 92.3 [70.8-129.9] | 73.3 [49.6-119.8] | 72.0 [46.8-97.5] |
| CD69 (nMFI) | 28.6 [12.0-55.4] | 36.6 [21.4-56.7] | 42.7 [19.4-57.2] | 45.9 [26.3-70.7] | 26.0 [6.4-66.9] | 16.5 [3.1-43.2] |
| CD81 (nMFI) | 122.8 [67.3-176.3] | 112.3 [76.0-148.5] | 148.3 [113.0-176.6] | 109.1 [57.2-161.6] | 153.1 [76.0-181.5] | 149.1 [107.4-192.8] |
| CD86 (nMFI) | 7.1 [0.4-16.2] | 8.1 [0.0-21.6] | 10.6 [1.7-21.8] | 10.8 [4.9-20.0] | 11.8 [0.3-26.9] | 10.6 [1.0-34.0] |
| CD105 (nMFI) | 7.9 [0.0-17.7] | 17.5 [4.8-33.9] | 11.9 [2.5-32.1] | 13.4 [5.8-29.7] | 10.8 [3.7-30.8] | 5.7 [0.0-21.5] |
| CD133/1 (nMFI) | 12.3 [3.5-28.5] | 15.8 [6.8-31.2] | 12.2 [1.7-37.4] | 15.8 [8.2-38.9] | 13.3 [4.1-38.5] | 6.6 [0.0-33.7] |
| CD142 (nMFI) | 7.0 [0.3-17.7] | 11.4 [0.6-28.5] | 13.5 [0.8-34.9] | 11.1 [3.4-29.6] | 15.0 [0.6-38.2] | 4.3 [0.0-15.5] |
| CD146 (nMFI) | 2.3 [0.0-6.2] | 4.9 [0.5-11.3] | 8.0 [0.0-16.5] | 5.8 [2.3-13.6] | 9.9 [2.5-22.4] | 2.2 [0.0-12.6] |
| CD209 (nMFI) | 4.8 [0.0-14.9] | 11.2 [0.7-40.2] | 12.0 [3.0-29.8] | 10.4 [2.7-24.8] | 14.4 [4.8-43.0] | 7.5 [0.0-28.7] |
| CD326 (nMFI) | 8.1 [1.2-41.2] | 12.0 [1.4-28.5] | 22.8 [8.9-45.7] | 19.8 [5.5-46.8] | 11.5 [3.9-52.7] | 8.1 [0.0-43.6] |
| HLA-I (nMFI) | 7.9 [0.0-25.0] | 19.3 [0.2-77.5] | 4.4 [0.0-22.7] | 23.8 [9.7-49.5] | 5.7 [0.0-34.3] | 1.9 [0.0-32.0] |
| HLA-II (nMFI) | 22.9 [14.3-36.9] | 37.9 [19.6-74.5] | 29.7 [19.0-55.4] | 24.4 [18.6-45-4] | 38.6 [24.3-59.1] | 38.3 [23.0-62.6] |
| MCSP (nMFI) | 7.3 [0.2-14.4] | 12.0 [3.9-24.1] | 14.6 [6.5-28.8] | 10.7 [5.2-23.3] | 5.5 [0.0-30.6] | 9.4 [0.1-33.0] |
| ROR1 (nMFI) | 20.1 [6.7-55.6] | 18.0 [9.6-38.2] | 35.7 [14.4-48.8] | 38.2 [17.8-62.4] | 27.7 [8.5-84.5] | 24.1 [6.8-49.7] |
| SSEA4 (nMFI) | 6.7 [0.0-13.4] | 0.0 [0.0-11.5] | 5.4 [0.0-18.0] | 11.3 [5.7-23.8] | 9.8 [0.0-21.4] | 4.2 [0.0-18.2] |

For each group, the table reports: the nanoparticle concentration (N°/mL) and diameter (nm) measured at nanoparticle tracking analysis (NTA), and the mean Median Fluorescence Intensity (MFI) of CD9, CD63, and CD81 and the MFI of all EV-surface markers normalized for the mean MFI of CD9, CD63, and CD81 (nMFI) measured at flow cytometry. Data are expressed as median and interquartile range.

Supplementary Table 5. Nanoparticle characterization and Normalized Median Fluorescence Intensity of all 37 EV-surface markers – Statistics

| Variable | HC vs. iRBD | HC vs. DeNovo | HC vs. Late PD | HC vs. Syn. | HC vs. Tau. | iRBD vs. DeNovo | iRBD vs. Late PD | iRBD vs. Syn. | iRBD vs. Tau. | DeNovo vs. Late PD | DeNovo vs. Syn. | DeNovo vs. Tau. | Late PD vs. Syn. | Late PD vs. Tau. | Syn. vs. Tau. |
|---|---|---|---|---|---|---|---|---|---|---|---|---|---|---|---|
| NTA All (N°/mL) *E10 | >0.999 | 0.111 | >0.999 | 0.207 | >0.999 | >0.999 | >0.999 | >0.999 | >0.999 | 0.429 | >0.999 | 0.257 | 0.650 | >0.999 | 0.386 |
| NTA 30-150nm (N°/mL) *E10 | >0.999 | 0.103 | >0.999 | 0.203 | >0.999 | >0.999 | >0.999 | >0.999 | >0.999 | 0.369 | >0.999 | 0.328 | 0.590 | >0.999 | 0.494 |
| NTA 151-500nm (N°/mL) *E10 | >0.999 | 0.308 | >0.999 | 0.813 | >0.999 | >0.999 | >0.999 | >0.999 | >0.999 | 0.780 | >0.999 | 0.203 | >0.999 | >0.999 | 0.506 |
| Diameter (nm) | >0.999 | 0.167 | >0.999 | 0.653 | >0.999 | 0.896 | >0.999 | >0.999 | >0.999 | 0.160 | >0.999 | >0.999 | 0.615 | >0.999 | >0.999 |
| Mean CD9-CD63-CD81 (MFI) | >0.999 | 0.005 | 0.007 | 0.969 | 0.000 | 0.193 | 0.003 | >0.999 | 0.030 | <0.0001 | >0.999 | >0.999 | 0.001 | <0.0001 | >0.999 |
| CD1c (nMFI) | 0.241 | >0.999 | 0.277 | 0.326 | 0.317 | >0.999 | >0.999 | >0.999 | >0.999 | >0.999 | >0.999 | >0.999 | >0.999 | >0.999 | >0.999 |
| CD2 (nMFI) | 0.000 | 0.007 | >0.999 | 0.971 | >0.999 | >0.999 | 0.034 | >0.999 | 0.204 | 0.273 | >0.999 | 0.780 | >0.999 | >0.999 | >0.999 |
| CD3 (nMFI) | 0.005 | 0.211 | 0.225 | 0.073 | 0.315 | >0.999 | >0.999 | >0.999 | >0.999 | >0.999 | >0.999 | >0.999 | >0.999 | >0.999 | >0.999 |
| CD4 (nMFI) | >0.999 | >0.999 | 0.290 | >0.999 | 0.183 | >0.999 | 0.683 | >0.999 | 0.391 | 0.068 | >0.999 | 0.042 | >0.999 | >0.999 | >0.999 |
| CD8 (nMFI) | <0.0001 | <0.0001 | >0.999 | 0.276 | 0.001 | >0.999 | 0.001 | >0.999 | >0.999 | 0.002 | >0.999 | >0.999 | >0.999 | 0.024 | >0.999 |
| CD9 (nMFI) | 0.002 | >0.999 | >0.999 | >0.999 | >0.999 | 0.000 | 0.044 | 0.003 | <0.0001 | 0.674 | >0.999 | >0.999 | >0.999 | 0.429 | >0.999 |
| CD11c (nMFI) | >0.999 | >0.999 | 0.352 | 0.741 | >0.999 | >0.999 | 0.496 | 0.800 | >0.999 | 0.494 | 0.684 | >0.999 | >0.999 | >0.999 | >0.999 |
| CD14 (nMFI) | >0.999 | >0.999 | >0.999 | >0.999 | >0.999 | >0.999 | 0.107 | >0.999 | >0.999 | >0.999 | >0.999 | >0.999 | >0.999 | >0.999 | >0.999 |
| CD19 (nMFI) | 0.061 | >0.999 | 0.008 | 0.371 | >0.999 | >0.999 | >0.999 | >0.999 | 0.884 | >0.999 | >0.999 | >0.999 | >0.999 | 0.336 | >0.999 |
| CD20 (nMFI) | 0.027 | >0.999 | 0.159 | >0.999 | >0.999 | 0.159 | >0.999 | 0.102 | 0.002 | 0.629 | >0.999 | >0.999 | 0.391 | 0.011 | >0.999 |
| CD24 (nMFI) | 0.431 | >0.999 | 0.055 | >0.999 | >0.999 | >0.999 | >0.999 | >0.999 | 0.129 | >0.999 | >0.999 | >0.999 | >0.999 | 0.019 | >0.999 |
| CD25 (nMFI) | 0.001 | 0.871 | 0.111 | >0.999 | >0.999 | >0.999 | >0.999 | 0.350 | 0.667 | >0.999 | >0.999 | >0.999 | >0.999 | >0.999 | >0.999 |
| CD29 (nMFI) | 0.003 | >0.999 | 0.239 | >0.999 | >0.999 | 0.052 | >0.999 | 0.655 | 0.314 | >0.999 | >0.999 | >0.999 | >0.999 | >0.999 | >0.999 |
| CD31 (nMFI) | 0.024 | >0.999 | 0.368 | >0.999 | >0.999 | 0.003 | >0.999 | 0.026 | 0.347 | 0.038 | >0.999 | >0.999 | 0.228 | >0.999 | >0.999 |
| CD40 (nMFI) | 0.000 | >0.999 | 0.030 | >0.999 | 0.976 | 0.033 | >0.999 | 0.056 | 0.540 | 0.901 | >0.999 | >0.999 | >0.999 | >0.999 | >0.999 |
| CD41b (nMFI) | 0.134 | >0.999 | 0.005 | >0.999 | >0.999 | 0.048 | >0.999 | 0.048 | >0.999 | 0.003 | >0.999 | >0.999 | 0.005 | 0.240 | >0.999 |
| CD42a (nMFI) | <0.0001 | >0.999 | 0.001 | >0.999 | 0.957 | <0.0001 | 0.907 | 0.001 | 0.035 | 0.021 | >0.999 | >0.999 | 0.088 | >0.999 | >0.999 |
| CD44 (nMFI) | 0.006 | >0.999 | >0.999 | >0.999 | 0.990 | >0.999 | 0.053 | >0.999 | >0.999 | >0.999 | >0.999 | >0.999 | >0.999 | >0.999 | >0.999 |
| CD45 (nMFI) | 0.230 | 0.443 | 0.003 | 0.110 | 0.004 | >0.999 | >0.999 | >0.999 | >0.999 | >0.999 | >0.999 | >0.999 | >0.999 | >0.999 | >0.999 |
| CD49e (nMFI) | 0.141 | 0.002 | 0.051 | >0.999 | >0.999 | >0.999 | >0.999 | >0.999 | >0.999 | >0.999 | >0.999 | 0.611 | >0.999 | >0.999 | >0.999 |
| CD56 (nMFI) | <0.0001 | 0.005 | >0.999 | >0.999 | 0.901 | 0.061 | <0.0001 | <0.0001 | <0.0001 | 0.226 | >0.999 | >0.999 | >0.999 | >0.999 | >0.999 |
| CD62P (nMFI) | 0.125 | >0.999 | >0.999 | 0.280 | 0.033 | >0.999 | 0.416 | >0.999 | >0.999 | >0.999 | >0.999 | 0.933 | 0.652 | 0.121 | >0.999 |

| | | | | | | | | | | | | | | | |
|---|---|---|---|---|---|---|---|---|---|---|---|---|---|---|---|
| CD63 (nMFI) | **0.001** | >0.999 | >0.999 | >0.999 | 0.076 | 0.723 | **<0.0001** | >0.999 | >0.999 | 0.481 | >0.999 | >0.999 | 0.266 | **0.004** | >0.999 |
| CD69 (nMFI) | >0.999 | >0.999 | 0.053 | >0.999 | 0.256 | >0.999 | >0.999 | >0.999 | **0.008** | >0.999 | >0.999 | **0.028** | 0.294 | **<0.0001** | >0.999 |
| CD81 (nMFI) | >0.999 | 0.549 | >0.999 | >0.999 | 0.118 | 0.230 | >0.999 | 0.977 | **0.047** | 0.136 | >0.999 | >0.999 | 0.727 | **0.020** | >0.999 |
| CD86 (nMFI) | >0.999 | >0.999 | 0.509 | >0.999 | >0.999 | >0.999 | >0.999 | >0.999 | >0.999 | >0.999 | >0.999 | >0.999 | >0.999 | >0.999 | >0.999 |
| CD105 (nMFI) | **0.004** | 0.568 | **0.010** | 0.706 | >0.999 | >0.999 | >0.999 | >0.999 | **0.024** | >0.999 | >0.999 | 0.884 | >0.999 | 0.058 | 0.99 |
| CD133/1 (nMFI) | >0.999 | >0.999 | 0.528 | >0.999 | >0.999 | >0.999 | >0.999 | >0.999 | 0.180 | >0.999 | >0.999 | >0.999 | >0.999 | **0.008** | 0.54 |
| CD142 (nMFI) | >0.999 | >0.999 | 0.428 | >0.999 | >0.999 | >0.999 | >0.999 | >0.999 | >0.999 | >0.999 | >0.999 | >0.999 | >0.999 | 0.219 | >0.999 |
| CD146 (nMFI) | 0.514 | 0.609 | **0.020** | **0.023** | >0.999 | >0.999 | >0.999 | >0.999 | >0.999 | >0.999 | >0.999 | >0.999 | >0.999 | 0.939 | 0.39 |
| CD209 (nMFI) | 0.122 | 0.389 | 0.069 | **0.034** | >0.999 | >0.999 | >0.999 | >0.999 | >0.999 | >0.999 | >0.999 | >0.999 | >0.999 | >0.999 | >0.999 |
| CD326 (nMFI) | >0.999 | 0.216 | 0.223 | >0.999 | >0.999 | 0.469 | 0.644 | >0.999 | >0.999 | >0.999 | >0.999 | 0.180 | >0.999 | 0.221 | >0.999 |
| HLA-I (nMFI) | 0.076 | >0.999 | **0.003** | >0.999 | >0.999 | 0.052 | >0.999 | >0.999 | 0.054 | **0.005** | >0.999 | >0.999 | 0.302 | **0.004** | >0.999 |
| HLA-II (nMFI) | **0.001** | 0.584 | 0.872 | **0.019** | **0.009** | >0.999 | 0.497 | >0.999 | >0.999 | >0.999 | >0.999 | >0.999 | 0.984 | >0.999 | >0.999 |
| MCSP (nMFI) | 0.184 | **0.034** | 0.108 | >0.999 | >0.999 | >0.999 | >0.999 | >0.999 | >0.999 | >0.999 | 0.572 | >0.999 | >0.999 | >0.999 | >0.999 |
| ROR1 (nMFI) | >0.999 | >0.999 | 0.072 | >0.999 | >0.999 | >0.999 | 0.067 | >0.999 | >0.999 | >0.999 | >0.999 | >0.999 | >0.999 | 0.857 | >0.999 |
| SSEA4 (nMFI) | 0.105 | >0.999 | **0.002** | >0.999 | >0.999 | >0.999 | **<0.0001** | 0.081 | 0.297 | **0.008** | >0.999 | >0.999 | 0.835 | **0.019** | >0.999 |

For each multiple comparison, the P-value is reported. P-values lower than 0.05 were considered significant and shown in bold.

Supplementary Figure 1.

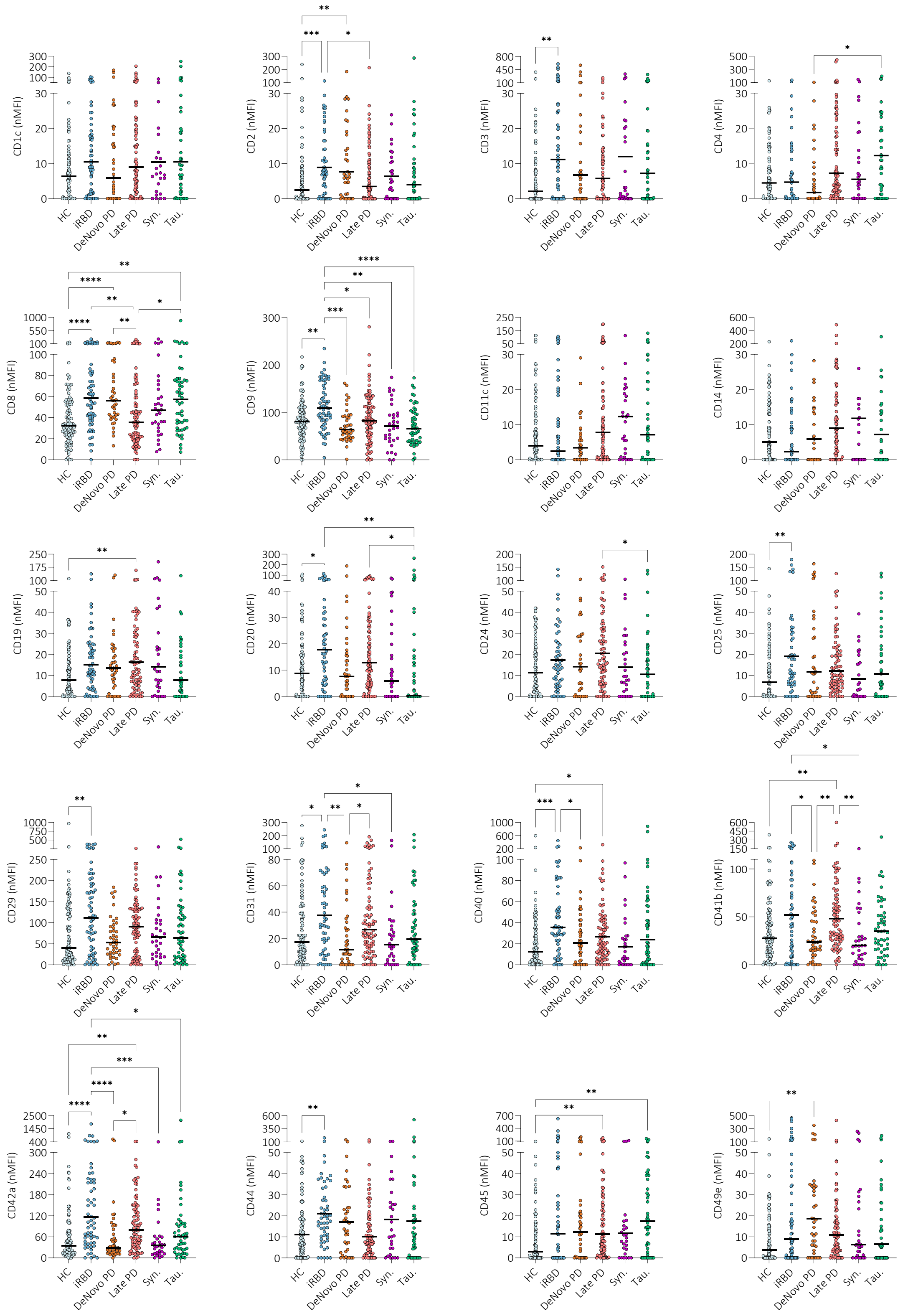

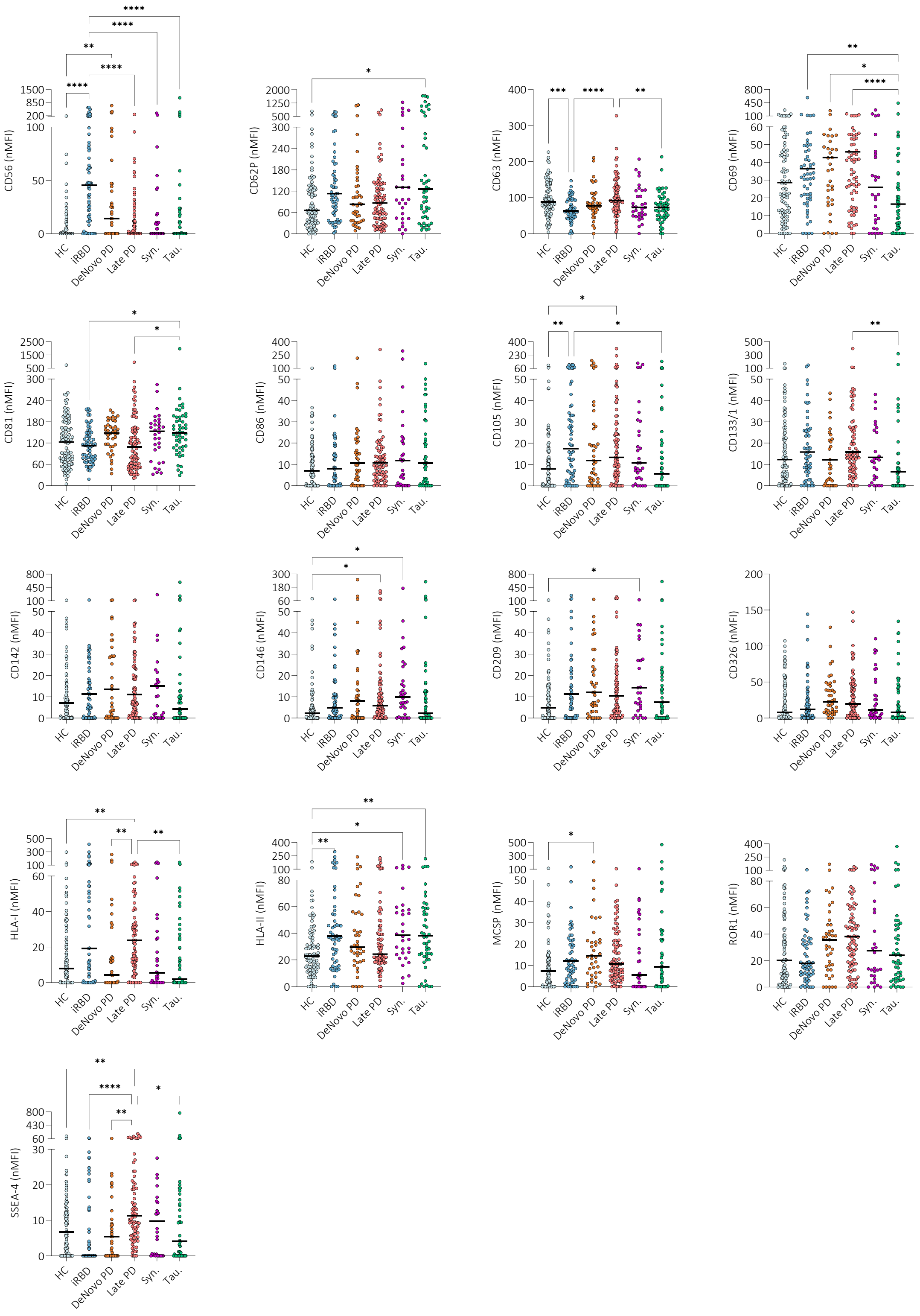


Normalized median fluorescence intensity (nMFI) of the 37 surface markers analyzed by flow cytometry. Each dot represents an individual patient, while the bar represents the median value of the group. (*P<0.05; **P<0.01; ***P<0.001, ****P<0.000). Data and statistics are reported in Supplementary Tables 4 and 5.

Supplementary Table 6. Demographic data and clinical scores of BioFIND cohort

| Variable | HC (n=26) | Late PD (n=51) | P-value |
|---|---|---|---|
| Age (y) | 71 ± 6 | 69 ± 5 | 0.208 |
| Sex (%M) | 62.0 | 65.0 | 0.785 |
| BMI | 26.6 [24.2-29.0] | 26.5 [23.4-29.4] | 0.956 |
| Disease Duration (y) | - | 7 ± 2 | - |
| Age of onset (y) | - | 62 ± 5 | - |
| H&Y | - | 2.0 [2.0-2.0] | - |
| MDS-UPDRS I | - | 12.0 [7.0-18.0] | - |
| MDS-UPDRS II | - | 10.0 [5.0-13.0] | - |
| MDS-UPDRS III | - | 39.0 [27.0-48.0] | - |
| MDS-UPDRS Total | - | 57.0 [45.0-77.0] | - |
| MoCA | - | 27.0 [26.0-29.0] | - |
| RBDSQ | - | 3.0 [1.0-6.0] | - |
| LEDD | - | 325.0 [200.0-457.3] | - |

For each group, demographic and clinical characteristics are reported as mean± standard deviation (SD), percentage (%), or median and [interquartile range]. BMI= Body Mass Index; H&Y = Hoehn and Yahr scale; MDS-UPDRS = Movement Disorder Society–Unified Parkinson's Disease Rating Scale; COMPASS 31 (OH= orthostatic hypotension, VM= vasomotor, SM= sudomotor, GI= gastrointestinal, BL = Bladder, PM= pupillomotor); BDI-II = Beck Depression Inventory II; MMSE = Mini-Mental State Examination; MoCA = Montreal Cognitive Assessment; RBDSQ = REM sleep Behavior Disorder Screening Questionnaire, LEDD = Levodopa Equivalent Daily Dose. For each comparison, the P-value is reported. P-values lower than 0.05 were considered significant and shown in bold.

Supplementary Table 7.

| Variable | HC (n=27) | Late PD (n=51) | P-value |
|---|---|---|---|
| EVs All (N°/mL) *E9 | 2.8 [1.2-3.5] | 2.1 [1.1-3.0] | 0.245 |
| EVs 30-150nm (N°/mL) *E9 | 0.9 [0.5-1.8] | 0.8 [0.4-1.3] | 0.598 |
| EVs 151-500nm (N°/mL) *E9 | 1.4 [0.9-2.2] | 1.3 [0.7-1.9] | 0.211 |
| Diameter (nm) | 203.1 [181.6-222.6] | 189.8 [181.5-209.5] | 0.204 |
| Mean CD9-CD63-CD81 (MFI) | 52.3 [44.6-62.5] | 58.9 [47.8-71.6] | 0.191 |
| CD1c (nMFI) | 0.0 [0.0-0.0] | 0.0 [0.0-0.4] | **0.019** |
| CD2 (nMFI) | 0.0 [0.0-0.0] | 0.0 [0.0-0.1] | 0.341 |
| CD3 (nMFI) | 0.0 [0.0-0.0] | 0.0 [0.0-0.0] | 0.290 |
| CD4 (nMFI) | 0.3 [0.1-0.8] | 0.4 [0.2-0.7] | 1.000 |
| CD8 (nMFI) | 1.4 [0.7-2.1] | 1.6 [0.9-2.3] | 0.714 |
| CD9 (nMFI) | 62.8 [56.9-67.2] | 61.2 [55.3-70.3] | 0.914 |
| CD11c (nMFI) | 0.0 [0.0-0.3] | 0.0 [0.0-0.3] | 0.565 |
| CD14 (nMFI) | 0.8 [0.2-2.0] | 0.8 [0.0-1.6] | 0.376 |
| CD19 (nMFI) | 0.0 [0.0-0.0] | 0.0 [0.0-0.2] | 0.065 |
| CD20 (nMFI) | 0.7 [0.3-0.9] | 0.6 [0.3-1.0] | 0.792 |
| CD24 (nMFI) | 1.3 [0.5-2.1] | 1.4 [0.8-2.0] | 0.658 |
| CD25 (nMFI) | 0.0 [0.0-0.4] | 0.0 [0.0-0.3] | 0.980 |
| CD29 (nMFI) | 0.9 [0.4-1.7] | 1.7 [0.8-2.6] | **0.031** |
| CD31 (nMFI) | 0.1 [0.0-0.5] | 0.1 [0.0-0.4] | 0.845 |
| CD40 (nMFI) | 0.4 [0.0-0.7] | 0.4 [0.0-0.6] | 0.485 |
| CD41b (nMFI) | 0.0 [0.0-0.1] | 0.0 [0.0-0.4] | 0.203 |
| CD42a (nMFI) | 0.0 [0.0-0.5] | 0.0 [0.0-0.4] | 0.759 |
| CD44 (nMFI) | 0.4 [0.0-0.9] | 0.4 [0.1-1.0] | 0.859 |
| CD45 (nMFI) | 0.0 [0.0-0.1] | 0.0 [0.0-0.0] | 0.520 |
| CD49e (nMFI) | 0.4 [0.0-0.9] | 0.2 [0.0-0.9] | 0.864 |
| CD56 (nMFI) | 0.0 [0.0-0.0] | 0.0 [0.0-0.0] | 0.488 |
| CD62P (nMFI) | 0.0 [0.0-0.2] | 0.0 [0.0-0.3] | 0.463 |
| CD63 (nMFI) | 38.9 [29.3-44.4] | 37.8 [33.4-43.7] | 0.674 |
| CD69 (nMFI) | 0.0 [0.0-0.1] | 0.0 [0.0-1.1] | **0.050** |
| CD81 (nMFI) | 196.9 [186.3-214.8] | 199.3 [185.1-209.3] | 0.636 |
| CD86 (nMFI) | 0.0 [0.0-0.6] | 0.4 [0.0-1.1] | 0.432 |
| CD105 (nMFI) | 0.0 [0.0-0.9] | 0.5 [0.0-1.0] | 0.104 |
| CD133/1 (nMFI) | 4.4 [3.1-7.2] | 6.1 [4.1-8.8] | 0.122 |
| CD142 (nMFI) | 0.2 [0.0-0.5] | 0.3 [0.0-0.8] | 0.585 |
| CD146 (nMFI) | 0.3 [0.0-0.8] | 0.1 [0.0-0.5] | 0.088 |
| CD209 (nMFI) | 0.0 [0.0-0.2] | 0.1 [0.0-0.3] | 0.407 |
| CD326 (nMFI) | 0.4 [0.0-0.7] | 1.1 [0.1-2.3] | **0.007** |
| HLA-I (nMFI) | 0.0 [0.0-0.0] | 0.0 [0.0-0.0] | 0.280 |
| HLA-II (nMFI) | 3.2 [2.1-4.4] | 3.8 [2.6-5.3] | 0.210 |
| MCSP (nMFI) | 0.1 [0.0-0.5] | 0.4 [0.0-0.7] | 0.228 |
| ROR1 (nMFI) | 0.5 [0.0-0.9] | 0.7 [0.0-2.0] | 0.108 |
| SSEA4 (nMFI) | 1.0 [0.3-2.0] | 1.3 [0.6-2.0] | 0.504 |

For the two groups are reported the nanoparticle concentration (N°/mL) and diameter (nm) measured at nanoparticle tracking analysis (NTA); the mean Median Fluorescence Intensity (MFI) of CD9, CD63, and CD81 measured at flow cytometry; the MFI of all EV-surface markers normalized for the mean MFI of CD9, CD63, and CD81 (nMFI). Data are expressed as median and interquartile range. P-value lower than 0.05 were considered significant and shown in bold.

Supplementary Table 8. Demographic data, medical history, and clinical scores of training and validation cohorts

| | HC | | | iRBD | | | DeNovo PD | | |
|---|---|---|---|---|---|---|---|---|---|
| | Training (n=67) | Validation (n=33) | P-value | Training (n=43) | Validation (n=21) | P-value | Training (n=27) | Validation (n=14) | P-value |
| Age (y) | 65±10 | 65±12 | 0.692 | 68±7 | 67±7 | 0.562 | 62±9 | 67±12 | 0.067 |
| Sex (%M) | 47.7 | 57.5 | 0.239 | 83.7 | 85.7 | 0.575 | 51.8 | 64.2 | 0.550 |
| BMI | 24.6 [22.5-27.6] | 25.9 [23.2-28.9] | 0.324 | 26.6 [24.2-29.9] | 27.0 [24.4-28.4] | 0.755 | 24.8 [22.9-28.8] | 26.0 [22.7-29.3] | 0.336 |
| Disease Duration (y) | - | - | - | 7±6 | 5±4 | 0.316 | 2±1 | 2±1 | 0.200 |
| Age of onset (y) | - | - | - | 62±8 | 62±7 | 0.956 | 60±9 | 65±12 | 0.092 |
| H&Y | - | - | - | - | - | - | 1.0 [1.0-2.0] | 1.0 [1.0-2.0] | 0.839 |
| MDS-UPDRS III | - | - | | 4.0 [2.0-6.0] | 5.0 [2.3-8.0] | 0.332 | 16.5 [9.5-23.5] | 12.5 [7.8-25.0] | 0.474 |
| MMSE | - | - | - | 29.0 [28.0-30.0] | 29.0 [28.0-30.0] | 0.642 | 30 [28.8-30.0] | 30.0 [28.0-30.0] | 0.952 |
| MoCA | - | - | - | 26.0 [22.0-28.0] | 26.0 [25.0-28.0] | 0.134 | 26.0 [23.0-28.0] | 27.5 [20.8-29.3] | 0.675 |
| LEDD | - | - | - | - | - | - | 100.0 [0.0-187.1] | 100.0 [37.5-127.5] | 0.944 |
| | **Late PD** | | | **Synucleinopathy** | | | **Tauopathy** | | |
| | Training (n=60) | Validation (n=29) | P-value | Training (n=21) | Validation (n=11) | P-value | Training (n=35) | Validation (n=17) | P-value |
| Age (y) | 69±7 | 67±9 | 0.322 | 70±8 | 67±11 | 0.531 | 71 14 | 73 8 | 0.868 |
| Sex (%M) | 68.3 | 62.1 | 0.635 | 57.1 | 54.5 | 0.590 | 48.5 | 82.4 | **0.034** |
| BMI | 25.7 [23.3-29.3] | 25.3 [22.8-28.9] | 0.647 | 25.8 [22.9-28.9] | 22.9 [21.6-24.9] | 0.076 | 24.8 [22.6-28.4] | 27.4 [24.1-30.7] | 0.102 |
| Disease Duration (y) | 7±4 | 7±2 | 0.867 | 4±2 | 4±3 | 0.755 | 4 2 | 3 2 | 0.350 |
| Age of onset (y) | 62±8 | 60±10 | 0.315 | 65±10 | 65±12 | 0.696 | 69 7 | 69 8 | 0.710 |
| H&Y | 2.0 [2.0-2.0] | 2.0 [1.5-3.0] | 0.780 | 3.5 [3.0-4.0] | 3.0 [3.0-5.0] | 0.729 | 4.0 [3.0-4.0] | 3.0 [2.5-4.0] | 0.526 |
| MDS-UPDRS III | 28.5 [17.3-44.0] | 29.0 [20.5-47.5] | 0.697 | 29.5 [19.5-40.5] | 34.0 [26.0-52.0] | 0.297 | 34.0 [25.5-41.5] | 29.0 [19.0-40.0] | 0.225 |
| MMSE | 29.0 [ 28.0-30.0] | 29.0 [29.0-30.0] | 0.238 | 28.0 [25.5-29.0] | 27.0 [25.0-29.0] | 0.820 | 26.0 [23.8-28.0] | 27.0 [23.5-29.0] | 0.561 |
| MoCA | 27.0 [24.0-29.0] | 28.0 [27.0-29.0] | 0.084 | 23.5 [18.8-26.0] | 22.0 [18.0-26.0] | 0.769 | 20.0 [16.5-24.52] | 21.0 [15.5-24.5] | 0.916 |
| LEDD | 306.5 [200.0-556.3] | 525.0 [363.8-597.0] | **0.031** | 300.0 [0.0-500.0] | 272.0 [0.0-518.8] | 0.735 | 525.0 [27.3-750.0] | 475.0 [381.8-660.0] | 0.545 |

Within each group, subjects in the training cohort were compared to those in the validation cohort. Demographic, medical, and clinical characteristics are reported as mean±standard deviation (SD), percentage (%), or median and [interquartile range]. BMI= Body Mass Index; H&Y = Hoehn and Yahr scale; MDS-UPDRS = Movement Disorder Society–Unified Parkinson's Disease Rating Scale; MMSE = Mini-Mental State Examination; MoCA = Montreal Cognitive Assessment; LEDD = Levodopa Equivalent Daily Dose. For comparison, the P-value is reported. P-values lower than 0.05 were considered significant and shown in bold.
