## Supplementary material for "Plasmatic immune extracellular vesicle profiles identify prodromal and early stages of Parkinson’s disease": Supplem.File2

### Supplementary File 2

Tables 1 and 2 show correlation matrices of EV antigens measured in plasma and CSF. For each correlation the Pearson's R value (R) and *P* -value (P) are reported.

Tables 3 to 7 show model ID, algorithm of oversampling (when applicable; SMOTE [Synthetic Minority Over-Sampling Technique], SMOTENN [SMOTE and Nearest Neighbors], or RO [Random Oversampling]), number of trees in the random forest regressors, maximum number of split for each tree (number of leaves), and their performance at validation: sensitivity, specificity, accuracy and macro-average accuracy (crude mean of sensitivity and specificity). The model with the highest performance for each discrimination (HC vs iRBD; HC vs DeNovo PD; HC vs Late PD; HC vs. Synucleinopathy; HC vs. Tauopathy ) expressed as macro-average accuracy was selected, highlighted in green and displayed in Figure 5.

| | | CD1c | CD2 | CD3 | CD4 | CD8 | CD9 | CD11c | CD14 | CD19 | CD20 | CD24 | CD25 | CD29 | CD31 | CD40 | CD41b | CD42a | CD44 | CD45 | CD49e | CD56 | CD62P | CD63 | CD69 | CD81 | CD86 | CD105 | CD133/1 | CD142 | CD146 | CD209 | CD326 | HLA-I | HLA-II | MCSP | ROR1 | SSEA4 |
|---|---|---|---|---|---|---|---|---|---|---|---|---|---|---|---|---|---|---|---|---|---|---|---|---|---|---|---|---|---|---|---|---|---|---|---|---|---|---|
| **CD1c** | **R** | 0.203 | -0.056 | -0.113 | -0.237 | -0.298 | -0.162 | -0.270 | -0.212 | -0.089 | -0.103 | -0.212 | -0.058 | -0.129 | 0.055 | -0.169 | 0.103 | -0.062 | -0.155 | -0.057 | -0.015 | -0.133 | 0.047 | -0.182 | -0.119 | 0.088 | 0.266 | 0.204 | -0.086 | -0.172 | 0.131 | -0.177 | 0.123 | -0.129 | -0.231 | -0.154 | -0.209 | -0.077 |
| | **P** | 0.320 | 0.787 | 0.582 | 0.243 | 0.139 | 0.431 | 0.182 | 0.298 | 0.665 | 0.618 | 0.298 | 0.777 | 0.530 | 0.789 | 0.409 | 0.617 | 0.763 | 0.448 | 0.781 | 0.942 | 0.517 | 0.821 | 0.374 | 0.563 | 0.669 | 0.190 | 0.317 | 0.677 | 0.401 | 0.525 | 0.388 | 0.549 | 0.529 | 0.257 | 0.453 | 0.306 | 0.709 |
| **CD2** | **R** | 0.794 | 0.204 | -0.102 | -0.177 | -0.355 | 0.022 | -0.177 | -0.165 | -0.027 | 0.174 | -0.288 | -0.135 | 0.027 | 0.521 | -0.020 | 0.635 | -0.135 | -0.070 | 0.230 | 0.043 | -0.062 | 0.166 | -0.110 | -0.128 | -0.102 | -0.128 | -0.068 | -0.085 | -0.011 | -0.001 | -0.070 | 0.629 | -0.070 | -0.245 | -0.152 | 0.012 | 0.097 |
| | **P** | **0.000** | 0.318 | 0.620 | 0.388 | 0.075 | 0.917 | 0.386 | 0.421 | 0.897 | 0.395 | 0.154 | 0.512 | 0.896 | **0.006** | 0.924 | **0.000** | 0.511 | 0.735 | 0.258 | 0.836 | 0.763 | 0.418 | 0.594 | 0.533 | 0.619 | 0.532 | 0.743 | 0.679 | 0.956 | 0.996 | 0.735 | **0.001** | 0.736 | 0.228 | 0.458 | 0.955 | 0.638 |
| **CD3** | **R** | 0.706 | 0.126 | -0.151 | -0.194 | -0.438 | 0.114 | -0.162 | -0.071 | -0.045 | 0.020 | -0.262 | -0.278 | -0.152 | 0.592 | 0.016 | 0.534 | -0.240 | -0.083 | 0.192 | -0.022 | -0.135 | 0.353 | -0.123 | -0.275 | -0.266 | -0.186 | -0.086 | -0.066 | -0.055 | -0.087 | 0.012 | 0.585 | -0.075 | -0.266 | -0.087 | 0.044 | 0.044 |
| | **P** | **0.000** | 0.539 | 0.462 | 0.343 | **0.025** | 0.578 | 0.430 | 0.731 | 0.828 | 0.922 | 0.195 | 0.169 | 0.459 | **0.001** | 0.938 | **0.005** | 0.238 | 0.686 | 0.347 | 0.915 | 0.509 | 0.077 | 0.549 | 0.174 | 0.190 | 0.364 | 0.677 | 0.750 | 0.790 | 0.674 | 0.953 | **0.002** | 0.717 | 0.189 | 0.673 | 0.830 | 0.829 |
| **CD4** | **R** | 0.000 | -0.152 | -0.031 | -0.224 | -0.210 | -0.162 | -0.165 | -0.116 | -0.126 | -0.119 | -0.117 | -0.027 | -0.087 | -0.021 | -0.145 | -0.022 | -0.059 | -0.134 | -0.076 | -0.030 | -0.156 | 0.116 | -0.128 | -0.059 | 0.052 | 0.264 | 0.259 | -0.084 | -0.077 | 0.139 | -0.115 | -0.011 | -0.119 | -0.099 | -0.192 | -0.111 | -0.101 |
| | **P** | 0.999 | 0.459 | 0.882 | 0.272 | 0.304 | 0.429 | 0.421 | 0.573 | 0.541 | 0.563 | 0.568 | 0.894 | 0.673 | 0.918 | 0.479 | 0.913 | 0.776 | 0.514 | 0.714 | 0.886 | 0.446 | 0.571 | 0.535 | 0.773 | 0.802 | 0.192 | 0.201 | 0.683 | 0.708 | 0.499 | 0.575 | 0.956 | 0.563 | 0.630 | 0.349 | 0.590 | 0.623 |
| **CD8** | **R** | 0.143 | 0.005 | -0.223 | -0.178 | -0.273 | -0.032 | -0.236 | -0.165 | -0.075 | -0.203 | -0.030 | 0.062 | -0.104 | 0.310 | -0.039 | 0.101 | 0.142 | -0.036 | 0.158 | 0.124 | -0.092 | 0.089 | -0.054 | -0.154 | -0.024 | 0.130 | 0.079 | -0.143 | 0.154 | 0.284 | -0.207 | 0.015 | -0.069 | -0.078 | -0.334 | -0.365 | -0.055 |
| | **P** | 0.485 | 0.979 | 0.273 | 0.384 | 0.177 | 0.876 | 0.245 | 0.421 | 0.715 | 0.321 | 0.883 | 0.765 | 0.613 | 0.124 | 0.851 | 0.623 | 0.489 | 0.860 | 0.442 | 0.545 | 0.654 | 0.666 | 0.792 | 0.453 | 0.907 | 0.527 | 0.702 | 0.485 | 0.452 | 0.160 | 0.311 | 0.944 | 0.736 | 0.705 | 0.096 | 0.067 | 0.789 |
| **CD9** | **R** | 0.550 | -0.004 | 0.226 | -0.109 | -0.306 | 0.147 | 0.034 | -0.117 | -0.108 | 0.183 | -0.149 | 0.054 | 0.217 | 0.359 | 0.072 | **0.467** | -0.039 | 0.080 | 0.196 | 0.045 | -0.234 | 0.194 | -0.117 | -0.051 | -0.204 | -0.170 | -0.038 | -0.131 | 0.284 | 0.048 | 0.011 | 0.352 | 0.268 | 0.054 | -0.395 | 0.306 | 0.284 |
| | **P** | **0.004** | 0.985 | 0.266 | 0.597 | 0.129 | 0.475 | 0.871 | 0.569 | 0.598 | 0.372 | 0.468 | 0.792 | 0.286 | 0.072 | 0.726 | 0.016 | 0.848 | 0.699 | 0.337 | 0.826 | 0.250 | 0.342 | 0.571 | 0.806 | 0.317 | 0.407 | 0.856 | 0.524 | 0.160 | 0.814 | 0.959 | 0.078 | 0.185 | 0.793 | **0.046** | 0.129 | 0.160 |
| **CD11c** | **R** | 0.774 | 0.182 | -0.131 | -0.247 | -0.388 | 0.006 | -0.204 | -0.227 | -0.060 | 0.087 | -0.305 | -0.090 | 0.051 | 0.479 | -0.059 | 0.634 | -0.140 | -0.126 | 0.171 | 0.015 | -0.117 | 0.136 | -0.185 | -0.065 | -0.075 | -0.065 | -0.092 | -0.103 | -0.075 | -0.059 | -0.117 | 0.566 | -0.096 | -0.283 | -0.164 | -0.019 | 0.065 |
| | **P** | **0.000** | 0.373 | 0.525 | 0.224 | **0.050** | 0.977 | 0.317 | 0.264 | 0.769 | 0.674 | 0.130 | 0.661 | 0.804 | **0.013** | 0.775 | **0.001** | 0.496 | 0.538 | 0.403 | 0.942 | 0.568 | 0.508 | 0.365 | 0.754 | 0.714 | 0.753 | 0.655 | 0.617 | 0.716 | 0.776 | 0.568 | **0.003** | 0.639 | 0.161 | 0.424 | 0.927 | 0.752 |
| **CD14** | **R** | -0.187 | -0.209 | -0.137 | -0.095 | -0.193 | -0.130 | -0.121 | -0.019 | -0.059 | -0.161 | -0.032 | -0.067 | -0.197 | -0.081 | -0.067 | -0.238 | -0.057 | -0.060 | -0.129 | -0.021 | -0.122 | 0.078 | -0.062 | -0.103 | 0.049 | 0.278 | 0.309 | -0.025 | -0.139 | 0.304 | -0.093 | -0.131 | -0.131 | -0.075 | -0.080 | -0.145 | -0.140 |
| | **P** | 0.362 | 0.305 | 0.505 | 0.643 | 0.344 | 0.528 | 0.556 | 0.926 | 0.775 | 0.432 | 0.876 | 0.747 | 0.335 | 0.693 | 0.743 | 0.242 | 0.781 | 0.770 | 0.531 | 0.920 | 0.552 | 0.704 | 0.765 | 0.616 | 0.811 | 0.169 | 0.124 | 0.902 | 0.500 | 0.131 | 0.650 | 0.524 | 0.525 | 0.717 | 0.696 | 0.479 | 0.495 |
| **CD19** | **R** | -0.140 | -0.247 | -0.236 | -0.061 | -0.092 | 0.073 | -0.175 | 0.144 | -0.116 | -0.219 | 0.063 | -0.120 | -0.020 | -0.133 | -0.085 | -0.137 | -0.321 | -0.179 | -0.184 | -0.199 | -0.190 | 0.431 | 0.149 | 0.048 | -0.277 | -0.040 | 0.134 | -0.149 | -0.245 | -0.178 | -0.155 | 0.249 | -0.191 | -0.319 | -0.110 | 0.517 | -0.279 |
| | **P** | 0.495 | 0.224 | 0.246 | 0.767 | 0.653 | 0.724 | 0.393 | 0.482 | 0.572 | 0.283 | 0.761 | 0.559 | 0.921 | 0.517 | 0.681 | 0.505 | 0.110 | 0.382 | 0.367 | 0.331 | 0.354 | **0.028** | 0.466 | 0.817 | 0.171 | 0.847 | 0.514 | 0.468 | 0.228 | 0.385 | 0.449 | 0.220 | 0.350 | 0.112 | 0.591 | **0.007** | 0.167 |
| **CD20** | **R** | 0.234 | -0.030 | -0.181 | -0.165 | -0.139 | 0.058 | -0.204 | 0.075 | -0.154 | -0.071 | -0.036 | -0.135 | 0.028 | 0.071 | -0.094 | 0.224 | -0.265 | -0.228 | 0.023 | -0.178 | -0.210 | 0.455 | -0.208 | -0.023 | -0.280 | -0.185 | -0.003 | -0.123 | -0.196 | -0.256 | -0.138 | 0.523 | -0.175 | -0.361 | -0.164 | 0.512 | -0.186 |
| | **P** | 0.250 | 0.883 | 0.376 | 0.421 | 0.498 | 0.778 | 0.318 | 0.715 | 0.451 | 0.731 | 0.861 | 0.512 | 0.894 | 0.730 | 0.649 | 0.271 | 0.190 | 0.262 | 0.913 | 0.385 | 0.302 | **0.019** | 0.308 | 0.912 | 0.166 | 0.366 | 0.989 | 0.549 | 0.337 | 0.206 | 0.502 | **0.006** | 0.393 | 0.070 | 0.425 | **0.007** | 0.363 |
| **CD24** | **R** | -0.043 | -0.119 | -0.263 | -0.021 | -0.036 | 0.066 | -0.146 | 0.110 | -0.144 | -0.153 | 0.021 | 0.048 | 0.159 | -0.081 | -0.157 | 0.111 | -0.242 | -0.158 | -0.051 | -0.163 | -0.269 | 0.445 | -0.223 | 0.236 | -0.280 | -0.091 | 0.063 | -0.171 | -0.137 | -0.167 | -0.144 | 0.277 | -0.152 | -0.263 | -0.205 | 0.611 | -0.127 |
| | **P** | 0.836 | 0.564 | 0.194 | 0.918 | 0.862 | 0.747 | 0.477 | 0.594 | 0.484 | 0.456 | 0.920 | 0.817 | 0.438 | 0.693 | 0.444 | 0.590 | 0.233 | 0.440 | 0.803 | 0.427 | 0.184 | **0.023** | 0.273 | 0.246 | 0.165 | 0.658 | 0.762 | 0.404 | 0.506 | 0.415 | 0.483 | 0.171 | 0.459 | 0.194 | 0.315 | **0.001** | 0.535 |
| **CD25** | **R** | -0.275 | -0.155 | -0.246 | -0.038 | -0.050 | 0.006 | -0.136 | 0.169 | -0.145 | -0.185 | 0.257 | -0.005 | 0.081 | -0.097 | 0.109 | -0.125 | -0.250 | -0.100 | 0.008 | -0.133 | -0.216 | 0.439 | -0.115 | 0.219 | -0.235 | -0.090 | 0.119 | -0.119 | -0.105 | -0.010 | -0.118 | 0.137 | -0.182 | -0.294 | -0.113 | 0.552 | -0.274 |
| | **P** | 0.175 | 0.451 | 0.225 | 0.855 | 0.810 | 0.975 | 0.508 | 0.409 | 0.479 | 0.367 | 0.206 | 0.982 | 0.693 | 0.639 | 0.596 | 0.544 | 0.218 | 0.628 | 0.970 | 0.516 | 0.288 | **0.025** | 0.575 | 0.283 | 0.247 | 0.660 | 0.563 | 0.563 | 0.610 | 0.963 | 0.565 | 0.504 | 0.372 | 0.144 | 0.581 | **0.003** | 0.176 |
| **CD29** | **R** | 0.853 | 0.228 | 0.008 | -0.177 | -0.358 | 0.031 | -0.106 | -0.228 | 0.015 | 0.190 | -0.263 | -0.085 | 0.119 | 0.482 | -0.034 | 0.638 | -0.150 | 0.004 | 0.183 | 0.083 | -0.169 | 0.064 | -0.117 | -0.067 | -0.055 | -0.089 | -0.034 | -0.089 | -0.025 | -0.090 | 0.011 | 0.650 | 0.039 | -0.240 | -0.114 | 0.101 | 0.223 |
| | **P** | **0.000** | 0.262 | 0.970 | 0.388 | 0.072 | 0.880 | 0.605 | 0.262 | 0.941 | 0.353 | 0.194 | 0.681 | 0.563 | **0.013** | 0.868 | **0.000** | 0.464 | 0.986 | 0.370 | 0.688 | 0.409 | 0.756 | 0.569 | 0.744 | 0.788 | 0.667 | 0.870 | 0.667 | 0.902 | 0.662 | 0.957 | **0.000** | 0.850 | 0.238 | 0.580 | 0.625 | 0.273 |
| **CD31** | **R** | 0.026 | -0.172 | -0.070 | -0.171 | -0.242 | 0.079 | -0.093 | 0.025 | -0.134 | -0.331 | -0.046 | 0.004 | 0.085 | -0.099 | -0.123 | 0.222 | -0.181 | -0.112 | -0.067 | -0.155 | -0.312 | 0.288 | -0.205 | 0.055 | -0.206 | -0.163 | -0.004 | -0.265 | -0.064 | -0.215 | -0.083 | 0.090 | 0.048 | -0.239 | -0.318 | 0.290 | -0.124 |
| | **P** | 0.901 | 0.401 | 0.732 | 0.404 | 0.233 | 0.703 | 0.652 | 0.904 | 0.514 | 0.099 | 0.824 | 0.985 | 0.680 | 0.630 | 0.549 | 0.276 | 0.375 | 0.586 | 0.744 | 0.450 | 0.121 | 0.153 | 0.316 | 0.789 | 0.312 | 0.426 | 0.986 | 0.190 | 0.756 | 0.291 | 0.686 | 0.660 | 0.816 | 0.240 | 0.114 | 0.151 | 0.546 |
| **CD40** | **R** | 0.012 | -0.235 | 0.046 | -0.198 | -0.184 | -0.043 | -0.016 | -0.006 | -0.187 | -0.275 | -0.015 | 0.060 | 0.145 | -0.208 | -0.014 | 0.215 | -0.055 | -0.031 | -0.101 | -0.162 | -0.221 | 0.257 | -0.169 | 0.020 | -0.105 | -0.055 | 0.002 | -0.222 | 0.078 | -0.220 | -0.062 | -0.061 | 0.344 | -0.145 | -0.286 | 0.273 | 0.066 |
| | **P** | 0.952 | 0.249 | 0.824 | 0.333 | 0.369 | 0.833 | 0.938 | 0.978 | 0.360 | 0.174 | 0.943 | 0.771 | 0.479 | 0.308 | 0.946 | 0.291 | 0.790 | 0.880 | 0.622 | 0.429 | 0.277 | 0.204 | 0.410 | 0.922 | 0.609 | 0.790 | 0.994 | 0.275 | 0.703 | 0.280 | 0.762 | 0.768 | 0.085 | 0.479 | 0.157 | 0.178 | 0.749 |
| **CD41b** | **R** | 0.603 | 0.043 | 0.232 | -0.268 | -0.442 | 0.001 | 0.169 | -0.225 | -0.148 | -0.030 | -0.184 | 0.016 | 0.078 | 0.240 | 0.028 | 0.482 | -0.004 | 0.026 | 0.146 | -0.048 | -0.280 | 0.124 | -0.107 | -0.102 | -0.066 | -0.092 | -0.148 | -0.218 | 0.167 | -0.166 | 0.134 | 0.171 | 0.435 | 0.020 | -0.354 | 0.083 | 0.378 |
| | **P** | **0.001** | 0.834 | 0.255 | 0.185 | **0.024** | 0.995 | 0.408 | 0.269 | 0.469 | 0.885 | 0.367 | 0.937 | 0.706 | 0.237 | 0.891 | **0.013** | 0.985 | 0.901 | 0.476 | 0.815 | 0.166 | 0.546 | 0.601 | 0.620 | 0.748 | 0.655 | 0.469 | 0.285 | 0.416 | 0.417 | 0.513 | 0.403 | **0.027** | 0.924 | 0.076 | 0.685 | 0.057 |
| **CD42a** | **R** | -0.253 | -0.010 | -0.052 | 0.119 | -0.130 | 0.374 | -0.260 | 0.033 | 0.099 | -0.145 | -0.189 | -0.230 | -0.349 | 0.190 | -0.290 | -0.047 | -0.259 | 0.000 | -0.091 | -0.045 | -0.247 | 0.267 | -0.054 | -0.306 | -0.401 | -0.197 | -0.230 | 0.070 | -0.236 | -0.104 | 0.147 | -0.022 | -0.270 | -0.223 | -0.148 | 0.009 | -0.121 |
| | **P** | 0.213 | 0.960 | 0.801 | 0.562 | 0.526 | 0.060 | 0.200 | 0.871 | 0.632 | 0.480 | 0.356 | 0.259 | 0.081 | 0.353 | 0.150 | 0.819 | 0.202 | 0.998 | 0.660 | 0.827 | 0.224 | 0.187 | 0.793 | 0.128 | **0.042** | 0.335 | 0.258 | 0.734 | 0.245 | 0.612 | 0.473 | 0.916 | 0.183 | 0.273 | 0.471 | 0.966 | 0.555 |
| **CD44** | **R** | 0.081 | -0.121 | -0.215 | -0.049 | -0.158 | 0.044 | -0.262 | 0.107 | -0.127 | -0.075 | 0.008 | -0.105 | 0.000 | 0.142 | 0.086 | 0.126 | -0.291 | -0.150 | -0.001 | -0.129 | -0.180 | 0.606 | -0.139 | 0.017 | -0.349 | -0.279 | -0.078 | -0.231 | -0.160 | -0.187 | -0.164 | 0.440 | -0.139 | -0.382 | -0.154 | 0.600 | -0.223 |
| | **P** | 0.695 | 0.557 | 0.291 | 0.812 | 0.442 | 0.832 | 0.196 | 0.601 | 0.537 | 0.715 | 0.970 | 0.611 | 1.000 | 0.489 | 0.676 | 0.540 | 0.149 | 0.466 | 0.997 | 0.529 | 0.379 | **0.001** | 0.499 | 0.932 | 0.080 | 0.168 | 0.703 | 0.256 | 0.436 | 0.359 | 0.424 | **0.024** | 0.497 | 0.054 | 0.453 | **0.001** | 0.273 |
| **CD45** | **R** | 0.337 | -0.071 | -0.226 | -0.235 | -0.531 | 0.051 | -0.326 | -0.122 | -0.148 | -0.155 | -0.204 | -0.104 | -0.190 | 0.379 | -0.025 | 0.274 | -0.151 | -0.111 | 0.132 | -0.023 | -0.218 | 0.325 | -0.188 | -0.201 | -0.207 | 0.010 | 0.185 | -0.116 | -0.044 | 0.182 | -0.195 | 0.259 | -0.089 | -0.288 | -0.281 | -0.057 | -0.088 |
| | **P** | 0.092 | 0.730 | 0.266 | 0.249 | **0.005** | 0.804 | 0.104 | 0.553 | 0.470 | 0.451 | 0.318 | 0.612 | 0.352 | 0.056 | 0.904 | 0.175 | 0.461 | 0.589 | 0.521 | 0.913 | 0.285 | 0.105 | 0.357 | 0.326 | 0.311 | 0.960 | 0.367 | 0.572 | 0.829 | 0.374 | 0.340 | 0.202 | 0.667 | 0.153 | 0.165 | 0.783 | 0.670 |
| **CD49e** | **R** | 0.586 | 0.110 | -0.198 | -0.275 | -0.544 | 0.210 | -0.119 | -0.074 | -0.050 | -0.136 | -0.194 | -0.159 | -0.109 | 0.460 | -0.146 | 0.517 | -0.223 | -0.054 | 0.182 | -0.016 | -0.216 | 0.232 | -0.091 | -0.160 | -0.268 | -0.099 | 0.011 | -0.170 | -0.040 | -0.033 | -0.002 | 0.397 | -0.011 | -0.240 | -0.281 | -0.020 | 0.054 |
| | **P** | **0.002** | 0.593 | 0.333 | 0.174 | **0.004** | 0.304 | 0.563 | 0.719 | 0.809 | 0.506 | 0.343 | 0.437 | 0.596 | **0.018** | 0.476 | **0.007** | 0.272 | 0.794 | 0.373 | 0.938 | 0.290 | 0.254 | 0.659 | 0.436 | 0.185 | 0.631 | 0.958 | 0.407 | 0.847 | 0.871 | 0.993 | **0.045** | 0.957 | 0.238 | 0.164 | 0.924 | 0.793 |
| **CD56** | **R** | 0.266 | -0.064 | -0.145 | 0.006 | -0.404 | 0.260 | -0.098 | 0.124 | -0.063 | -0.079 | 0.093 | -0.243 | -0.205 | 0.625 | 0.390 | 0.148 | -0.214 | 0.136 | 0.328 | 0.069 | -0.128 | 0.505 | 0.049 | -0.282 | -0.447 | -0.312 | 0.163 | -0.009 | 0.270 | 0.243 | 0.016 | 0.288 | 0.030 | -0.099 | -0.062 | 0.159 | -0.035 |
| | **P** | 0.190 | 0.754 | 0.480 | 0.975 | **0.040** | 0.199 | 0.634 | 0.545 | 0.758 | 0.700 | 0.650 | 0.231 | 0.314 | **0.001** | **0.049** | 0.469 | 0.294 | 0.506 | 0.102 | 0.737 | 0.534 | **0.009** | 0.813 | 0.163 | **0.022** | 0.120 | 0.425 | 0.965 | 0.183 | 0.232 | 0.937 | 0.154 | 0.885 | 0.629 | 0.765 | 0.437 | 0.866 |
| **CD62P** | **R** | -0.131 | -0.020 | 0.052 | -0.234 | -0.076 | 0.136 | -0.282 | 0.010 | -0.047 | -0.022 | -0.087 | -0.200 | -0.176 | -0.155 | -0.100 | 0.157 | -0.216 | -0.176 | 0.043 | -0.141 | 0.313 | 0.309 | 0.165 | -0.126 | -0.257 | -0.290 | -0.202 | -0.137 | -0.111 | -0.220 | -0.085 | 0.043 | -0.112 | -0.306 | -0.285 | 0.037 | -0.321 |
| | **P** | 0.522 | 0.922 | 0.801 | 0.251 | 0.711 | 0.508 | 0.162 | 0.960 | 0.818 | 0.915 | 0.674 | 0.328 | 0.391 | 0.451 | 0.625 | 0.443 | 0.290 | 0.389 | 0.834 | 0.492 | 0.120 | 0.124 | 0.422 | 0.541 | 0.204 | 0.151 | 0.321 | 0.505 | 0.591 | 0.280 | 0.681 | 0.836 | 0.586 | 0.129 | 0.159 | 0.859 | 0.110 |
| **CD63** | **R** | -0.232 | -0.067 | -0.302 | 0.097 | 0.071 | 0.048 | -0.233 | 0.015 | 0.203 | -0.171 | 0.027 | -0.154 | -0.216 | -0.180 | -0.121 | -0.274 | -0.241 | -0.009 | -0.326 | 0.019 | -0.115 | -0.113 | -0.075 | -0.028 | 0.026 | 0.128 | 0.225 | 0.264 | -0.550 | -0.070 | 0.014 | 0.106 | -0.333 | -0.420 | 0.499 | 0.057 | -0.195 |
| | **P** | 0.255 | 0.744 | 0.133 | 0.637 | 0.731 | 0.815 | 0.253 | 0.941 | 0.319 | 0.405 | 0.894 | 0.451 | 0.290 | 0.380 | 0.556 | 0.175 | 0.236 | 0.964 | 0.104 | 0.928 | 0.576 | 0.581 | 0.716 | 0.893 | 0.899 | 0.532 | 0.268 | 0.193 | **0.004** | 0.734 | 0.946 | 0.605 | 0.097 | **0.033** | **0.009** | 0.781 | 0.339 |
| **CD69** | **R** | -0.330 | -0.281 | -0.032 | 0.117 | -0.008 | 0.142 | -0.117 | 0.109 | -0.092 | -0.150 | -0.065 | -0.031 | -0.040 | -0.304 | -0.238 | -0.103 | -0.272 | -0.096 | -0.212 | -0.168 | -0.310 | 0.222 | -0.252 | 0.096 | -0.215 | -0.013 | 0.157 | -0.028 | -0.184 | 0.017 | -0.013 | -0.061 | -0.140 | -0.144 | -0.102 | 0.430 | -0.118 |
| | **P** | 0.100 | 0.165 | 0.876 | 0.569 | 0.971 | 0.490 | 0.570 | 0.597 | 0.656 | 0.464 | 0.753 | 0.882 | 0.845 | 0.131 | 0.241 | 0.616 | 0.179 | 0.640 | 0.299 | 0.411 | 0.123 | 0.276 | 0.215 | 0.639 | 0.291 | 0.948 | 0.445 | 0.893 | 0.368 | 0.934 | 0.951 | 0.767 | 0.494 | 0.483 | 0.621 | **0.028** | 0.567 |
| **CD81** | **R** | -0.323 | 0.083 | 0.109 | 0.003 | 0.247 | -0.215 | 0.237 | 0.109 | -0.122 | 0.002 | 0.129 | 0.122 | 0.018 | -0.178 | 0.064 | -0.184 | 0.325 | -0.076 | 0.171 | -0.071 | 0.388 | -0.077 | 0.214 | 0.087 | 0.190 | 0.033 | -0.224 | -0.167 | 0.338 | 0.030 | -0.028 | -0.506 | 0.099 | 0.434 | -0.159 | -0.398 | -0.078 |
| | **P** | 0.107 | 0.687 | 0.595 | 0.987 | 0.223 | 0.291 | 0.244 | 0.598 | 0.553 | 0.992 | 0.530 | 0.552 | 0.931 | 0.385 | 0.755 | 0.369 | 0.105 | 0.713 | 0.404 | 0.729 | **0.050** | 0.708 | 0.295 | 0.672 | 0.351 | 0.872 | 0.272 | 0.413 | 0.091 | 0.885 | 0.891 | **0.008** | 0.629 | **0.027** | 0.439 | **0.044** | 0.704 |
| **CD86** | **R** | 0.448 | 0.045 | -0.167 | -0.129 | -0.124 | -0.011 | -0.160 | -0.038 | -0.079 | 0.079 | -0.082 | -0.050 | 0.242 | 0.189 | -0.022 | 0.439 | -0.263 | -0.137 | 0.058 | -0.075 | -0.161 | 0.350 | -0.187 | 0.178 | -0.176 | -0.172 | -0.057 | -0.155 | -0.126 | -0.207 | -0.122 | 0.617 | -0.128 | -0.347 | -0.153 | 0.510 | -0.035 |
| | **P** | **0.022** | 0.825 | 0.415 | 0.529 | 0.547 | 0.958 | 0.436 | 0.854 | 0.700 | 0.702 | 0.689 | 0.810 | 0.234 | 0.356 | 0.917 | **0.025** | 0.194 | 0.504 | 0.780 | 0.716 | 0.432 | 0.080 | 0.361 | 0.378 | 0.389 | 0.400 | 0.783 | 0.448 | 0.539 | 0.310 | 0.552 | **0.001** | 0.532 | 0.083 | 0.457 | **0.008** | 0.865 |
| **CD105** | **R** | -0.031 | -0.157 | -0.179 | -0.068 | 0.006 | -0.026 | -0.179 | 0.063 | -0.105 | -0.047 | -0.001 | -0.014 | 0.117 | -0.131 | -0.083 | 0.023 | -0.244 | -0.193 | -0.162 | -0.168 | -0.135 | 0.396 | -0.160 | 0.160 | -0.190 | -0.072 | -0.045 | -0.153 | -0.256 | -0.232 | -0.145 | 0.351 | -0.170 | -0.335 | -0.141 | 0.595 | -0.190 |
| | **P** | 0.881 | 0.444 | 0.381 | 0.741 | 0.977 | 0.900 | 0.382 | 0.761 | 0.609 | 0.819 | 0.994 | 0.944 | 0.569 | 0.525 | 0.687 | 0.912 | 0.229 | 0.344 | 0.428 | 0.413 | 0.510 | **0.045** | 0.434 | 0.435 | 0.352 | 0.728 | 0.829 | 0.454 | 0.207 | 0.255 | 0.481 | 0.079 | 0.407 | 0.094 | 0.491 | **0.001** | 0.353 |
| **CD133/1** | **R** | 0.167 | -0.087 | -0.227 | -0.111 | -0.121 | -0.033 | -0.129 | 0.017 | -0.051 | -0.078 | 0.029 | -0.166 | 0.046 | 0.013 | 0.021 | 0.057 | -0.297 | -0.148 | -0.100 | -0.120 | -0.179 | 0.352 | -0.128 | 0.027 | -0.162 | -0.096 | -0.016 | -0.133 | -0.303 | -0.111 | -0.092 | 0.492 | -0.198 | -0.396 | -0.093 | 0.464 | -0.197 |
| | **P** | 0.415 | 0.673 | 0.265 | 0.588 | 0.555 | 0.875 | 0.530 | 0.933 | 0.803 | 0.706 | 0.887 | 0.417 | 0.823 | 0.948 | 0.917 | 0.781 | 0.141 | 0.472 | 0.627 | 0.559 | 0.381 | 0.078 | 0.532 | 0.895 | 0.428 | 0.642 | 0.937 | 0.517 | 0.132 | 0.591 | 0.659 | **0.011** | 0.332 | **0.045** | 0.652 | **0.017** | 0.335 |
| **CD142** | **R** | 0.091 | -0.071 | -0.126 | -0.170 | -0.038 | -0.126 | -0.263 | -0.026 | -0.170 | -0.020 | -0.093 | -0.038 | 0.055 | -0.107 | -0.199 | 0.119 | -0.138 | -0.257 | -0.086 | -0.169 | -0.183 | 0.344 | -0.234 | -0.010 | -0.093 | 0.030 | -0.014 | -0.123 | -0.232 | -0.170 | -0.183 | 0.390 | -0.181 | -0.348 | -0.248 | 0.391 | -0.162 |
| | **P** | 0.657 | 0.729 | 0.539 | 0.407 | 0.852 | 0.539 | 0.193 | 0.901 | 0.408 | 0.924 | 0.650 | 0.853 | 0.789 | 0.602 | 0.329 | 0.564 | 0.501 | 0.205 | 0.677 | 0.409 | 0.371 | 0.085 | 0.250 | 0.962 | 0.651 | 0.884 | 0.945 | 0.550 | 0.255 | 0.407 | 0.370 | **0.049** | 0.375 | 0.081 | 0.223 | **0.048** | 0.430 |
| **CD146** | **R** | 0.297 | -0.031 | -0.136 | -0.125 | -0.125 | -0.084 | -0.283 | -0.041 | -0.090 | 0.056 | -0.022 | -0.146 | 0.055 | 0.128 | 0.055 | 0.210 | -0.222 | -0.156 | 0.002 | -0.081 | -0.087 | 0.434 | -0.113 | -0.051 | -0.170 | -0.143 | -0.110 | -0.217 | -0.148 | -0.148 | -0.200 | 0.558 | -0.131 | -0.412 | -0.195 | 0.420 | -0.148 |
| | **P** | 0.141 | 0.882 | 0.507 | 0.543 | 0.544 | 0.683 | 0.161 | 0.842 | 0.663 | 0.787 | 0.914 | 0.478 | 0.790 | 0.533 | 0.789 | 0.303 | 0.275 | 0.447 | 0.994 | 0.696 | 0.674 | **0.027** | 0.583 | 0.804 | 0.407 | 0.486 | 0.593 | 0.287 | 0.472 | 0.469 | 0.327 | **0.003** | 0.524 | **0.037** | 0.339 | **0.033** | 0.471 |
| **CD209** | **R** | 0.414 | 0.137 | -0.095 | -0.129 | -0.133 | -0.027 | -0.155 | 0.051 | -0.112 | 0.189 | 0.004 | 0.049 | 0.247 | 0.262 | 0.043 | 0.498 | -0.171 | -0.091 | 0.291 | 0.005 | -0.141 | 0.372 | -0.131 | 0.221 | -0.177 | -0.200 | 0.055 | -0.154 | 0.073 | -0.026 | -0.077 | 0.605 | -0.119 | -0.244 | -0.191 | 0.532 | -0.024 |
| | **P** | **0.035** | 0.506 | 0.645 | 0.530 | 0.517 | 0.895 | 0.449 | 0.806 | 0.585 | 0.355 | 0.983 | 0.811 | 0.223 | 0.196 | 0.836 | **0.010** | 0.403 | 0.658 | 0.150 | 0.981 | 0.492 | 0.061 | 0.524 | 0.279 | 0.388 | 0.326 | 0.788 | 0.453 | 0.724 | 0.901 | 0.707 | **0.001** | 0.563 | 0.229 | 0.351 | **0.005** | 0.906 |
| **CD326** | **R** | -0.278 | -0.194 | -0.222 | -0.002 | 0.037 | -0.032 | -0.119 | 0.033 | 0.020 | -0.279 | 0.194 | -0.032 | 0.065 | -0.280 | 0.014 | -0.272 | -0.259 | -0.054 | -0.287 | -0.056 | -0.227 | 0.018 | -0.159 | 0.226 | 0.013 | 0.152 | 0.287 | 0.031 | -0.327 | -0.075 | -0.123 | -0.001 | -0.226 | -0.305 | 0.275 | 0.268 | -0.248 |
| | **P** | 0.169 | 0.343 | 0.276 | 0.994 | 0.859 | 0.875 | 0.564 | 0.874 | 0.923 | 0.168 | 0.342 | 0.876 | 0.752 | 0.167 | 0.945 | 0.178 | 0.201 | 0.792 | 0.155 | 0.784 | 0.265 | 0.932 | 0.437 | 0.268 | 0.949 | 0.459 | 0.155 | 0.880 | 0.102 | 0.714 | 0.549 | 0.996 | 0.268 | 0.129 | 0.174 | 0.186 | 0.221 |
| **HLA-I** | **R** | 0.827 | 0.129 | 0.015 | -0.215 | -0.407 | 0.117 | -0.074 | -0.238 | -0.112 | 0.088 | -0.367 | -0.169 | 0.018 | 0.433 | -0.073 | 0.640 | -0.197 | -0.128 | 0.145 | -0.085 | -0.129 | 0.182 | -0.176 | -0.155 | -0.101 | -0.187 | -0.206 | -0.122 | -0.042 | -0.190 | -0.023 | 0.546 | 0.056 | -0.209 | -0.247 | 0.080 | 0.158 |
| | **P** | **0.000** | 0.530 | 0.943 | 0.291 | **0.039** | 0.568 | 0.718 | 0.243 | 0.586 | 0.668 | 0.065 | 0.408 | 0.931 | **0.027** | 0.722 | **0.000** | 0.335 | 0.532 | 0.480 | 0.680 | 0.530 | 0.374 | 0.388 | 0.450 | 0.386 | 0.360 | 0.313 | 0.553 | 0.838 | 0.352 | 0.911 | **0.004** | 0.784 | 0.306 | 0.224 | 0.698 | 0.442 |
| **HLA-II** | **R** | 0.558 | 0.063 | -0.200 | -0.004 | -0.509 | 0.181 | -0.271 | -0.218 | -0.112 | -0.053 | -0.464 | -0.092 | -0.203 | 0.587 | -0.003 | 0.414 | -0.210 | -0.141 | 0.170 | -0.047 | -0.183 | 0.296 | -0.237 | -0.125 | -0.282 | -0.300 | -0.105 | -0.148 | -0.145 | -0.036 | -0.130 | 0.383 | -0.106 | -0.247 | -0.139 | 0.005 | -0.021 |
| | **P** | **0.003** | 0.761 | 0.328 | 0.985 | **0.008** | 0.375 | 0.180 | 0.284 | 0.585 | 0.797 | **0.017** | 0.655 | 0.321 | **0.002** | 0.988 | **0.035** | 0.302 | 0.493 | 0.406 | 0.820 | 0.370 | 0.142 | 0.244 | 0.542 | 0.163 | 0.136 | 0.609 | 0.471 | 0.479 | 0.863 | 0.527 | 0.054 | 0.607 | 0.223 | 0.498 | 0.981 | 0.919 |
| **MCSP** | **R** | 0.395 | 0.000 | -0.170 | -0.184 | -0.151 | 0.014 | -0.237 | -0.068 | -0.109 | 0.017 | -0.082 | -0.112 | 0.136 | 0.130 | -0.046 | 0.342 | -0.277 | -0.191 | -0.003 | -0.109 | -0.140 | 0.394 | -0.179 | 0.062 | -0.217 | -0.151 | -0.065 | -0.173 | -0.187 | -0.239 | -0.205 | 0.569 | -0.142 | -0.391 | -0.178 | 0.460 | -0.106 |
| | **P** | **0.046** | 0.998 | 0.407 | 0.367 | 0.461 | 0.946 | 0.244 | 0.742 | 0.596 | 0.935 | 0.692 | 0.586 | 0.507 | 0.527 | 0.823 | 0.087 | 0.171 | 0.349 | 0.987 | 0.597 | 0.496 | **0.047** | 0.381 | 0.764 | 0.287 | 0.461 | 0.752 | 0.399 | 0.360 | 0.239 | 0.315 | **0.002** | 0.490 | **0.048** | 0.384 | **0.018** | 0.605 |
| **ROR1** | **R** | 0.401 | 0.023 | -0.274 | -0.187 | -0.248 | -0.005 | -0.167 | -0.118 | -0.013 | -0.156 | -0.093 | -0.163 | 0.026 | 0.184 | -0.105 | 0.231 | -0.270 | -0.133 | -0.082 | -0.061 | -0.305 | 0.172 | -0.213 | -0.031 | -0.087 | 0.024 | 0.062 | -0.030 | -0.336 | -0.151 | -0.075 | 0.525 | -0.236 | -0.438 | -0.019 | 0.198 | -0.099 |
| | **P** | **0.042** | 0.911 | 0.175 | 0.360 | 0.221 | 0.980 | 0.416 | 0.564 | 0.951 | 0.446 | 0.653 | 0.425 | 0.898 | 0.369 | 0.611 | 0.256 | 0.182 | 0.516 | 0.692 | 0.766 | 0.130 | 0.401 | 0.297 | 0.880 | 0.673 | 0.908 | 0.764 | 0.885 | 0.093 | 0.460 | 0.717 | **0.006** | 0.245 | **0.025** | 0.928 | 0.333 | 0.631 |
| **SSEA4** | **R** | 0.287 | -0.141 | -0.282 | -0.194 | -0.536 | 0.069 | -0.124 | -0.171 | -0.165 | -0.380 | -0.123 | -0.191 | -0.136 | 0.392 | 0.215 | 0.128 | -0.321 | -0.132 | 0.030 | -0.097 | -0.318 | 0.262 | -0.252 | -0.051 | -0.186 | -0.068 | 0.097 | -0.068 | -0.190 | 0.099 | -0.103 | 0.187 | -0.200 | -0.328 | -0.028 | -0.030 | -0.246 |
| | **P** | 0.155 | 0.494 | 0.163 | 0.341 | **0.005** | 0.739 | 0.545 | 0.403 | 0.421 | 0.055 | 0.550 | 0.350 | 0.507 | **0.048** | 0.291 | 0.535 | 0.110 | 0.520 | 0.886 | 0.638 | 0.114 | 0.196 | 0.215 | 0.803 | 0.364 | 0.740 | 0.637 | 0.741 | 0.353 | 0.631 | 0.616 | 0.360 | 0.328 | 0.102 | 0.891 | 0.884 | 0.225 |

| | | CD1c | CD2 | CD3 | CD4 | CD8 | CD9 | CD11c | CD14 | CD19 | CD20 | CD24 | CD25 | CD29 | CD31 | CD40 | CD41b | CD42a | CD44 | CD45 | CD49e | CD56 | CD62P | CD63 | CD69 | CD81 | CD86 | CD105 | CD133/1 | CD142 | CD146 | CD209 | CD326 | HLA-I | HLA-II | MCSP | ROR1 | SSEA4 |
|---|---|---|---|---|---|---|---|---|---|---|---|---|---|---|---|---|---|---|---|---|---|---|---|---|---|---|---|---|---|---|---|---|---|---|---|---|---|---|
| CD1c | R | -0.175 | -0.053 | 0.193 | -0.086 | -0.090 | -0.160 | -0.133 | 0.004 | -0.074 | -0.067 | -0.054 | -0.016 | -0.057 | 0.001 | -0.145 | -0.038 | -0.262 | 0.090 | -0.020 | -0.029 | -0.108 | 0.127 | -0.101 | -0.009 | 0.016 | -0.169 | 0.032 | -0.337 | -0.027 | 0.046 | 0.311 | -0.104 | -0.148 | -0.035 | -0.154 | -0.069 | -0.052 |
| | P | 0.219 | 0.710 | 0.174 | 0.550 | 0.528 | 0.261 | 0.353 | 0.977 | 0.606 | 0.639 | 0.705 | 0.909 | 0.691 | 0.997 | 0.312 | 0.792 | 0.063 | 0.530 | 0.890 | 0.840 | 0.450 | 0.375 | 0.480 | 0.949 | 0.911 | 0.235 | 0.825 | **0.016** | 0.849 | 0.749 | **0.027** | 0.466 | 0.300 | 0.808 | 0.282 | 0.630 | 0.715 |
| CD2 | R | -0.017 | 0.175 | -0.054 | 0.224 | -0.017 | -0.099 | 0.093 | 0.137 | 0.027 | -0.065 | 0.001 | -0.028 | 0.007 | 0.080 | 0.037 | 0.024 | -0.121 | 0.076 | -0.032 | 0.038 | -0.052 | 0.287 | -0.077 | -0.049 | -0.156 | -0.078 | 0.059 | -0.337 | -0.145 | 0.228 | 0.006 | 0.047 | -0.010 | 0.080 | -0.093 | 0.014 | 0.101 |
| | P | 0.906 | 0.220 | 0.709 | 0.114 | 0.908 | 0.488 | 0.517 | 0.339 | 0.853 | 0.650 | 0.993 | 0.847 | 0.960 | 0.578 | 0.797 | 0.866 | 0.396 | 0.595 | 0.825 | 0.789 | 0.719 | **0.041** | 0.593 | 0.734 | 0.275 | 0.586 | 0.683 | **0.015** | 0.311 | 0.107 | 0.966 | 0.744 | 0.947 | 0.575 | 0.518 | 0.922 | 0.483 |
| CD3 | R | -0.180 | -0.022 | 0.071 | 0.056 | -0.085 | -0.046 | -0.055 | 0.041 | -0.071 | -0.167 | -0.145 | -0.064 | -0.067 | 0.037 | -0.016 | -0.051 | -0.318 | 0.080 | -0.079 | -0.030 | -0.081 | 0.241 | -0.072 | -0.073 | -0.158 | -0.273 | 0.044 | -0.398 | -0.203 | 0.017 | 0.256 | -0.074 | -0.095 | -0.041 | -0.087 | -0.073 | 0.010 |
| | P | 0.205 | 0.875 | 0.618 | 0.696 | 0.553 | 0.749 | 0.699 | 0.777 | 0.619 | 0.240 | 0.311 | 0.654 | 0.641 | 0.795 | 0.912 | 0.720 | **0.023** | 0.578 | 0.583 | 0.834 | 0.571 | 0.088 | 0.614 | 0.611 | 0.267 | 0.053 | 0.760 | **0.004** | 0.153 | 0.904 | 0.070 | 0.606 | 0.508 | 0.776 | 0.544 | 0.612 | 0.945 |
| CD4 | R | -0.114 | -0.017 | 0.246 | -0.077 | -0.012 | -0.170 | -0.093 | 0.021 | -0.034 | 0.024 | -0.040 | 0.080 | 0.035 | -0.047 | -0.116 | -0.039 | -0.204 | 0.113 | -0.085 | 0.038 | -0.098 | 0.174 | -0.074 | 0.078 | -0.014 | -0.104 | 0.071 | -0.258 | 0.049 | -0.001 | 0.210 | -0.031 | -0.104 | 0.006 | -0.068 | 0.011 | 0.024 |
| | P | 0.427 | 0.905 | 0.082 | 0.591 | 0.934 | 0.232 | 0.517 | 0.883 | 0.813 | 0.865 | 0.781 | 0.577 | 0.806 | 0.744 | 0.418 | 0.784 | 0.151 | 0.428 | 0.551 | 0.791 | 0.493 | 0.222 | 0.607 | 0.586 | 0.921 | 0.470 | 0.620 | 0.067 | 0.734 | 0.993 | 0.138 | 0.830 | 0.468 | 0.966 | 0.633 | 0.941 | 0.869 |
| CD8 | R | -0.137 | 0.033 | 0.109 | -0.072 | -0.087 | -0.074 | -0.065 | -0.014 | -0.049 | -0.045 | -0.082 | 0.000 | 0.011 | -0.010 | 0.001 | -0.027 | -0.128 | -0.055 | -0.153 | 0.041 | -0.065 | 0.149 | -0.043 | 0.019 | -0.065 | -0.141 | 0.043 | -0.207 | 0.057 | -0.016 | 0.104 | -0.014 | -0.152 | -0.005 | -0.034 | -0.002 | 0.009 |
| | P | 0.339 | 0.818 | 0.447 | 0.616 | 0.543 | 0.608 | 0.650 | 0.920 | 0.733 | 0.753 | 0.566 | 0.998 | 0.941 | 0.944 | 0.994 | 0.853 | 0.369 | 0.699 | 0.283 | 0.773 | 0.650 | 0.296 | 0.762 | 0.894 | 0.650 | 0.324 | 0.764 | 0.145 | 0.691 | 0.913 | 0.470 | 0.922 | 0.288 | 0.970 | 0.815 | 0.989 | 0.950 |
| CD9 | R | 0.211 | -0.001 | 0.222 | 0.256 | 0.038 | 0.067 | 0.303 | 0.259 | 0.338 | 0.237 | 0.238 | 0.269 | 0.311 | -0.062 | -0.199 | 0.215 | 0.185 | -0.092 | -0.192 | 0.353 | 0.319 | 0.270 | -0.145 | 0.298 | -0.264 | 0.099 | 0.239 | -0.013 | -0.106 | -0.266 | -0.038 | 0.308 | 0.237 | 0.017 | 0.122 | 0.302 | 0.263 |
| | P | 0.138 | 0.996 | 0.117 | 0.070 | 0.790 | 0.638 | **0.031** | 0.067 | **0.015** | 0.095 | 0.093 | 0.056 | **0.026** | 0.667 | 0.161 | 0.130 | 0.195 | 0.522 | 0.176 | **0.011** | **0.023** | 0.056 | 0.309 | **0.034** | 0.062 | 0.492 | 0.092 | 0.927 | 0.459 | 0.059 | 0.792 | **0.028** | 0.094 | 0.907 | 0.392 | **0.031** | 0.063 |
| CD11c | R | -0.147 | 0.048 | -0.105 | 0.083 | -0.068 | -0.108 | 0.028 | 0.014 | -0.114 | 0.009 | -0.054 | -0.086 | -0.052 | -0.010 | -0.058 | -0.057 | -0.010 | -0.046 | -0.159 | 0.018 | -0.098 | 0.081 | -0.133 | -0.086 | 0.014 | 0.010 | -0.002 | -0.403 | 0.098 | 0.250 | -0.033 | -0.032 | -0.071 | 0.125 | -0.047 | -0.069 | -0.013 |
| | P | 0.303 | 0.737 | 0.462 | 0.565 | 0.634 | 0.451 | 0.847 | 0.922 | 0.424 | 0.952 | 0.707 | 0.548 | 0.719 | 0.942 | 0.688 | 0.694 | 0.945 | 0.749 | 0.265 | 0.900 | 0.492 | 0.574 | 0.352 | 0.547 | 0.920 | 0.946 | 0.989 | **0.003** | 0.496 | 0.077 | 0.817 | 0.824 | 0.622 | 0.384 | 0.744 | 0.632 | 0.926 |
| CD14 | R | -0.123 | 0.008 | 0.262 | -0.119 | -0.079 | -0.133 | -0.096 | 0.079 | 0.011 | -0.025 | -0.017 | 0.057 | 0.013 | -0.052 | -0.127 | -0.031 | -0.219 | 0.041 | -0.016 | 0.008 | -0.091 | 0.141 | -0.048 | 0.069 | -0.016 | -0.147 | 0.013 | -0.250 | -0.016 | -0.025 | 0.177 | -0.046 | -0.097 | -0.006 | -0.120 | 0.003 | -0.011 |
| | P | 0.391 | 0.958 | 0.063 | 0.407 | 0.584 | 0.354 | 0.504 | 0.580 | 0.939 | 0.864 | 0.907 | 0.693 | 0.927 | 0.719 | 0.375 | 0.831 | 0.122 | 0.774 | 0.912 | 0.958 | 0.527 | 0.323 | 0.738 | 0.629 | 0.914 | 0.304 | 0.930 | 0.077 | 0.911 | 0.860 | 0.214 | 0.749 | 0.497 | 0.965 | 0.403 | 0.984 | 0.938 |
| CD19 | R | 0.022 | 0.009 | 0.337 | 0.121 | 0.055 | -0.211 | 0.093 | 0.192 | 0.066 | -0.007 | 0.049 | 0.147 | 0.153 | 0.056 | -0.144 | 0.112 | -0.061 | 0.176 | -0.123 | 0.127 | 0.007 | 0.362 | -0.084 | 0.082 | -0.134 | -0.089 | 0.189 | -0.293 | -0.062 | 0.011 | 0.136 | 0.132 | 0.059 | 0.063 | -0.006 | 0.137 | 0.055 |
| | P | 0.877 | 0.952 | **0.016** | 0.399 | 0.700 | 0.136 | 0.518 | 0.176 | 0.646 | 0.962 | 0.732 | 0.303 | 0.283 | 0.695 | 0.312 | 0.434 | 0.673 | 0.216 | 0.391 | 0.374 | 0.962 | **0.009** | 0.557 | 0.566 | 0.348 | 0.534 | 0.185 | **0.037** | 0.666 | 0.939 | 0.342 | 0.357 | 0.683 | 0.662 | 0.964 | 0.336 | 0.703 |
| CD20 | R | 0.069 | -0.028 | 0.419 | 0.027 | 0.187 | -0.269 | 0.189 | 0.205 | 0.105 | -0.019 | 0.045 | 0.191 | 0.218 | 0.042 | -0.167 | 0.154 | 0.001 | 0.076 | -0.157 | 0.167 | 0.016 | 0.372 | -0.028 | 0.107 | -0.100 | -0.001 | 0.268 | -0.178 | -0.008 | -0.029 | 0.054 | 0.192 | 0.153 | 0.196 | 0.009 | 0.182 | 0.085 |
| | P | 0.631 | 0.846 | **0.002** | 0.853 | 0.190 | 0.056 | 0.184 | 0.150 | 0.465 | 0.897 | 0.755 | 0.179 | 0.125 | 0.771 | 0.242 | 0.281 | 0.993 | 0.596 | 0.273 | 0.241 | 0.913 | **0.007** | 0.846 | 0.456 | 0.486 | 0.994 | 0.058 | 0.211 | 0.954 | 0.839 | 0.707 | 0.176 | 0.282 | 0.169 | 0.948 | 0.200 | 0.551 |
| CD24 | R | 0.060 | -0.023 | 0.288 | 0.074 | 0.057 | -0.093 | 0.225 | 0.251 | 0.118 | -0.049 | 0.052 | 0.144 | 0.207 | -0.101 | -0.148 | 0.115 | -0.022 | 0.028 | -0.194 | 0.144 | 0.038 | 0.405 | -0.116 | 0.097 | -0.254 | -0.113 | 0.163 | -0.255 | -0.118 | -0.066 | 0.055 | 0.187 | 0.140 | 0.033 | 0.027 | 0.177 | 0.051 |
| | P | 0.673 | 0.875 | **0.041** | 0.605 | 0.693 | 0.517 | 0.112 | 0.076 | 0.411 | 0.733 | 0.719 | 0.313 | 0.145 | 0.481 | 0.302 | 0.423 | 0.878 | 0.847 | 0.171 | 0.313 | 0.789 | **0.003** | 0.417 | 0.500 | 0.072 | 0.428 | 0.253 | 0.070 | 0.408 | 0.646 | 0.701 | 0.189 | 0.326 | 0.820 | 0.849 | 0.215 | 0.723 |
| CD25 | R | 0.016 | -0.017 | 0.339 | -0.058 | 0.108 | -0.294 | 0.165 | 0.167 | 0.043 | 0.001 | 0.048 | 0.144 | 0.160 | -0.082 | -0.191 | 0.082 | -0.054 | 0.000 | -0.186 | 0.154 | -0.008 | 0.301 | -0.030 | 0.089 | -0.025 | -0.034 | 0.207 | -0.283 | -0.018 | 0.010 | 0.193 | 0.119 | 0.107 | 0.163 | -0.040 | 0.121 | 0.086 |
| | P | 0.913 | 0.905 | **0.015** | 0.687 | 0.449 | **0.036** | 0.248 | 0.240 | 0.765 | 0.993 | 0.739 | 0.314 | 0.262 | 0.567 | 0.179 | 0.565 | 0.708 | 0.998 | 0.191 | 0.280 | 0.956 | **0.032** | 0.837 | 0.533 | 0.860 | 0.815 | 0.145 | **0.044** | 0.900 | 0.943 | 0.174 | 0.407 | 0.456 | 0.254 | 0.779 | 0.396 | 0.547 |
| CD29 | R | -0.084 | -0.002 | 0.183 | 0.009 | -0.054 | 0.186 | 0.050 | -0.001 | 0.126 | -0.112 | 0.114 | -0.055 | -0.041 | 0.103 | -0.145 | 0.108 | -0.085 | -0.131 | 0.099 | 0.016 | 0.097 | 0.067 | -0.052 | -0.061 | -0.188 | -0.078 | 0.000 | -0.194 | -0.278 | -0.170 | 0.172 | -0.004 | 0.122 | -0.079 | -0.067 | 0.009 | -0.050 |
| | P | 0.557 | 0.989 | 0.200 | 0.949 | 0.706 | 0.192 | 0.730 | 0.996 | 0.379 | 0.433 | 0.427 | 0.703 | 0.776 | 0.472 | 0.311 | 0.452 | 0.554 | 0.360 | 0.490 | 0.913 | 0.498 | 0.642 | 0.718 | 0.671 | 0.185 | 0.586 | 0.998 | 0.172 | **0.048** | 0.232 | 0.228 | 0.979 | 0.392 | 0.582 | 0.641 | 0.950 | 0.728 |
| CD31 | R | 0.217 | 0.010 | 0.289 | 0.097 | 0.271 | -0.280 | 0.325 | 0.124 | 0.161 | 0.173 | 0.144 | 0.251 | 0.319 | -0.146 | -0.116 | 0.173 | 0.162 | -0.113 | -0.250 | 0.345 | 0.047 | 0.242 | 0.051 | 0.172 | 0.011 | 0.184 | 0.313 | 0.004 | 0.110 | 0.028 | 0.012 | 0.274 | 0.264 | 0.293 | 0.100 | 0.244 | 0.187 |
| | P | 0.126 | 0.944 | **0.040** | 0.501 | 0.054 | **0.047** | **0.020** | 0.388 | 0.258 | 0.225 | 0.312 | 0.076 | **0.023** | 0.308 | 0.418 | 0.224 | 0.257 | 0.430 | 0.077 | **0.013** | 0.744 | 0.087 | 0.720 | 0.227 | 0.937 | 0.195 | **0.025** | 0.978 | 0.443 | 0.847 | 0.933 | **0.052** | 0.062 | **0.037** | 0.486 | 0.085 | 0.189 |
| CD40 | R | 0.119 | -0.006 | 0.216 | 0.044 | 0.225 | -0.211 | 0.288 | 0.079 | 0.087 | 0.077 | 0.077 | 0.166 | 0.220 | -0.176 | -0.078 | 0.119 | 0.009 | -0.102 | -0.278 | 0.272 | 0.020 | 0.248 | -0.070 | 0.121 | -0.043 | 0.077 | 0.258 | -0.110 | 0.050 | 0.061 | 0.133 | 0.188 | 0.180 | 0.209 | 0.099 | 0.158 | 0.188 |
| | P | 0.405 | 0.967 | 0.128 | 0.761 | 0.112 | 0.137 | **0.040** | 0.579 | 0.543 | 0.595 | 0.589 | 0.244 | 0.121 | 0.216 | 0.587 | 0.405 | 0.953 | 0.478 | **0.048** | 0.053 | 0.891 | 0.079 | 0.626 | 0.396 | 0.763 | 0.593 | 0.068 | 0.444 | 0.726 | 0.670 | 0.351 | 0.187 | 0.205 | 0.142 | 0.489 | 0.269 | 0.186 |
| CD41b | R | 0.158 | 0.152 | 0.074 | 0.068 | 0.195 | -0.083 | 0.267 | 0.087 | 0.153 | 0.206 | 0.144 | 0.165 | 0.210 | -0.163 | -0.142 | 0.119 | 0.086 | -0.191 | -0.123 | 0.306 | 0.037 | 0.133 | -0.012 | 0.143 | -0.046 | 0.171 | 0.170 | -0.058 | -0.013 | 0.152 | 0.016 | 0.217 | 0.218 | 0.226 | 0.125 | 0.186 | 0.148 |
| | P | 0.268 | 0.286 | 0.607 | 0.635 | 0.170 | 0.563 | 0.058 | 0.546 | 0.284 | 0.148 | 0.315 | 0.247 | 0.139 | 0.253 | 0.321 | 0.405 | 0.550 | 0.179 | 0.391 | **0.029** | 0.797 | 0.350 | 0.931 | 0.318 | 0.750 | 0.230 | 0.233 | 0.685 | 0.929 | 0.286 | 0.913 | 0.126 | 0.125 | 0.110 | 0.381 | 0.192 | 0.301 |
| CD42a | R | 0.025 | 0.056 | -0.100 | 0.066 | 0.084 | -0.012 | 0.153 | -0.133 | 0.026 | 0.099 | 0.048 | 0.044 | 0.050 | -0.219 | 0.007 | -0.027 | -0.015 | -0.161 | -0.268 | 0.151 | 0.046 | -0.047 | -0.049 | 0.013 | 0.046 | 0.233 | 0.103 | -0.005 | 0.053 | 0.078 | 0.032 | 0.043 | 0.148 | 0.155 | 0.012 | 0.006 | 0.145 |
| | P | 0.864 | 0.697 | 0.485 | 0.647 | 0.560 | 0.934 | 0.284 | 0.354 | 0.854 | 0.488 | 0.740 | 0.761 | 0.728 | 0.123 | 0.961 | 0.852 | 0.917 | 0.258 | 0.057 | 0.290 | 0.748 | 0.744 | 0.731 | 0.928 | 0.749 | 0.100 | 0.472 | 0.974 | 0.713 | 0.588 | 0.822 | 0.763 | 0.300 | 0.279 | 0.935 | 0.966 | 0.311 |
| CD44 | R | 0.077 | -0.011 | 0.394 | 0.057 | 0.071 | -0.178 | 0.185 | 0.155 | 0.109 | 0.100 | 0.119 | 0.208 | 0.228 | 0.030 | -0.121 | 0.133 | 0.046 | 0.059 | -0.125 | 0.180 | 0.049 | 0.366 | -0.076 | 0.132 | -0.162 | 0.005 | 0.265 | -0.257 | -0.051 | -0.100 | 0.033 | 0.206 | 0.098 | 0.105 | 0.012 | 0.199 | 0.119 |
| | P | 0.593 | 0.939 | **0.004** | 0.689 | 0.623 | 0.213 | 0.194 | 0.277 | 0.445 | 0.483 | 0.406 | 0.144 | 0.107 | 0.835 | 0.399 | 0.352 | 0.747 | 0.681 | 0.380 | 0.207 | 0.731 | **0.008** | 0.596 | 0.357 | 0.256 | 0.970 | 0.060 | 0.069 | 0.724 | 0.484 | 0.819 | 0.148 | 0.492 | 0.464 | 0.934 | 0.162 | 0.405 |
| CD45 | R | -0.048 | 0.016 | 0.324 | -0.101 | 0.046 | -0.130 | 0.093 | 0.152 | 0.086 | 0.035 | -0.022 | 0.127 | 0.129 | 0.008 | -0.054 | 0.087 | -0.129 | 0.054 | -0.087 | 0.133 | -0.073 | 0.287 | -0.046 | 0.080 | -0.134 | -0.081 | 0.149 | -0.306 | -0.041 | 0.066 | 0.100 | 0.103 | 0.080 | 0.128 | -0.024 | 0.112 | 0.071 |
| | P | 0.739 | 0.912 | **0.020** | 0.480 | 0.750 | 0.365 | 0.517 | 0.288 | 0.548 | 0.806 | 0.876 | 0.373 | 0.368 | 0.956 | 0.706 | 0.546 | 0.366 | 0.705 | 0.544 | 0.351 | 0.612 | **0.041** | 0.749 | 0.575 | 0.349 | 0.571 | 0.296 | **0.029** | 0.774 | 0.645 | 0.487 | 0.471 | 0.578 | 0.369 | 0.867 | 0.434 | 0.623 |
| CD49e | R | 0.067 | 0.059 | 0.121 | 0.053 | 0.095 | 0.024 | 0.261 | 0.139 | 0.120 | 0.020 | 0.062 | 0.066 | 0.135 | 0.040 | 0.006 | 0.103 | -0.017 | -0.104 | -0.097 | 0.164 | 0.042 | 0.257 | -0.040 | 0.002 | -0.222 | -0.007 | 0.099 | -0.204 | -0.132 | 0.168 | -0.071 | 0.163 | 0.218 | 0.197 | 0.004 | 0.122 | 0.071 |
| | P | 0.640 | 0.679 | 0.398 | 0.713 | 0.508 | 0.868 | 0.064 | 0.329 | 0.402 | 0.888 | 0.668 | 0.644 | 0.345 | 0.778 | 0.965 | 0.472 | 0.907 | 0.467 | 0.496 | 0.250 | 0.771 | 0.069 | 0.778 | 0.987 | 0.118 | 0.960 | 0.490 | 0.150 | 0.356 | 0.240 | 0.619 | 0.254 | 0.125 | 0.167 | 0.978 | 0.394 | 0.622 |
| CD56 | R | -0.050 | 0.098 | -0.040 | 0.045 | -0.209 | 0.235 | 0.068 | 0.104 | 0.047 | -0.188 | -0.083 | -0.085 | -0.022 | 0.065 | 0.125 | 0.031 | -0.191 | -0.116 | -0.050 | -0.019 | 0.061 | 0.269 | -0.091 | -0.072 | -0.385 | -0.320 | -0.094 | -0.315 | -0.366 | 0.037 | -0.012 | 0.042 | -0.044 | -0.124 | 0.009 | 0.014 | -0.012 |
| | P | 0.730 | 0.494 | 0.779 | 0.752 | 0.141 | 0.097 | 0.637 | 0.466 | 0.742 | 0.185 | 0.562 | 0.553 | 0.878 | 0.649 | 0.380 | 0.831 | 0.178 | 0.418 | 0.730 | 0.894 | 0.669 | 0.056 | 0.524 | 0.616 | **0.005** | **0.022** | 0.513 | **0.024** | **0.008** | 0.794 | 0.931 | 0.769 | 0.759 | 0.387 | 0.952 | 0.921 | 0.936 |
| CD62P | R | -0.117 | -0.121 | 0.083 | 0.099 | -0.075 | 0.550 | -0.109 | 0.073 | 0.046 | -0.120 | 0.190 | -0.108 | -0.094 | 0.293 | -0.131 | -0.005 | -0.080 | 0.045 | 0.237 | -0.065 | -0.083 | 0.046 | -0.158 | -0.126 | -0.437 | -0.225 | -0.120 | -0.160 | -0.283 | -0.071 | -0.007 | -0.023 | -0.076 | -0.170 | -0.101 | -0.051 | -0.130 |
| | P | 0.415 | 0.397 | 0.564 | 0.488 | 0.602 | **0.000** | 0.446 | 0.610 | 0.749 | 0.403 | 0.183 | 0.452 | 0.511 | **0.037** | 0.359 | 0.971 | 0.579 | 0.755 | 0.094 | 0.651 | 0.562 | 0.747 | 0.268 | 0.380 | **0.001** | 0.112 | 0.403 | 0.262 | **0.044** | 0.619 | 0.962 | 0.872 | 0.598 | 0.234 | 0.481 | 0.723 | 0.362 |
| CD63 | R | -0.059 | -0.053 | 0.030 | -0.286 | -0.028 | -0.190 | -0.032 | -0.088 | -0.141 | -0.166 | -0.081 | -0.080 | -0.112 | 0.003 | -0.076 | 0.002 | -0.159 | 0.027 | 0.066 | -0.181 | -0.133 | -0.079 | 0.137 | -0.151 | 0.201 | -0.036 | -0.081 | -0.094 | -0.126 | 0.136 | 0.145 | -0.113 | 0.038 | 0.123 | -0.107 | -0.108 | -0.210 |
| | P | 0.679 | 0.714 | 0.836 | **0.042** | 0.847 | 0.181 | 0.822 | 0.537 | 0.323 | 0.246 | 0.571 | 0.576 | 0.433 | 0.981 | 0.598 | 0.986 | 0.264 | 0.849 | 0.645 | 0.204 | 0.352 | 0.582 | 0.338 | 0.290 | 0.157 | 0.802 | 0.574 | 0.513 | 0.379 | 0.343 | 0.311 | 0.431 | 0.793 | 0.392 | 0.454 | 0.451 | 0.140 |
| CD69 | R | 0.219 | -0.084 | 0.434 | -0.025 | 0.338 | -0.310 | 0.424 | 0.314 | 0.202 | 0.148 | 0.295 | 0.301 | 0.335 | -0.117 | -0.355 | 0.201 | 0.237 | -0.061 | -0.208 | 0.344 | 0.152 | 0.261 | 0.028 | 0.207 | 0.019 | 0.201 | 0.356 | 0.010 | 0.027 | -0.069 | 0.045 | 0.290 | 0.439 | 0.274 | 0.022 | 0.281 | 0.132 |
| | P | 0.123 | 0.558 | **0.001** | 0.860 | **0.015** | **0.027** | **0.002** | **0.025** | 0.154 | 0.301 | **0.035** | **0.032** | **0.016** | 0.412 | **0.010** | 0.156 | 0.094 | 0.671 | 0.144 | **0.013** | 0.285 | 0.064 | 0.846 | 0.145 | 0.897 | 0.158 | **0.010** | 0.945 | 0.852 | 0.631 | 0.753 | **0.039** | **0.001** | **0.052** | 0.879 | **0.046** | 0.356 |
| CD81 | R | -0.143 | 0.061 | -0.257 | 0.069 | -0.007 | 0.149 | -0.267 | -0.159 | -0.179 | -0.049 | -0.146 | -0.179 | -0.184 | 0.058 | 0.286 | -0.219 | -0.004 | 0.061 | 0.118 | -0.148 | -0.168 | -0.181 | -0.010 | -0.127 | 0.035 | -0.058 | -0.147 | 0.120 | 0.250 | 0.112 | -0.127 | -0.181 | -0.281 | -0.157 | -0.001 | -0.180 | -0.025 |
| | P | 0.315 | 0.673 | 0.069 | 0.628 | 0.963 | 0.296 | 0.058 | 0.266 | 0.210 | 0.734 | 0.308 | 0.210 | 0.197 | 0.685 | **0.042** | 0.123 | 0.980 | 0.671 | 0.411 | 0.299 | 0.237 | 0.205 | 0.942 | 0.374 | 0.808 | 0.687 | 0.302 | 0.401 | 0.077 | 0.433 | 0.375 | 0.204 | **0.046** | 0.273 | 0.996 | 0.205 | 0.864 |
| CD86 | R | 0.099 | -0.010 | 0.387 | -0.027 | 0.242 | -0.385 | 0.247 | 0.161 | 0.068 | 0.128 | 0.094 | 0.234 | 0.253 | -0.087 | -0.082 | 0.133 | 0.095 | 0.089 | -0.225 | 0.253 | -0.019 | 0.383 | 0.005 | 0.139 | -0.024 | 0.070 | 0.314 | -0.240 | 0.074 | 0.117 | 0.098 | 0.209 | 0.192 | 0.271 | 0.053 | 0.208 | 0.177 |
| | P | 0.488 | 0.945 | **0.005** | 0.851 | 0.088 | **0.005** | 0.080 | 0.258 | 0.638 | 0.370 | 0.514 | 0.099 | 0.074 | 0.545 | 0.566 | 0.354 | 0.508 | 0.534 | 0.112 | 0.073 | 0.897 | **0.006** | 0.970 | 0.330 | 0.866 | 0.623 | **0.025** | 0.091 | 0.608 | 0.412 | 0.496 | 0.140 | 0.177 | 0.054 | 0.713 | 0.144 | 0.213 |
| CD105 | R | 0.106 | 0.026 | 0.425 | -0.083 | 0.158 | -0.286 | 0.193 | 0.156 | 0.090 | 0.129 | -0.006 | 0.277 | 0.291 | -0.097 | -0.087 | 0.156 | 0.052 | 0.023 | -0.214 | 0.261 | 0.011 | 0.405 | -0.026 | 0.216 | -0.114 | 0.020 | 0.265 | -0.226 | 0.053 | -0.034 | 0.156 | 0.244 | 0.130 | 0.145 | 0.181 | 0.249 | 0.140 |
| | P | 0.458 | 0.854 | **0.002** | 0.563 | 0.269 | **0.042** | 0.176 | 0.275 | 0.532 | 0.367 | 0.966 | **0.049** | **0.039** | 0.498 | 0.543 | 0.275 | 0.716 | 0.873 | 0.132 | 0.064 | 0.937 | **0.003** | 0.854 | 0.127 | 0.426 | 0.891 | 0.060 | 0.110 | 0.712 | 0.810 | 0.273 | 0.085 | 0.364 | 0.310 | 0.203 | 0.078 | 0.328 |
| CD133/1 | R | -0.041 | 0.030 | 0.324 | -0.122 | 0.037 | -0.253 | 0.079 | 0.189 | 0.004 | -0.041 | -0.014 | 0.110 | 0.076 | -0.063 | -0.072 | 0.035 | -0.136 | 0.050 | -0.080 | 0.041 | -0.119 | 0.250 | -0.022 | 0.070 | -0.015 | -0.099 | 0.151 | -0.291 | -0.044 | 0.085 | 0.198 | 0.044 | 0.060 | 0.095 | -0.007 | 0.071 | 0.028 |
| | P | 0.776 | 0.835 | **0.020** | 0.393 | 0.798 | 0.073 | 0.580 | 0.184 | 0.975 | 0.775 | 0.922 | 0.441 | 0.595 | 0.662 | 0.617 | 0.810 | 0.340 | 0.726 | 0.575 | 0.779 | 0.407 | 0.077 | 0.880 | 0.627 | 0.919 | 0.488 | 0.289 | **0.038** | 0.761 | 0.554 | 0.163 | 0.757 | 0.674 | 0.509 | 0.959 | 0.620 | 0.845 |
| CD142 | R | -0.043 | -0.136 | 0.297 | -0.024 | 0.180 | -0.261 | 0.103 | 0.114 | -0.005 | -0.027 | 0.068 | 0.106 | 0.140 | -0.053 | -0.285 | 0.030 | -0.029 | -0.013 | -0.244 | 0.092 | 0.022 | 0.227 | -0.104 | 0.070 | 0.009 | -0.041 | 0.160 | -0.124 | 0.086 | -0.106 | 0.137 | 0.076 | 0.040 | 0.144 | -0.103 | 0.078 | 0.020 |
| | P | 0.765 | 0.340 | **0.035** | 0.865 | 0.206 | 0.064 | 0.471 | 0.426 | 0.973 | 0.850 | 0.633 | 0.458 | 0.326 | 0.710 | **0.043** | 0.836 | 0.840 | 0.926 | 0.084 | 0.519 | 0.879 | 0.109 | 0.468 | 0.624 | 0.948 | 0.774 | 0.261 | 0.387 | 0.548 | 0.459 | 0.336 | 0.596 | 0.780 | 0.315 | 0.470 | 0.587 | 0.890 |
| CD146 | R | -0.088 | -0.057 | 0.336 | -0.109 | 0.025 | -0.147 | -0.086 | 0.023 | 0.013 | 0.063 | -0.028 | 0.130 | 0.078 | -0.060 | -0.077 | -0.015 | -0.126 | 0.081 | -0.078 | 0.042 | -0.064 | 0.147 | -0.062 | 0.149 | -0.010 | -0.090 | 0.109 | -0.125 | 0.096 | -0.107 | 0.154 | 0.012 | -0.100 | -0.017 | 0.015 | 0.064 | 0.082 |
| | P | 0.539 | 0.689 | **0.016** | 0.445 | 0.860 | 0.303 | 0.550 | 0.870 | 0.925 | 0.659 | 0.846 | 0.364 | 0.588 | 0.674 | 0.592 | 0.916 | 0.378 | 0.573 | 0.587 | 0.768 | 0.654 | 0.303 | 0.667 | 0.297 | 0.945 | 0.529 | 0.447 | 0.382 | 0.503 | 0.455 | 0.282 | 0.935 | 0.484 | 0.905 | 0.914 | 0.656 | 0.568 |
| CD209 | R | 0.058 | -0.110 | 0.459 | 0.008 | 0.139 | -0.221 | 0.166 | 0.188 | 0.120 | 0.039 | 0.055 | 0.237 | 0.244 | 0.028 | -0.185 | 0.163 | 0.029 | 0.046 | -0.189 | 0.216 | 0.043 | 0.327 | -0.058 | 0.172 | -0.099 | 0.034 | 0.305 | -0.159 | 0.035 | -0.094 | 0.098 | 0.193 | 0.126 | 0.153 | 0.023 | 0.202 | 0.121 |
| | P | 0.686 | 0.443 | **0.001** | 0.957 | 0.330 | 0.119 | 0.244 | 0.188 | 0.400 | 0.785 | 0.699 | 0.094 | 0.084 | 0.844 | 0.194 | 0.254 | 0.842 | 0.751 | 0.185 | 0.127 | 0.765 | **0.019** | 0.686 | 0.227 | 0.491 | 0.814 | **0.030** | 0.265 | 0.809 | 0.513 | 0.495 | 0.175 | 0.377 | 0.285 | 0.875 | 0.156 | 0.399 |
| CD326 | R | 0.054 | -0.081 | 0.343 | -0.254 | 0.212 | -0.358 | 0.119 | 0.077 | -0.031 | -0.022 | 0.046 | 0.117 | 0.084 | -0.085 | -0.078 | 0.082 | -0.027 | 0.121 | -0.005 | 0.027 | -0.167 | 0.147 | 0.003 | 0.006 | 0.145 | 0.021 | 0.162 | -0.122 | 0.051 | 0.053 | 0.118 | 0.042 | 0.162 | 0.169 | 0.005 | 0.060 | -0.062 |
| | P | 0.704 | 0.574 | **0.014** | 0.072 | 0.136 | **0.010** | 0.404 | 0.590 | 0.827 | 0.881 | 0.751 | 0.414 | 0.559 | 0.552 | 0.585 | 0.565 | 0.851 | 0.400 | 0.973 | 0.849 | 0.242 | 0.305 | 0.984 | 0.965 | 0.312 | 0.882 | 0.256 | 0.396 | 0.721 | 0.709 | 0.411 | 0.770 | 0.257 | 0.236 | 0.970 | 0.676 | 0.664 |
| HLA-I | R | 0.188 | 0.044 | 0.140 | 0.067 | 0.234 | -0.113 | 0.344 | 0.125 | 0.139 | 0.107 | 0.073 | 0.192 | 0.261 | -0.158 | -0.106 | 0.125 | 0.073 | -0.158 | -0.166 | 0.290 | 0.010 | 0.242 | 0.024 | 0.109 | -0.110 | 0.112 | 0.230 | -0.037 | -0.054 | 0.049 | 0.051 | 0.266 | 0.288 | 0.244 | 0.155 | 0.207 | 0.123 |
| | P | 0.187 | 0.760 | 0.329 | 0.642 | 0.098 | 0.428 | **0.014** | 0.384 | 0.330 | 0.455 | 0.611 | 0.177 | 0.064 | 0.268 | 0.458 | 0.383 | 0.612 | 0.267 | 0.245 | **0.039** | 0.944 | 0.087 | 0.865 | 0.447 | 0.443 | 0.435 | 0.104 | 0.797 | 0.709 | 0.734 | 0.724 | 0.059 | **0.040** | 0.084 | 0.277 | 0.145 | 0.391 |
| HLA-II | R | -0.111 | 0.081 | -0.041 | 0.024 | 0.079 | 0.083 | 0.086 | -0.135 | -0.058 | -0.063 | -0.018 | -0.128 | -0.099 | 0.170 | 0.026 | -0.020 | -0.191 | -0.124 | 0.088 | -0.034 | -0.035 | 0.117 | -0.167 | -0.149 | -0.153 | -0.061 | -0.096 | -0.132 | -0.021 | 0.157 | -0.057 | -0.045 | -0.031 | -0.051 | -0.095 | -0.096 | -0.035 |
| | P | 0.439 | 0.571 | 0.777 | 0.865 | 0.581 | 0.564 | 0.547 | 0.345 | 0.687 | 0.661 | 0.900 | 0.373 | 0.487 | 0.233 | 0.855 | 0.891 | 0.180 | 0.386 | 0.539 | 0.811 | 0.809 | 0.415 | 0.242 | 0.298 | 0.284 | 0.671 | 0.502 | 0.355 | 0.884 | 0.271 | 0.690 | 0.753 | 0.828 | 0.721 | 0.509 | 0.502 | 0.808 |
| MCSP | R | 0.014 | -0.016 | 0.325 | 0.053 | 0.073 | -0.253 | 0.049 | 0.168 | 0.029 | 0.018 | 0.072 | 0.156 | 0.155 | -0.016 | -0.125 | 0.071 | -0.063 | 0.123 | -0.164 | 0.139 | -0.050 | 0.348 | -0.030 | 0.105 | -0.092 | -0.104 | 0.168 | -0.269 | -0.043 | -0.027 | 0.131 | 0.115 | 0.026 | 0.067 | 0.015 | 0.135 | 0.079 |
| | P | 0.922 | 0.914 | **0.020** | 0.712 | 0.611 | 0.073 | 0.732 | 0.239 | 0.840 | 0.900 | 0.618 | 0.273 | 0.279 | 0.912 | 0.382 | 0.621 | 0.659 | 0.390 | 0.250 | 0.330 | 0.730 | **0.012** | 0.835 | 0.464 | 0.521 | 0.466 | 0.238 | 0.056 | 0.767 | 0.849 | 0.359 | 0.420 | 0.854 | 0.642 | 0.918 | 0.345 | 0.582 |
| ROR1 | R | -0.043 | 0.017 | 0.187 | -0.129 | 0.017 | -0.193 | 0.091 | 0.095 | -0.059 | -0.100 | -0.035 | 0.039 | 0.037 | -0.033 | -0.071 | 0.018 | -0.144 | 0.012 | -0.109 | 0.028 | -0.147 | 0.240 | 0.048 | -0.032 | -0.050 | -0.159 | 0.055 | -0.304 | -0.134 | 0.067 | 0.141 | 0.022 | 0.105 | 0.135 | 0.008 | 0.020 | -0.041 |
| | P | 0.764 | 0.907 | 0.188 | 0.369 | 0.905 | 0.176 | 0.524 | 0.506 | 0.682 | 0.487 | 0.807 | 0.784 | 0.797 | 0.819 | 0.620 | 0.899 | 0.313 | 0.936 | 0.448 | 0.847 | 0.305 | 0.090 | 0.737 | 0.826 | 0.725 | 0.265 | 0.699 | **0.030** | 0.347 | 0.642 | 0.324 | 0.878 | 0.463 | 0.347 | 0.957 | 0.890 | 0.773 |
| SSEA4 | R | -0.121 | 0.010 | -0.015 | -0.049 | 0.028 | -0.084 | 0.181 | -0.028 | -0.078 | -0.101 | 0.041 | -0.130 | -0.107 | 0.054 | -0.072 | -0.086 | -0.147 | -0.122 | -0.078 | -0.069 | -0.042 | 0.066 | -0.091 | -0.152 | 0.009 | -0.057 | 0.039 | -0.302 | -0.028 | 0.321 | 0.026 | -0.093 | 0.007 | 0.229 | -0.217 | -0.121 | -0.034 |
| | P | 0.398 | 0.942 | 0.918 | 0.733 | 0.845 | 0.557 | 0.203 | 0.846 | 0.585 | 0.480 | 0.776 | 0.362 | 0.453 | 0.709 | 0.618 | 0.548 | 0.303 | 0.395 | 0.586 | 0.633 | 0.768 | 0.645 | 0.525 | 0.286 | 0.952 | 0.689 | 0.786 | **0.031** | 0.848 | **0.022** | 0.855 | 0.514 | 0.959 | 0.106 | 0.126 | 0.398 | 0.812 |

| Model ID | Oversampling | Trees (n) | Leaves (n) | Sensitivity (%) | Specificity (%) | Accuracy (%) | Macro average accuracy (%) |
|---|---|---|---|---|---|---|---|
| **1** | SMOTE | 10 | 10 | 76.1 | 78.7 | 77.8 | 77.4 |
| **2** | SMOTE | 10 | 20 | 76.1 | 78.7 | 77.8 | 77.4 |
| **3** | SMOTE | 10 | 40 | 76.1 | 78.7 | 77.8 | 77.4 |
| **4** | SMOTE | 10 | 80 | 76.1 | 78.7 | 77.8 | 77.4 |
| **5** | SMOTE | 10 | 160 | 76.1 | 78.7 | 77.8 | 77.4 |
| **6** | SMOTE | 10 | 320 | 76.1 | 78.7 | 77.8 | 77.4 |
| **7** | SMOTE | 50 | 10 | 71.4 | 78.8 | 75.9 | 75.1 |
| **8** | SMOTE | 50 | 20 | 71.4 | 78.8 | 75.9 | 75.1 |
| **9** | SMOTE | 50 | 40 | 71.4 | 78.8 | 75.9 | 75.1 |
| **10** | SMOTE | 50 | 80 | 71.4 | 78.8 | 75.9 | 75.1 |
| **11** | SMOTE | 50 | 160 | 71.4 | 78.8 | 75.9 | 75.1 |
| **12** | SMOTE | 50 | 320 | 71.4 | 78.8 | 75.9 | 75.1 |
| **13** | SMOTE | 100 | 10 | 71.4 | 75.8 | 74.1 | 73.6 |
| **14** | SMOTE | 100 | 20 | 71.4 | 75.8 | 74.1 | 73.6 |
| **15** | SMOTE | 100 | 40 | 71.4 | 75.8 | 74.1 | 73.6 |
| **16** | SMOTE | 100 | 80 | 71.4 | 75.8 | 74.1 | 73.6 |
| **17** | SMOTE | 100 | 160 | 71.4 | 75.8 | 74.1 | 73.6 |
| **18** | SMOTE | 100 | 320 | 71.4 | 75.8 | 74.1 | 73.6 |
| **19** | SMOTE | 200 | 10 | 71.4 | 75.8 | 74.1 | 73.6 |
| **20** | SMOTE | 200 | 20 | 71.4 | 75.8 | 74.1 | 73.6 |
| **21** | SMOTE | 200 | 40 | 71.4 | 75.8 | 74.1 | 73.6 |
| **22** | SMOTE | 200 | 80 | 71.4 | 75.8 | 74.1 | 73.6 |
| **23** | SMOTE | 200 | 160 | 71.4 | 75.8 | 74.1 | 73.6 |
| **24** | SMOTE | 200 | 320 | 71.4 | 75.8 | 74.1 | 73.6 |
| **25** | SMOTEENN | 10 | 10 | 57.1 | 93.9 | 79.6 | 75.5 |
| **26** | SMOTEENN | 10 | 20 | 57.1 | 93.9 | 79.6 | 75.5 |
| **27** | SMOTEENN | 10 | 40 | 57.1 | 93.9 | 79.6 | 75.5 |
| **28** | SMOTEENN | 10 | 80 | 57.1 | 93.9 | 79.6 | 75.5 |
| **29** | SMOTEENN | 10 | 160 | 57.1 | 93.9 | 79.6 | 75.5 |
| **30** | SMOTEENN | 10 | 320 | 57.1 | 93.9 | 79.6 | 75.5 |
| **31** | SMOTEENN | 50 | 10 | 61.9 | 78.8 | 72.2 | 70.3 |
| **32** | SMOTEENN | 50 | 20 | 61.9 | 78.8 | 72.2 | 70.3 |
| **33** | SMOTEENN | 50 | 40 | 61.9 | 78.8 | 72.2 | 70.3 |
| **34** | SMOTEENN | 50 | 80 | 61.9 | 78.8 | 72.2 | 70.3 |
| **35** | SMOTEENN | 50 | 160 | 61.9 | 78.8 | 72.2 | 70.3 |
| **36** | SMOTEENN | 50 | 320 | 61.9 | 78.8 | 72.2 | 70.3 |
| **37** | SMOTEENN | 100 | 10 | 61.9 | 78.8 | 72.2 | 70.3 |
| **38** | SMOTEENN | 100 | 20 | 61.9 | 78.8 | 72.2 | 70.3 |
| **39** | SMOTEENN | 100 | 40 | 61.9 | 78.8 | 72.2 | 70.3 |
| **40** | SMOTEENN | 100 | 80 | 61.9 | 78.8 | 72.2 | 70.3 |
| **41** | SMOTEENN | 100 | 160 | 61.9 | 78.8 | 72.2 | 70.3 |
| **42** | SMOTEENN | 100 | 320 | 61.9 | 78.8 | 72.2 | 70.3 |
| **43** | SMOTEENN | 200 | 10 | 66.7 | 78.8 | 74.1 | 72.7 |
| **44** | SMOTEENN | 200 | 20 | 66.7 | 78.8 | 74.1 | 72.7 |
| **45** | SMOTEENN | 200 | 40 | 66.7 | 78.8 | 74.1 | 72.7 |
| **46** | SMOTEENN | 200 | 80 | 66.7 | 78.8 | 74.1 | 72.7 |
| **47** | SMOTEENN | 200 | 160 | 66.7 | 78.8 | 74.1 | 72.7 |
| **48** | SMOTEENN | 200 | 320 | 66.7 | 78.8 | 74.1 | 72.7 |
| **49** | RO | 10 | 10 | 76.2 | 72.7 | 74.1 | 74.5 |
| **50** | RO | 10 | 20 | 76.2 | 72.7 | 74.1 | 74.5 |
| **51** | RO | 10 | 40 | 76.2 | 72.7 | 74.1 | 74.5 |
| **52** | RO | 10 | 80 | 76.2 | 72.7 | 74.1 | 74.5 |
| **53** | RO | 10 | 160 | 76.2 | 72.7 | 74.1 | 74.5 |
| **54** | RO | 10 | 320 | 76.2 | 72.7 | 74.1 | 74.5 |
| **55** | RO | 50 | 10 | 71.4 | 75.8 | 74.1 | 73.6 |
| **56** | RO | 50 | 20 | 71.4 | 75.8 | 74.1 | 73.6 |
| **57** | RO | 50 | 40 | 71.4 | 75.8 | 74.1 | 73.6 |
| **58** | RO | 50 | 80 | 71.4 | 75.8 | 74.1 | 73.6 |
| **59** | RO | 50 | 160 | 71.4 | 75.8 | 74.1 | 73.6 |
| **60** | RO | 50 | 320 | 71.4 | 75.8 | 74.1 | 73.6 |
| **61** | RO | 100 | 10 | 71.4 | 72.7 | 72.2 | 72.1 |
| **62** | RO | 100 | 20 | 71.4 | 72.7 | 72.2 | 72.1 |
| **63** | RO | 100 | 40 | 71.4 | 72.7 | 72.2 | 72.1 |
| **64** | RO | 100 | 80 | 71.4 | 72.7 | 72.2 | 72.1 |
| **65** | RO | 100 | 160 | 71.4 | 72.7 | 72.2 | 72.1 |
| **66** | RO | 100 | 320 | 71.4 | 72.7 | 72.2 | 72.1 |
| **67** | RO | 200 | 10 | 71.4 | 72.7 | 72.2 | 72.1 |
| **68** | RO | 200 | 20 | 71.4 | 72.7 | 72.2 | 72.1 |
| **69** | RO | 200 | 40 | 71.4 | 72.7 | 72.2 | 72.1 |
| **70** | RO | 200 | 80 | 71.4 | 72.7 | 72.2 | 72.1 |
| **71** | RO | 200 | 160 | 71.4 | 72.7 | 72.2 | 72.1 |
| **72** | RO | 200 | 320 | 71.4 | 72.7 | 72.2 | 72.1 |
| **73** | None | 10 | 10 | 61.9 | 81.8 | 74.1 | 71.9 |
| **74** | None | 10 | 20 | 61.9 | 81.8 | 74.1 | 71.9 |
| **75** | None | 10 | 40 | 61.9 | 81.8 | 74.1 | 71.9 |
| **76** | None | 10 | 80 | 61.9 | 81.8 | 74.1 | 71.9 |
| **77** | None | 10 | 160 | 61.9 | 81.8 | 74.1 | 71.9 |
| **78** | None | 10 | 320 | 61.9 | 81.8 | 74.1 | 71.9 |
| **79** | None | 50 | 10 | 61.9 | 81.8 | 74.1 | 71.9 |
| **80** | None | 50 | 20 | 61.9 | 81.8 | 74.1 | 71.9 |
| **81** | None | 50 | 40 | 61.9 | 81.8 | 74.1 | 71.9 |
| **82** | None | 50 | 80 | 61.9 | 81.8 | 74.1 | 71.9 |
| **83** | None | 50 | 160 | 61.9 | 81.8 | 74.1 | 71.9 |
| **84** | None | 50 | 320 | 61.9 | 81.8 | 74.1 | 71.9 |
| **85** | None | 100 | 10 | 66.7 | 81.8 | 75.9 | 74.2 |
| **86** | None | 100 | 20 | 66.7 | 81.8 | 75.9 | 74.2 |
| **87** | None | 100 | 40 | 66.7 | 81.8 | 75.9 | 74.2 |
| **88** | None | 100 | 80 | 66.7 | 81.8 | 75.9 | 74.2 |
| **89** | None | 100 | 160 | 66.7 | 81.8 | 75.9 | 74.2 |
| **90** | None | 100 | 320 | 66.7 | 81.8 | 75.9 | 74.2 |
| **91** | None | 200 | 10 | 52.4 | 81.8 | 70.4 | 67.1 |
| **92** | None | 200 | 20 | 52.4 | 81.8 | 70.4 | 67.1 |
| **93** | None | 200 | 40 | 52.4 | 81.8 | 70.4 | 67.1 |
| **94** | None | 200 | 80 | 52.4 | 81.8 | 70.4 | 67.1 |
| **95** | None | 200 | 160 | 52.4 | 81.8 | 70.4 | 67.1 |
| **96** | None | 200 | 320 | 52.4 | 81.8 | 70.4 | 67.1 |

| Model ID | Oversampling | Trees (n) | Leaves (n) | Sensitivity (%) | Specificity (%) | Accuracy (%) | Macro average accuracy (%) |
|---|---|---|---|---|---|---|---|
| **1** | SMOTE | 10 | 10 | 57.1 | 78.8 | 72.3 | 68.0 |
| **2** | SMOTE | 10 | 20 | 57.1 | 78.8 | 72.3 | 68.0 |
| **3** | SMOTE | 10 | 40 | 57.1 | 78.8 | 72.3 | 68.0 |
| **4** | SMOTE | 10 | 80 | 57.1 | 78.8 | 72.3 | 68.0 |
| **5** | SMOTE | 10 | 160 | 57.1 | 78.8 | 72.3 | 68.0 |
| **6** | SMOTE | 10 | 320 | 57.1 | 78.8 | 72.3 | 68.0 |
| **7** | SMOTE | 50 | 10 | 57.1 | 84.8 | 76.6 | 71.0 |
| **8** | SMOTE | 50 | 20 | 57.1 | 84.8 | 76.6 | 71.0 |
| **9** | SMOTE | 50 | 40 | 57.1 | 84.8 | 76.6 | 71.0 |
| **10** | SMOTE | 50 | 80 | 57.1 | 84.8 | 76.6 | 71.0 |
| **11** | SMOTE | 50 | 160 | 57.1 | 84.8 | 76.6 | 71.0 |
| **12** | SMOTE | 50 | 320 | 57.1 | 84.8 | 76.6 | 71.0 |
| **13** | SMOTE | 100 | 10 | 57.1 | 81.8 | 74.5 | 69.5 |
| **14** | SMOTE | 100 | 20 | 57.1 | 81.8 | 74.5 | 69.5 |
| **15** | SMOTE | 100 | 40 | 57.1 | 81.8 | 74.5 | 69.5 |
| **16** | SMOTE | 100 | 80 | 57.1 | 81.8 | 74.5 | 69.5 |
| **17** | SMOTE | 100 | 160 | 57.1 | 81.8 | 74.5 | 69.5 |
| **18** | SMOTE | 100 | 320 | 57.1 | 81.8 | 74.5 | 69.5 |
| **19** | SMOTE | 200 | 10 | 57.1 | 78.8 | 72.3 | 68.0 |
| **20** | SMOTE | 200 | 20 | 57.1 | 78.8 | 72.3 | 68.0 |
| **21** | SMOTE | 200 | 40 | 57.1 | 78.8 | 72.3 | 68.0 |
| **22** | SMOTE | 200 | 80 | 57.1 | 78.8 | 72.3 | 68.0 |
| **23** | SMOTE | 200 | 160 | 57.1 | 78.8 | 72.3 | 68.0 |
| **24** | SMOTE | 200 | 320 | 57.1 | 78.8 | 72.3 | 68.0 |
| **25** | SMOTEENN | 10 | 10 | 64.3 | 54.5 | 57.4 | 59.4 |
| **26** | SMOTEENN | 10 | 20 | 64.3 | 54.5 | 57.4 | 59.4 |
| **27** | SMOTEENN | 10 | 40 | 64.3 | 54.5 | 57.4 | 59.4 |
| **28** | SMOTEENN | 10 | 80 | 64.3 | 54.5 | 57.4 | 59.4 |
| **29** | SMOTEENN | 10 | 160 | 64.3 | 54.5 | 57.4 | 59.4 |
| **30** | SMOTEENN | 10 | 320 | 64.3 | 54.5 | 57.4 | 59.4 |
| **31** | SMOTEENN | 50 | 10 | 64.3 | 69.7 | 68.1 | 67.0 |
| **32** | SMOTEENN | 50 | 20 | 64.3 | 69.7 | 68.1 | 67.0 |
| **33** | SMOTEENN | 50 | 40 | 64.3 | 69.7 | 68.1 | 67.0 |
| **34** | SMOTEENN | 50 | 80 | 64.3 | 69.7 | 68.1 | 67.0 |
| **35** | SMOTEENN | 50 | 160 | 64.3 | 69.7 | 68.1 | 67.0 |
| **36** | SMOTEENN | 50 | 320 | 64.3 | 69.7 | 68.1 | 67.0 |
| **37** | SMOTEENN | 100 | 10 | 64.3 | 66.7 | 66.0 | 65.5 |
| **38** | SMOTEENN | 100 | 20 | 64.3 | 66.7 | 66.0 | 65.5 |
| **39** | SMOTEENN | 100 | 40 | 64.3 | 66.7 | 66.0 | 65.5 |
| **40** | SMOTEENN | 100 | 80 | 64.3 | 66.7 | 66.0 | 65.5 |
| **41** | SMOTEENN | 100 | 160 | 64.3 | 66.7 | 66.0 | 65.5 |
| **42** | SMOTEENN | 100 | 320 | 64.3 | 66.7 | 66.0 | 65.5 |
| **43** | SMOTEENN | 200 | 10 | 71.4 | 66.7 | 68.1 | 69.0 |
| **44** | SMOTEENN | 200 | 20 | 71.4 | 66.7 | 68.1 | 69.0 |
| **45** | SMOTEENN | 200 | 40 | 71.4 | 66.7 | 68.1 | 69.0 |
| **46** | SMOTEENN | 200 | 80 | 71.4 | 66.7 | 68.1 | 69.0 |
| **47** | SMOTEENN | 200 | 160 | 71.4 | 66.7 | 68.1 | 69.0 |
| **48** | SMOTEENN | 200 | 320 | 71.4 | 66.7 | 68.1 | 69.0 |
| **49** | RO | 10 | 10 | 64.3 | 66.7 | 66.0 | 65.5 |
| **50** | RO | 10 | 20 | 64.3 | 66.7 | 66.0 | 65.5 |
| **51** | RO | 10 | 40 | 64.3 | 66.7 | 66.0 | 65.5 |
| **52** | RO | 10 | 80 | 64.3 | 66.7 | 66.0 | 65.5 |
| **53** | RO | 10 | 160 | 64.3 | 66.7 | 66.0 | 65.5 |
| **54** | RO | 10 | 320 | 64.3 | 66.7 | 66.0 | 65.5 |
| **55** | RO | 50 | 10 | 57.1 | 72.7 | 68.1 | 64.9 |
| **56** | RO | 50 | 20 | 57.1 | 72.7 | 68.1 | 64.9 |
| **57** | RO | 50 | 40 | 57.1 | 72.7 | 68.1 | 64.9 |
| **58** | RO | 50 | 80 | 57.1 | 72.7 | 68.1 | 64.9 |
| **59** | RO | 50 | 160 | 57.1 | 72.7 | 68.1 | 64.9 |
| **60** | RO | 50 | 320 | 57.1 | 72.7 | 68.1 | 64.9 |
| **61** | RO | 100 | 10 | 71.1 | 70.7 | 70.8 | 70.9 |
| **62** | RO | 100 | 20 | 71.1 | 70.7 | 70.8 | 70.9 |
| **63** | RO | 100 | 40 | 71.1 | 70.7 | 70.8 | 70.9 |
| **64** | RO | 100 | 80 | 71.1 | 70.7 | 70.8 | 70.9 |
| **65** | RO | 100 | 160 | 71.1 | 70.7 | 70.8 | 70.9 |
| **66** | RO | 100 | 320 | 71.1 | 70.7 | 70.8 | 70.9 |
| **67** | RO | 200 | 10 | 71.1 | 70.7 | 70.8 | 70.9 |
| **68** | RO | 200 | 20 | 71.1 | 70.7 | 70.8 | 70.9 |
| **69** | RO | 200 | 40 | 71.1 | 70.7 | 70.8 | 70.9 |
| **70** | RO | 200 | 80 | 71.1 | 70.7 | 70.8 | 70.9 |
| **71** | RO | 200 | 160 | 71.1 | 70.7 | 70.8 | 70.9 |
| **72** | RO | 200 | 320 | 71.1 | 70.7 | 70.8 | 70.9 |
| **73** | None | 10 | 10 | 21.4 | 81.8 | 63.8 | 51.6 |
| **74** | None | 10 | 20 | 21.4 | 81.8 | 63.8 | 51.6 |
| **75** | None | 10 | 40 | 21.4 | 81.8 | 63.8 | 51.6 |
| **76** | None | 10 | 80 | 21.4 | 81.8 | 63.8 | 51.6 |
| **77** | None | 10 | 160 | 21.4 | 81.8 | 63.8 | 51.6 |
| **78** | None | 10 | 320 | 21.4 | 81.8 | 63.8 | 51.6 |
| **79** | None | 50 | 10 | 42.9 | 84.8 | 72.3 | 63.9 |
| **80** | None | 50 | 20 | 42.9 | 84.8 | 72.3 | 63.9 |
| **81** | None | 50 | 40 | 42.9 | 84.8 | 72.3 | 63.9 |
| **82** | None | 50 | 80 | 42.9 | 84.8 | 72.3 | 63.9 |
| **83** | None | 50 | 160 | 42.9 | 84.8 | 72.3 | 63.9 |
| **84** | None | 50 | 320 | 42.9 | 84.8 | 72.3 | 63.9 |
| **85** | None | 100 | 10 | 35.7 | 90.9 | 74.5 | 63.3 |
| **86** | None | 100 | 20 | 35.7 | 90.9 | 74.5 | 63.3 |
| **87** | None | 100 | 40 | 35.7 | 90.9 | 74.5 | 63.3 |
| **88** | None | 100 | 80 | 35.7 | 90.9 | 74.5 | 63.3 |
| **89** | None | 100 | 160 | 35.7 | 90.9 | 74.5 | 63.3 |
| **90** | None | 100 | 320 | 35.7 | 90.9 | 74.5 | 63.3 |
| **91** | None | 200 | 10 | 42.9 | 90.9 | 76.6 | 66.9 |
| **92** | None | 200 | 20 | 42.9 | 90.9 | 76.6 | 66.9 |
| **93** | None | 200 | 40 | 42.9 | 90.9 | 76.6 | 66.9 |
| **94** | None | 200 | 80 | 42.9 | 90.9 | 76.6 | 66.9 |
| **95** | None | 200 | 160 | 42.9 | 90.9 | 76.6 | 66.9 |
| **96** | None | 200 | 320 | 42.9 | 90.9 | 76.6 | 66.9 |

| Model ID | Oversampling | Trees (n) | Leaves (n) | Sensitivity (%) | Specificity (%) | Accuracy (%) | Macro average accuracy (%) |
|---|---|---|---|---|---|---|---|
| **1** | SMOTE | 10 | 10 | 62.1 | 63.6 | 62.9 | 62.9 |
| **2** | SMOTE | 10 | 20 | 62.1 | 63.6 | 62.9 | 62.9 |
| **3** | SMOTE | 10 | 40 | 62.1 | 63.6 | 62.9 | 62.9 |
| **4** | SMOTE | 10 | 80 | 62.1 | 63.6 | 62.9 | 62.9 |
| **5** | SMOTE | 10 | 160 | 62.1 | 63.6 | 62.9 | 62.9 |
| **6** | SMOTE | 10 | 320 | 62.1 | 63.6 | 62.9 | 62.9 |
| **7** | SMOTE | 50 | 10 | 69.0 | 60.6 | 64.5 | 64.8 |
| **8** | SMOTE | 50 | 20 | 69.0 | 60.6 | 64.5 | 64.8 |
| **9** | SMOTE | 50 | 40 | 69.0 | 60.6 | 64.5 | 64.8 |
| **10** | SMOTE | 50 | 80 | 69.0 | 60.6 | 64.5 | 64.8 |
| **11** | SMOTE | 50 | 160 | 69.0 | 60.6 | 64.5 | 64.8 |
| **12** | SMOTE | 50 | 320 | 69.0 | 60.6 | 64.5 | 64.8 |
| **13** | SMOTE | 100 | 10 | 69.0 | 60.6 | 64.5 | 64.8 |
| **14** | SMOTE | 100 | 20 | 69.0 | 60.6 | 64.5 | 64.8 |
| **15** | SMOTE | 100 | 40 | 69.0 | 60.6 | 64.5 | 64.8 |
| **16** | SMOTE | 100 | 80 | 69.0 | 60.6 | 64.5 | 64.8 |
| **17** | SMOTE | 100 | 160 | 69.0 | 60.6 | 64.5 | 64.8 |
| **18** | SMOTE | 100 | 320 | 69.0 | 60.6 | 64.5 | 64.8 |
| **19** | SMOTE | 200 | 10 | 69.0 | 57.6 | 62.9 | 63.3 |
| **20** | SMOTE | 200 | 20 | 69.0 | 57.6 | 62.9 | 63.3 |
| **21** | SMOTE | 200 | 40 | 69.0 | 57.6 | 62.9 | 63.3 |
| **22** | SMOTE | 200 | 80 | 69.0 | 57.6 | 62.9 | 63.3 |
| **23** | SMOTE | 200 | 160 | 69.0 | 57.6 | 62.9 | 63.3 |
| **24** | SMOTE | 200 | 320 | 69.0 | 57.6 | 62.9 | 63.3 |
| **25** | SMOTEENN | 10 | 10 | 72.4 | 45.5 | 58.1 | 58.9 |
| **26** | SMOTEENN | 10 | 20 | 72.4 | 45.5 | 58.1 | 58.9 |
| **27** | SMOTEENN | 10 | 40 | 72.4 | 45.5 | 58.1 | 58.9 |
| **28** | SMOTEENN | 10 | 80 | 72.4 | 45.5 | 58.1 | 58.9 |
| **29** | SMOTEENN | 10 | 160 | 72.4 | 45.5 | 58.1 | 58.9 |
| **30** | SMOTEENN | 10 | 320 | 72.4 | 45.5 | 58.1 | 58.9 |
| **31** | SMOTEENN | 50 | 10 | 79.3 | 48.5 | 62.9 | 63.9 |
| **32** | SMOTEENN | 50 | 20 | 79.3 | 48.5 | 62.9 | 63.9 |
| **33** | SMOTEENN | 50 | 40 | 79.3 | 48.5 | 62.9 | 63.9 |
| **34** | SMOTEENN | 50 | 80 | 79.3 | 48.5 | 62.9 | 63.9 |
| **35** | SMOTEENN | 50 | 160 | 79.3 | 48.5 | 62.9 | 63.9 |
| **36** | SMOTEENN | 50 | 320 | 79.3 | 48.5 | 62.9 | 63.9 |
| **37** | SMOTEENN | 100 | 10 | 82.8 | 48.5 | 64.5 | 65.6 |
| **38** | SMOTEENN | 100 | 20 | 82.8 | 48.5 | 64.5 | 65.6 |
| **39** | SMOTEENN | 100 | 40 | 82.8 | 48.5 | 64.5 | 65.6 |
| **40** | SMOTEENN | 100 | 80 | 82.8 | 48.5 | 64.5 | 65.6 |
| **41** | SMOTEENN | 100 | 160 | 82.8 | 48.5 | 64.5 | 65.6 |
| **42** | SMOTEENN | 100 | 320 | 82.8 | 48.5 | 64.5 | 65.6 |
| **43** | SMOTEENN | 200 | 10 | 82.8 | 48.5 | 64.5 | 65.6 |
| **44** | SMOTEENN | 200 | 20 | 82.8 | 48.5 | 64.5 | 65.6 |
| **45** | SMOTEENN | 200 | 40 | 82.8 | 48.5 | 64.5 | 65.6 |
| **46** | SMOTEENN | 200 | 80 | 82.8 | 48.5 | 64.5 | 65.6 |
| **47** | SMOTEENN | 200 | 160 | 82.8 | 48.5 | 64.5 | 65.6 |
| **48** | SMOTEENN | 200 | 320 | 82.8 | 48.5 | 64.5 | 65.6 |
| **49** | RO | 10 | 10 | 72.4 | 57.6 | 64.5 | 65.0 |
| **50** | RO | 10 | 20 | 72.4 | 57.6 | 64.5 | 65.0 |
| **51** | RO | 10 | 40 | 72.4 | 57.6 | 64.5 | 65.0 |
| **52** | RO | 10 | 80 | 72.4 | 57.6 | 64.5 | 65.0 |
| **53** | RO | 10 | 160 | 72.4 | 57.6 | 64.5 | 65.0 |
| **54** | RO | 10 | 320 | 72.4 | 57.6 | 64.5 | 65.0 |
| **55** | RO | 50 | 10 | 72.4 | 60.0 | 66.1 | 66.2 |
| **56** | RO | 50 | 20 | 72.4 | 60.0 | 66.1 | 66.2 |
| **57** | RO | 50 | 40 | 72.4 | 60.0 | 66.1 | 66.2 |
| **58** | RO | 50 | 80 | 72.4 | 60.0 | 66.1 | 66.2 |
| **59** | RO | 50 | 160 | 72.4 | 60.0 | 66.1 | 66.2 |
| **60** | RO | 50 | 320 | 72.4 | 60.0 | 66.1 | 66.2 |
| **61** | RO | 100 | 10 | 72.4 | 57.6 | 64.5 | 65.0 |
| **62** | RO | 100 | 20 | 72.4 | 57.6 | 64.5 | 65.0 |
| **63** | RO | 100 | 40 | 72.4 | 57.6 | 64.5 | 65.0 |
| **64** | RO | 100 | 80 | 72.4 | 57.6 | 64.5 | 65.0 |
| **65** | RO | 100 | 160 | 72.4 | 57.6 | 64.5 | 65.0 |
| **66** | RO | 100 | 320 | 72.4 | 57.6 | 64.5 | 65.0 |
| **67** | RO | 200 | 10 | 72.4 | 57.6 | 64.5 | 65.0 |
| **68** | RO | 200 | 20 | 72.4 | 57.6 | 64.5 | 65.0 |
| **69** | RO | 200 | 40 | 72.4 | 57.6 | 64.5 | 65.0 |
| **70** | RO | 200 | 80 | 72.4 | 57.6 | 64.5 | 65.0 |
| **71** | RO | 200 | 160 | 72.4 | 57.6 | 64.5 | 65.0 |
| **72** | RO | 200 | 320 | 72.4 | 57.6 | 64.5 | 65.0 |
| **73** | None | 10 | 10 | 69.0 | 60.6 | 64.5 | 64.8 |
| **74** | None | 10 | 20 | 69.0 | 60.6 | 64.5 | 64.8 |
| **75** | None | 10 | 40 | 69.0 | 60.6 | 64.5 | 64.8 |
| **76** | None | 10 | 80 | 69.0 | 60.6 | 64.5 | 64.8 |
| **77** | None | 10 | 160 | 69.0 | 60.6 | 64.5 | 64.8 |
| **78** | None | 10 | 320 | 69.0 | 60.6 | 64.5 | 64.8 |
| **79** | None | 50 | 10 | 62.1 | 54.5 | 58.1 | 58.3 |
| **80** | None | 50 | 20 | 62.1 | 54.5 | 58.1 | 58.3 |
| **81** | None | 50 | 40 | 62.1 | 54.5 | 58.1 | 58.3 |
| **82** | None | 50 | 80 | 62.1 | 54.5 | 58.1 | 58.3 |
| **83** | None | 50 | 160 | 62.1 | 54.5 | 58.1 | 58.3 |
| **84** | None | 50 | 320 | 62.1 | 54.5 | 58.1 | 58.3 |
| **85** | None | 100 | 10 | 69.0 | 54.5 | 61.3 | 61.8 |
| **86** | None | 100 | 20 | 69.0 | 54.5 | 61.3 | 61.8 |
| **87** | None | 100 | 40 | 69.0 | 54.5 | 61.3 | 61.8 |
| **88** | None | 100 | 80 | 69.0 | 54.5 | 61.3 | 61.8 |
| **89** | None | 100 | 160 | 69.0 | 54.5 | 61.3 | 61.8 |
| **90** | None | 100 | 320 | 69.0 | 54.5 | 61.3 | 61.8 |
| **91** | None | 200 | 10 | 65.5 | 54.5 | 59.7 | 60.0 |
| **92** | None | 200 | 20 | 65.5 | 54.5 | 59.7 | 60.0 |
| **93** | None | 200 | 40 | 65.5 | 54.5 | 59.7 | 60.0 |
| **94** | None | 200 | 80 | 65.5 | 54.5 | 59.7 | 60.0 |
| **95** | None | 200 | 160 | 65.5 | 54.5 | 59.7 | 60.0 |
| **96** | None | 200 | 320 | 65.5 | 54.5 | 59.7 | 60.0 |

| Model ID | Oversampling | Trees (n) | Leaves (n) | Sensitivity (%) | Specificity (%) | Accuracy (%) | Macro average accuracy (%) |
|---|---|---|---|---|---|---|---|
| **1** | SMOTE | 10 | 10 | 54.5 | 69.7 | 65.9 | 62.1 |
| **2** | SMOTE | 10 | 20 | 54.5 | 69.7 | 65.9 | 62.1 |
| **3** | SMOTE | 10 | 40 | 54.5 | 69.7 | 65.9 | 62.1 |
| **4** | SMOTE | 10 | 80 | 54.5 | 69.7 | 65.9 | 62.1 |
| **5** | SMOTE | 10 | 160 | 54.5 | 69.7 | 65.9 | 62.1 |
| **6** | SMOTE | 10 | 320 | 54.5 | 69.7 | 65.9 | 62.1 |
| **7** | SMOTE | 50 | 10 | 27.3 | 78.8 | 65.9 | 53.0 |
| **8** | SMOTE | 50 | 20 | 27.3 | 78.8 | 65.9 | 53.0 |
| **9** | SMOTE | 50 | 40 | 27.3 | 78.8 | 65.9 | 53.0 |
| **10** | SMOTE | 50 | 80 | 27.3 | 78.8 | 65.9 | 53.0 |
| **11** | SMOTE | 50 | 160 | 27.3 | 78.8 | 65.9 | 53.0 |
| **12** | SMOTE | 50 | 320 | 27.3 | 78.8 | 65.9 | 53.0 |
| **13** | SMOTE | 100 | 10 | 27.3 | 78.8 | 65.9 | 53.0 |
| **14** | SMOTE | 100 | 20 | 27.3 | 78.8 | 65.9 | 53.0 |
| **15** | SMOTE | 100 | 40 | 27.3 | 78.8 | 65.9 | 53.0 |
| **16** | SMOTE | 100 | 80 | 27.3 | 78.8 | 65.9 | 53.0 |
| **17** | SMOTE | 100 | 160 | 27.3 | 78.8 | 65.9 | 53.0 |
| **18** | SMOTE | 100 | 320 | 27.3 | 78.8 | 65.9 | 53.0 |
| **19** | SMOTE | 200 | 10 | 27.3 | 78.8 | 65.9 | 53.0 |
| **20** | SMOTE | 200 | 20 | 27.3 | 78.8 | 65.9 | 53.0 |
| **21** | SMOTE | 200 | 40 | 27.3 | 78.8 | 65.9 | 53.0 |
| **22** | SMOTE | 200 | 80 | 27.3 | 78.8 | 65.9 | 53.0 |
| **23** | SMOTE | 200 | 160 | 27.3 | 78.8 | 65.9 | 53.0 |
| **24** | SMOTE | 200 | 320 | 27.3 | 78.8 | 65.9 | 53.0 |
| **25** | SMOTEENN | 10 | 10 | 54.5 | 45.5 | 47.7 | 50.0 |
| **26** | SMOTEENN | 10 | 20 | 54.5 | 45.5 | 47.7 | 50.0 |
| **27** | SMOTEENN | 10 | 40 | 54.5 | 45.5 | 47.7 | 50.0 |
| **28** | SMOTEENN | 10 | 80 | 54.5 | 45.5 | 47.7 | 50.0 |
| **29** | SMOTEENN | 10 | 160 | 54.5 | 45.5 | 47.7 | 50.0 |
| **30** | SMOTEENN | 10 | 320 | 54.5 | 45.5 | 47.7 | 50.0 |
| **31** | SMOTEENN | 50 | 10 | 36.4 | 57.6 | 52.3 | 47.0 |
| **32** | SMOTEENN | 50 | 20 | 36.4 | 57.6 | 52.3 | 47.0 |
| **33** | SMOTEENN | 50 | 40 | 36.4 | 57.6 | 52.3 | 47.0 |
| **34** | SMOTEENN | 50 | 80 | 36.4 | 57.6 | 52.3 | 47.0 |
| **35** | SMOTEENN | 50 | 160 | 36.4 | 57.6 | 52.3 | 47.0 |
| **36** | SMOTEENN | 50 | 320 | 36.4 | 57.6 | 52.3 | 47.0 |
| **37** | SMOTEENN | 100 | 10 | 36.4 | 60.6 | 54.5 | 48.5 |
| **38** | SMOTEENN | 100 | 20 | 36.4 | 60.6 | 54.5 | 48.5 |
| **39** | SMOTEENN | 100 | 40 | 36.4 | 60.6 | 54.5 | 48.5 |
| **40** | SMOTEENN | 100 | 80 | 36.4 | 60.6 | 54.5 | 48.5 |
| **41** | SMOTEENN | 100 | 160 | 36.4 | 60.6 | 54.5 | 48.5 |
| **42** | SMOTEENN | 100 | 320 | 36.4 | 60.6 | 54.5 | 48.5 |
| **43** | SMOTEENN | 200 | 10 | 27.3 | 66.7 | 56.8 | 47.0 |
| **44** | SMOTEENN | 200 | 20 | 27.3 | 66.7 | 56.8 | 47.0 |
| **45** | SMOTEENN | 200 | 40 | 27.3 | 66.7 | 56.8 | 47.0 |
| **46** | SMOTEENN | 200 | 80 | 27.3 | 66.7 | 56.8 | 47.0 |
| **47** | SMOTEENN | 200 | 160 | 27.3 | 66.7 | 56.8 | 47.0 |
| **48** | SMOTEENN | 200 | 320 | 27.3 | 66.7 | 56.8 | 47.0 |
| **49** | RO | 10 | 10 | 36.4 | 81.8 | 70.5 | 59.1 |
| **50** | RO | 10 | 20 | 36.4 | 81.8 | 70.5 | 59.1 |
| **51** | RO | 10 | 40 | 36.4 | 81.8 | 70.5 | 59.1 |
| **52** | RO | 10 | 80 | 36.4 | 81.8 | 70.5 | 59.1 |
| **53** | RO | 10 | 160 | 36.4 | 81.8 | 70.5 | 59.1 |
| **54** | RO | 10 | 320 | 36.4 | 81.8 | 70.5 | 59.1 |
| **55** | RO | 50 | 10 | 45.4 | 78.7 | 70.5 | 62.1 |
| **56** | RO | 50 | 20 | 45.4 | 78.8 | 70.5 | 62.1 |
| **57** | RO | 50 | 40 | 45.4 | 78.8 | 70.5 | 62.1 |
| **58** | RO | 50 | 80 | 45.4 | 78.8 | 70.5 | 62.1 |
| **59** | RO | 50 | 160 | 45.4 | 78.8 | 70.5 | 62.1 |
| **60** | RO | 50 | 320 | 45.4 | 78.8 | 70.5 | 62.1 |
| **61** | RO | 100 | 10 | 45.4 | 78.8 | 70.5 | 62.1 |
| **62** | RO | 100 | 20 | 45.4 | 78.8 | 70.5 | 62.1 |
| **63** | RO | 100 | 40 | 45.4 | 78.8 | 70.5 | 62.1 |
| **64** | RO | 100 | 80 | 45.4 | 78.8 | 70.5 | 62.1 |
| **65** | RO | 100 | 160 | 45.4 | 78.8 | 70.5 | 62.1 |
| **66** | RO | 100 | 320 | 45.4 | 78.8 | 70.5 | 62.1 |
| **67** | RO | 200 | 10 | 36.4 | 78.8 | 68.2 | 57.6 |
| **68** | RO | 200 | 20 | 36.4 | 78.8 | 68.2 | 57.6 |
| **69** | RO | 200 | 40 | 36.4 | 78.8 | 68.2 | 57.6 |
| **70** | RO | 200 | 80 | 36.4 | 78.8 | 68.2 | 57.6 |
| **71** | RO | 200 | 160 | 36.4 | 78.8 | 68.2 | 57.6 |
| **72** | RO | 200 | 320 | 36.4 | 78.8 | 68.2 | 57.6 |
| **73** | None | 10 | 10 | 0.0 | 90.9 | 68.2 | 45.5 |
| **74** | None | 10 | 20 | 0.0 | 90.9 | 68.2 | 45.5 |
| **75** | None | 10 | 40 | 0.0 | 90.9 | 68.2 | 45.5 |
| **76** | None | 10 | 80 | 0.0 | 90.9 | 68.2 | 45.5 |
| **77** | None | 10 | 160 | 0.0 | 90.9 | 68.2 | 45.5 |
| **78** | None | 10 | 320 | 0.0 | 90.9 | 68.2 | 45.5 |
| **79** | None | 50 | 10 | 0.0 | 93.9 | 70.5 | 47.0 |
| **80** | None | 50 | 20 | 0.0 | 93.9 | 70.5 | 47.0 |
| **81** | None | 50 | 40 | 0.0 | 93.9 | 70.5 | 47.0 |
| **82** | None | 50 | 80 | 0.0 | 93.9 | 70.5 | 47.0 |
| **83** | None | 50 | 160 | 0.0 | 93.9 | 70.5 | 47.0 |
| **84** | None | 50 | 320 | 0.0 | 93.9 | 70.5 | 47.0 |
| **85** | None | 100 | 10 | 0.0 | 93.9 | 70.5 | 47.0 |
| **86** | None | 100 | 20 | 0.0 | 93.9 | 70.5 | 47.0 |
| **87** | None | 100 | 40 | 0.0 | 93.9 | 70.5 | 47.0 |
| **88** | None | 100 | 80 | 0.0 | 93.9 | 70.5 | 47.0 |
| **89** | None | 100 | 160 | 0.0 | 93.9 | 70.5 | 47.0 |
| **90** | None | 100 | 320 | 0.0 | 93.9 | 70.5 | 47.0 |
| **91** | None | 200 | 10 | 0.0 | 97.0 | 72.7 | 48.5 |
| **92** | None | 200 | 20 | 0.0 | 97.0 | 72.7 | 48.5 |
| **93** | None | 200 | 40 | 0.0 | 97.0 | 72.7 | 48.5 |
| **94** | None | 200 | 80 | 0.0 | 97.0 | 72.7 | 48.5 |
| **95** | None | 200 | 160 | 0.0 | 97.0 | 72.7 | 48.5 |
| **96** | None | 200 | 320 | 0.0 | 97.0 | 72.7 | 48.5 |

| Model ID | Oversampling | Trees (n) | Leaves (n) | Sensitivity (%) | Specificity (%) | Accuracy (%) | Macro average accuracy (%) |
|---|---|---|---|---|---|---|---|
| **1** | SMOTE | 10 | 10 | 47.1 | 78.8 | 68.0 | 62.9 |
| **2** | SMOTE | 10 | 20 | 47.1 | 78.8 | 68.0 | 62.9 |
| **3** | SMOTE | 10 | 40 | 47.1 | 78.8 | 68.0 | 62.9 |
| **4** | SMOTE | 10 | 80 | 47.1 | 78.8 | 68.0 | 62.9 |
| **5** | SMOTE | 10 | 160 | 47.1 | 78.8 | 68.0 | 62.9 |
| **6** | SMOTE | 10 | 320 | 47.1 | 78.8 | 68.0 | 62.9 |
| **7** | SMOTE | 50 | 10 | 58.8 | 81.8 | 74.0 | 70.3 |
| **8** | SMOTE | 50 | 20 | 58.8 | 81.8 | 74.0 | 70.3 |
| **9** | SMOTE | 50 | 40 | 58.8 | 81.8 | 74.0 | 70.3 |
| **10** | SMOTE | 50 | 80 | 58.8 | 81.8 | 74.0 | 70.3 |
| **11** | SMOTE | 50 | 160 | 58.8 | 81.8 | 74.0 | 70.3 |
| **12** | SMOTE | 50 | 320 | 58.8 | 81.8 | 74.0 | 70.3 |
| **13** | SMOTE | 100 | 10 | 64.7 | 81.8 | 76.0 | 73.3 |
| **14** | SMOTE | 100 | 20 | 64.7 | 81.8 | 76.0 | 73.3 |
| **15** | SMOTE | 100 | 40 | 64.7 | 81.8 | 76.0 | 73.3 |
| **16** | SMOTE | 100 | 80 | 64.7 | 81.8 | 76.0 | 73.3 |
| **17** | SMOTE | 100 | 160 | 64.7 | 81.8 | 76.0 | 73.3 |
| **18** | SMOTE | 100 | 320 | 64.7 | 81.8 | 76.0 | 73.3 |
| **19** | SMOTE | 200 | 10 | 58.8 | 84.8 | 76.0 | 71.8 |
| **20** | SMOTE | 200 | 20 | 58.8 | 84.8 | 76.0 | 71.8 |
| **21** | SMOTE | 200 | 40 | 58.8 | 84.8 | 76.0 | 71.8 |
| **22** | SMOTE | 200 | 80 | 58.8 | 84.8 | 76.0 | 71.8 |
| **23** | SMOTE | 200 | 160 | 58.8 | 84.8 | 76.0 | 71.8 |
| **24** | SMOTE | 200 | 320 | 58.8 | 84.8 | 76.0 | 71.8 |
| **25** | SMOTEENN | 10 | 10 | 58.8 | 69.7 | 66.0 | 64.3 |
| **26** | SMOTEENN | 10 | 20 | 58.8 | 69.7 | 66.0 | 64.3 |
| **27** | SMOTEENN | 10 | 40 | 58.8 | 69.7 | 66.0 | 64.3 |
| **28** | SMOTEENN | 10 | 80 | 58.8 | 69.7 | 66.0 | 64.3 |
| **29** | SMOTEENN | 10 | 160 | 58.8 | 69.7 | 66.0 | 64.3 |
| **30** | SMOTEENN | 10 | 320 | 58.8 | 69.7 | 66.0 | 64.3 |
| **31** | SMOTEENN | 50 | 10 | 76.5 | 69.7 | 72.0 | 73.1 |
| **32** | SMOTEENN | 50 | 20 | 76.5 | 69.7 | 72.0 | 73.1 |
| **33** | SMOTEENN | 50 | 40 | 76.5 | 69.7 | 72.0 | 73.1 |
| **34** | SMOTEENN | 50 | 80 | 76.5 | 69.7 | 72.0 | 73.1 |
| **35** | SMOTEENN | 50 | 160 | 76.5 | 69.7 | 72.0 | 73.1 |
| **36** | SMOTEENN | 50 | 320 | 76.5 | 69.7 | 72.0 | 73.1 |
| **37** | SMOTEENN | 100 | 10 | 82.3 | 72.7 | 76.0 | 77.5 |
| **38** | SMOTEENN | 100 | 20 | 82.3 | 72.7 | 76.0 | 77.5 |
| **39** | SMOTEENN | 100 | 40 | 82.3 | 72.7 | 76.0 | 77.5 |
| **40** | SMOTEENN | 100 | 80 | 82.3 | 72.7 | 76.0 | 77.5 |
| **41** | SMOTEENN | 100 | 160 | 82.3 | 72.7 | 76.0 | 77.5 |
| **42** | SMOTEENN | 100 | 320 | 82.3 | 72.7 | 76.0 | 77.5 |
| **43** | SMOTEENN | 200 | 10 | 82.3 | 72.7 | 76.0 | 77.5 |
| **44** | SMOTEENN | 200 | 20 | 82.3 | 72.7 | 76.0 | 77.5 |
| **45** | SMOTEENN | 200 | 40 | 82.3 | 72.7 | 76.0 | 77.5 |
| **46** | SMOTEENN | 200 | 80 | 82.3 | 72.7 | 76.0 | 77.5 |
| **47** | SMOTEENN | 200 | 160 | 82.3 | 72.7 | 76.0 | 77.5 |
| **48** | SMOTEENN | 200 | 320 | 82.3 | 72.7 | 76.0 | 77.5 |
| **49** | RO | 10 | 10 | 64.7 | 78.8 | 74.0 | 71.7 |
| **50** | RO | 10 | 20 | 64.7 | 78.8 | 74.0 | 71.7 |
| **51** | RO | 10 | 40 | 64.7 | 78.8 | 74.0 | 71.7 |
| **52** | RO | 10 | 80 | 64.7 | 78.8 | 74.0 | 71.7 |
| **53** | RO | 10 | 160 | 64.7 | 78.8 | 74.0 | 71.7 |
| **54** | RO | 10 | 320 | 64.7 | 78.8 | 74.0 | 71.7 |
| **55** | RO | 50 | 10 | 52.9 | 78.8 | 70.0 | 65.9 |
| **56** | RO | 50 | 20 | 52.9 | 78.8 | 70.0 | 65.9 |
| **57** | RO | 50 | 40 | 52.9 | 78.8 | 70.0 | 65.9 |
| **58** | RO | 50 | 80 | 52.9 | 78.8 | 70.0 | 65.9 |
| **59** | RO | 50 | 160 | 52.9 | 78.8 | 70.0 | 65.9 |
| **60** | RO | 50 | 320 | 52.9 | 78.8 | 70.0 | 65.9 |
| **61** | RO | 100 | 10 | 58.8 | 81.8 | 74.0 | 70.3 |
| **62** | RO | 100 | 20 | 58.8 | 81.8 | 74.0 | 70.3 |
| **63** | RO | 100 | 40 | 58.8 | 81.8 | 74.0 | 70.3 |
| **64** | RO | 100 | 80 | 58.8 | 81.8 | 74.0 | 70.3 |
| **65** | RO | 100 | 160 | 58.8 | 81.8 | 74.0 | 70.3 |
| **66** | RO | 100 | 320 | 58.8 | 81.8 | 74.0 | 70.3 |
| **67** | RO | 200 | 10 | 64.7 | 81.8 | 76.0 | 73.3 |
| **68** | RO | 200 | 20 | 64.7 | 81.8 | 76.0 | 73.3 |
| **69** | RO | 200 | 40 | 64.7 | 81.8 | 76.0 | 73.3 |
| **70** | RO | 200 | 80 | 64.7 | 81.8 | 76.0 | 73.3 |
| **71** | RO | 200 | 160 | 64.7 | 81.8 | 76.0 | 73.3 |
| **72** | RO | 200 | 320 | 64.7 | 81.8 | 76.0 | 73.3 |
| **73** | None | 10 | 10 | 35.3 | 84.8 | 68.0 | 60.1 |
| **74** | None | 10 | 20 | 35.3 | 84.8 | 68.0 | 60.1 |
| **75** | None | 10 | 40 | 35.3 | 84.8 | 68.0 | 60.1 |
| **76** | None | 10 | 80 | 35.3 | 84.8 | 68.0 | 60.1 |
| **77** | None | 10 | 160 | 35.3 | 84.8 | 68.0 | 60.1 |
| **78** | None | 10 | 320 | 35.3 | 84.8 | 68.0 | 60.1 |
| **79** | None | 50 | 10 | 23.5 | 90.9 | 68.0 | 57.2 |
| **80** | None | 50 | 20 | 23.5 | 90.9 | 68.0 | 57.2 |
| **81** | None | 50 | 40 | 23.5 | 90.9 | 68.0 | 57.2 |
| **82** | None | 50 | 80 | 23.5 | 90.9 | 68.0 | 57.2 |
| **83** | None | 50 | 160 | 23.5 | 90.9 | 68.0 | 57.2 |
| **84** | None | 50 | 320 | 23.5 | 90.9 | 68.0 | 57.2 |
| **85** | None | 100 | 10 | 29.4 | 87.9 | 68.0 | 58.6 |
| **86** | None | 100 | 20 | 29.4 | 87.9 | 68.0 | 58.6 |
| **87** | None | 100 | 40 | 29.4 | 87.9 | 68.0 | 58.6 |
| **88** | None | 100 | 80 | 29.4 | 87.9 | 68.0 | 58.6 |
| **89** | None | 100 | 160 | 29.4 | 87.9 | 68.0 | 58.6 |
| **90** | None | 100 | 320 | 29.4 | 87.9 | 68.0 | 58.6 |
| **91** | None | 200 | 10 | 17.6 | 87.9 | 64.0 | 52.8 |
| **92** | None | 200 | 20 | 17.6 | 87.9 | 64.0 | 52.8 |
| **93** | None | 200 | 40 | 17.6 | 87.9 | 64.0 | 52.8 |
| **94** | None | 200 | 80 | 17.6 | 87.9 | 64.0 | 52.8 |
| **95** | None | 200 | 160 | 17.6 | 87.9 | 64.0 | 52.8 |
| **96** | None | 200 | 320 | 17.6 | 87.9 | 64.0 | 52.8 |
